# Metabolite genome-wide association study (mGWAS) and gene-metabolite interaction network analysis reveal potential biomarkers for feed efficiency in pigs

**DOI:** 10.1101/2020.04.11.036939

**Authors:** Xiao Wang, Haja N. Kadarmideen

## Abstract

Metabolites represent the ultimate response of biological systems, so metabolomics is considered to link the genotypes and phenotypes. Feed efficiency is one of the most important phenotypes in sustainable pig production and is the main breeding goal trait. We utilized metabolic and genomic datasets from a total of 108 pigs from our own previously published studies that involved 59 Duroc and 49 Landrace pigs with data on feed efficiency (residual feed intake or RFI), genotype (PorcineSNP80 BeadChip) data and metabolomic data (45 final metabolite datasets derived from LC-MS system). Utilizing these datasets, our main aim was to identify genetic variants (single-nucleotide polymorphisms or SNPs) that affect 45 different metabolite concentrations in plasma collected at the start and end of the performance testing of pigs categorized as high or low in their feed efficiency (based on RFI values). Genome-wide significant genetic variants could be then used as potential genetic or biomarkers in breeding programs for feed efficiency. The other objective was to reveal the biochemical mechanisms underlying genetic variations for pigs’ feed efficiency. In order to achieve these objectives, we firstly conducted a metabolite genome-wide association study (mGWAS) based on mixed linear models and found 152 genome-wide significant SNPs (*P*-value < 1.06E-06) in association with 17 metabolites that included 90 significant SNPs annotated to 52 genes. On chromosome one alone, 51 significant SNPs associated with isovalerylcarnitine and propionylcarnitine were found to be in strong linkage disequilibrium (LD). SNPs in strong LD annotated to *FBXL4* and *CCNC* consisted of two haplotype blocks where three SNPs (ALGA0004000, ALGA0004041 and ALGA0004042) were in the intron regions of *FBXL4* and *CCNC*. The interaction network revealed that *CCNC* and *FBXL4* were linked by the hub gene *N6AMT1* that was associated with isovalerylcarnitine and propionylcarnitine. Moreover, three metabolites (i.e., isovalerylcarnitine, propionylcarnitine and pyruvic acid) were clustered in one group based on the low-high RFI pigs.

This study performed a comprehensive metabolite-based GWAS analysis for pigs with differences in feed efficiency and provided significant metabolites for which there is a significant genetic variation as well as biological interaction networks. The identified metabolite genetic variants, genes and networks in high versus low feed efficient pigs could be considered as potential genetic or biomarkers for feed efficiency.

## 1. Introduction

Large populations are generally essential for genome-wide association study (GWAS) to obtain sufficient statistical power for the identification of genetic polymorphisms [1]. However, some intermediate phenotypes like metabolites could potentially avoid this problem, as they are directly involved in metabolite conversion modification [2,3]. As the end products of cellular regulatory processes, metabolites represent the ultimate response of biological systems associated with genetic changes, so metabolomics is considered to link the genotypes and phenotypes [4]. Metabolomics refers to the measurements of all endogenous metabolites, intermediates and products of metabolism and has been applied to measure the dynamic metabolic responses in pigs [5,6] and dairy cows [7,8]. Additionally, metabolites could provide details of physiological state, so genetic variants associated metabolites are expected to display larger effect sizes [9]. Gieger et al. (2008) firstly used metabolite concentrations as quantitative traits in association with genotypes and found their available applications in GWAS [9]. Do et al. [10] conducted GWAS using residual feed intake (RFI) phenotypes to identify single-nucleotide polymorphisms (SNPs) that explain significant variations in feed efficiency for pigs. Our previous study found two metabolites (i.e., α-ketoglutarate and succinic acid) in a RFI-related network of dairy cows, which could represent biochemical mechanisms underlying variation for phenotypes of feed efficiency [8].

In this study, we aimed to identify genetic variants (SNP markers) affecting concentrations of metabolites and to reveal the biochemical mechanisms underlying genetic variations for pigs’ feed efficiency. Our study is based on two of our previously published papers and datasets used therein [6,11]. Briefly, the experiment consisted of 59 Duroc and 49 Landrace pigs with data on feed efficiency (RFI), genotype (PorcineSNP80 BeadChip) data and metabolomic data (45 final metabolite datasets derived from liquid chromatography-mass spectrometry (LC-MS) system). While our previous studies only looked at metabolome-phenotype associations, we report an integrated systems genomics approach to identify quantitative trait loci (QTLs) or SNPs affecting metabolite concentration via metabolite GWAS methods (mGWAS), where each metabolite is itself a phenotype. To the best of our knowledge, this is the first study to link the genomics with metabolomics to identify significant genetic variants associated with known metabolites that differ in pigs with different levels of feed efficiency. Main aims of our study are as follows:

1. Find significant SNP markers associated with all the metabolites in the metabolomics dataset using mGWAS method, and then to reveal the biochemical mechanisms underlying genetic variations for porcine feed efficiency using 108 Danish pigs in low and high RFI conditions, genotyped by 68K PorcineSNP80 BeadChip array.
2. Annotate identified significant SNP markers to porcine genes.
3. Annotate metabolites and identify enriched metabolic pathways and gene-metabolite networks to find the potential biomarkers that were strongly associated with feed efficiency.

## 2. Results

### 2.1. First component score and significant metabolic pathways of 45 metabolites

The partial least squares-discriminant analysis (PLS-DA) results revealed that the first component score (Component 1) explained more than 75% variations of all 45 metabolites. In addition, the Duroc and Landrace pigs were clearly stratified, especially using the metabolite values between Duroc from first sampling time and Landrace from second sampling time (Figure 1A). The most significant metabolic pathways were the aminoacyl-tRNA biosynthesis, following by the arginine biosynthesis, the arginine and proline metabolism, the alanine, aspartate and glutamate metabolism (Figure 1B). As the pathway impact of the aminoacyl-tRNA biosynthesis was zero, we discarded this significant pathway and only used the metabolites enriched in the other three significant pathways for GWAS (Table 1). Thus, the metabolite means for 5 compounds in the arginine biosynthesis (arginine, aspartic acid, citrulline, glutamic acid, ornithine), 5 compounds in the arginine and proline metabolism (arginine, glutamic acid, ornithine, proline, pyruvic acid) and 4 compounds in the alanine, aspartate and glutamate metabolism (alanine, aspartic acid, glutamic acid, pyruvic acid) metabolites were calculated and shown in Table 1.

**Table 1.** Significant metabolic pathways (FDR < 0.1) using *Homo sapiens* as the library.

| Metabolic pathway | Match Status | Involved metabolites | *P*-value | $-\log(P\text{-value})$ | False discovery rate (FDR) | Pathway impact value |
| --- | --- | --- | --- | --- | --- | --- |
| Aminoacyl-tRNA biosynthesis (ssc00970) | 9/48 | Alanine, Arginine, Aspartic acid, Glutamic acid, Isoleucine, Methionine, Phenylalanine, Proline, Threonine | 3.55E-07 | 14.85 | 2.98E-5 | 0 |
| Arginine biosynthesis (ssc00220) | 5/14 | Arginine, Aspartic acid, Citrulline, Glutamic acid, Ornithine | 6.53E-06 | 11.94 | 2.74E-4 | 0.48 |
| Arginine and proline metabolism (ssc00330) | 5/38 | Arginine, Glutamic acid, Ornithine, Proline, Pyruvic acid | 1.12E-03 | 6.79 | 0.031 | 0.33 |
| Alanine, aspartate and glutamate metabolism (ssc00250) | 4/28 | Alanine, Aspartic acid, Glutamic acid, Pyruvic acid | 2.72E-03 | 5.91 | 0.057 | 0.42 |

**Figure 1.**
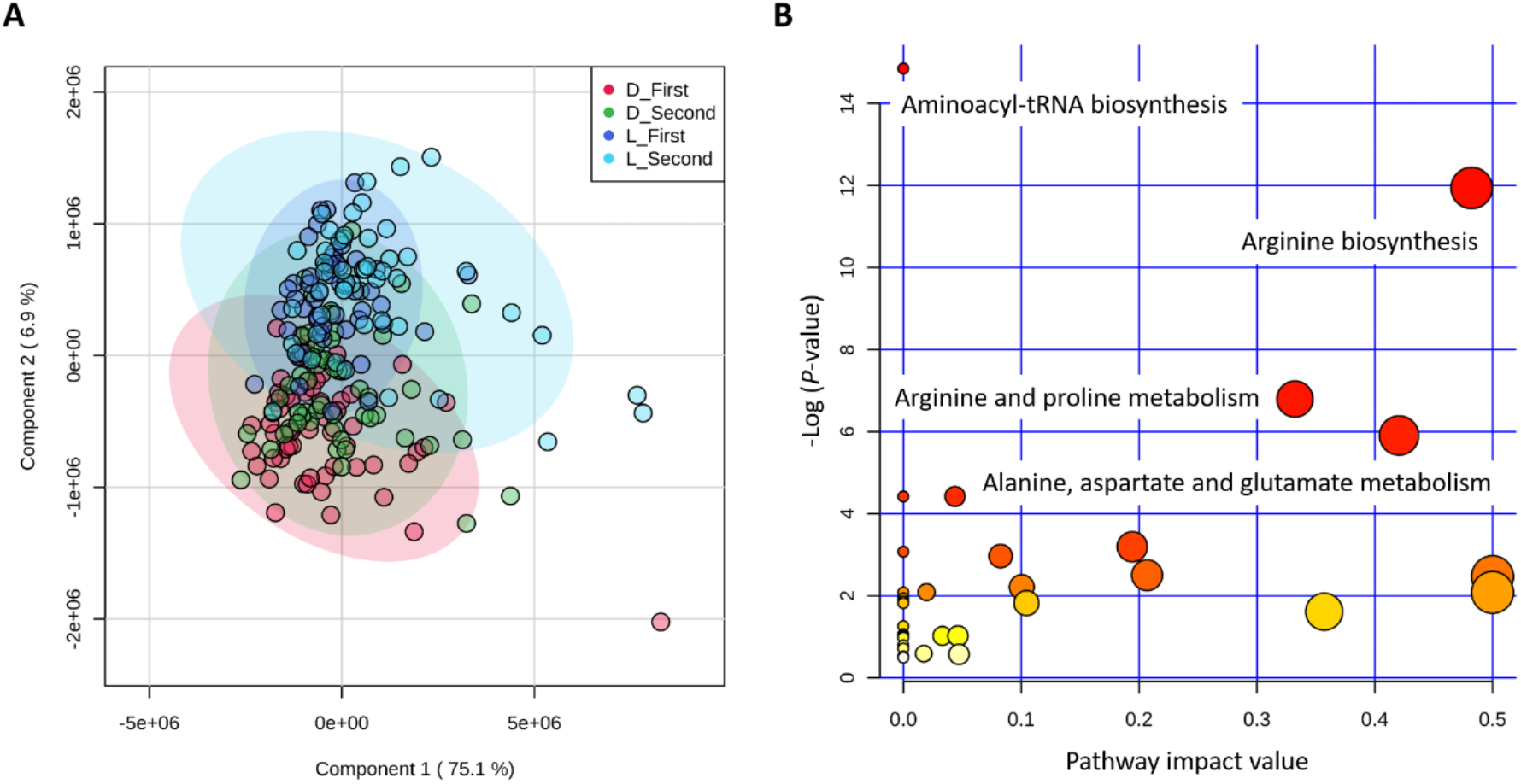
(**A**) Partial least squares-discriminant analysis (PLS-DA) of 45 metabolites. Note: D/L with First/Second indicates the sampling time of Duroc/Landrace pigs. (**B**) Metabolic pathways for 45 metabolites using *Homo sapiens* as the library. Note: The size and color of the circles for each pathway indicate the matched metabolite ratio and the −log (*P*-value), respectively.

### 2.2. Genome-wide significant SNPs and gene annotation

Metabolite based GWAS for first, second and combined two sampling times revealed 152 genome-wide significant SNPs (Supplementary table 1) in association with 17 metabolites (Supplementary table 2). Unfortunately, no significant SNP was detected in association with first component scores (*P*-values ≥ 2.78E-06) and metabolites enriched in the significant metabolic pathways (*P*-values ≥ 1.74E-04), thus, GWAS results of the two scenarios were not listed. Manhattan plots of genome-wide association for isovalerylcarnitine and propionylcarnitine are shown in the Figure 2 and Manhattan plots for the other 43 metabolites are shown in the Supplementary figure 1. Along the whole genome, significant SNPs associated with isovalerylcarnitine and propionylcarnitine from the second sampling time were mainly located on the chromosome one (Figure 2). The overlapped significant SNPs associated with more than two different metabolites were shown in Table 2, where 57 significant SNPs on genome level were associated with isovalerylcarnitine and propionylcarnitine from the second sampling time. In addition, another 3 metabolites (1-hexadecyl-sn-glycero-3-phosphocholine, 1-myristoyl-sn-glycero-3-phosphocholine, lysoPC(16:0)) were also significantly associated with 10 SNPs (Table 2).

**Table 2.**
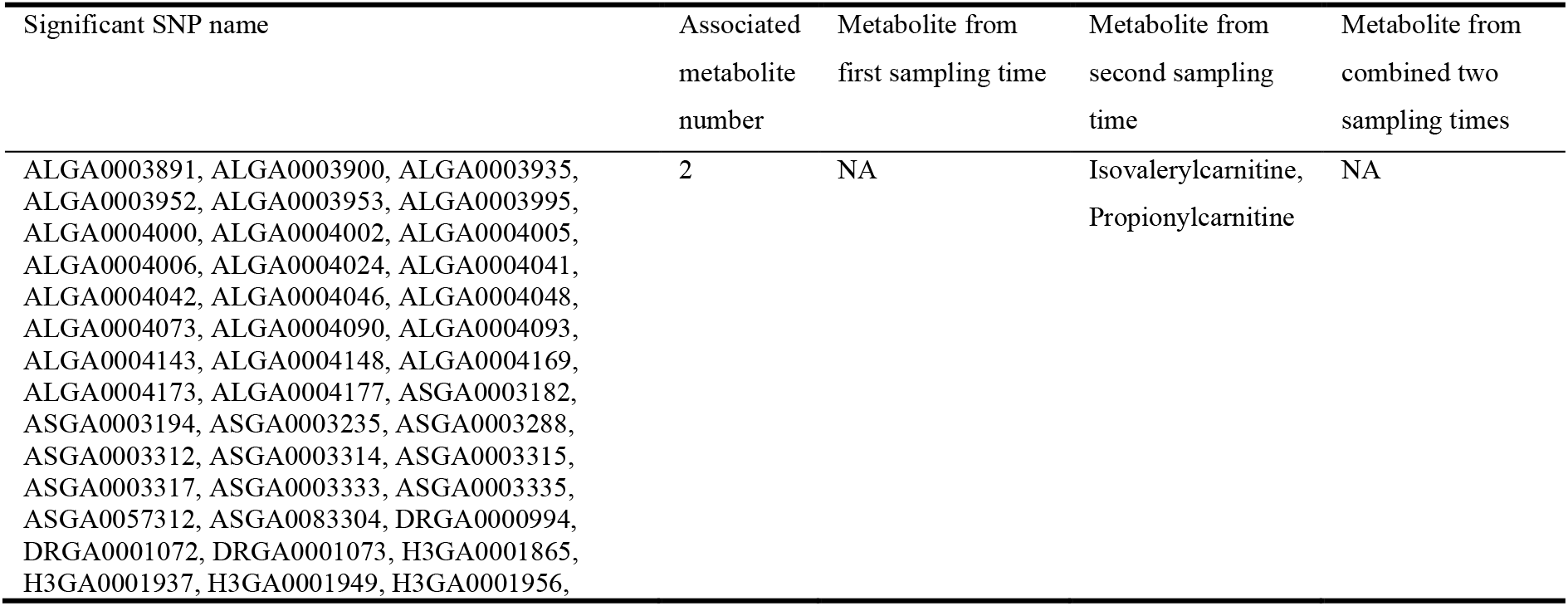
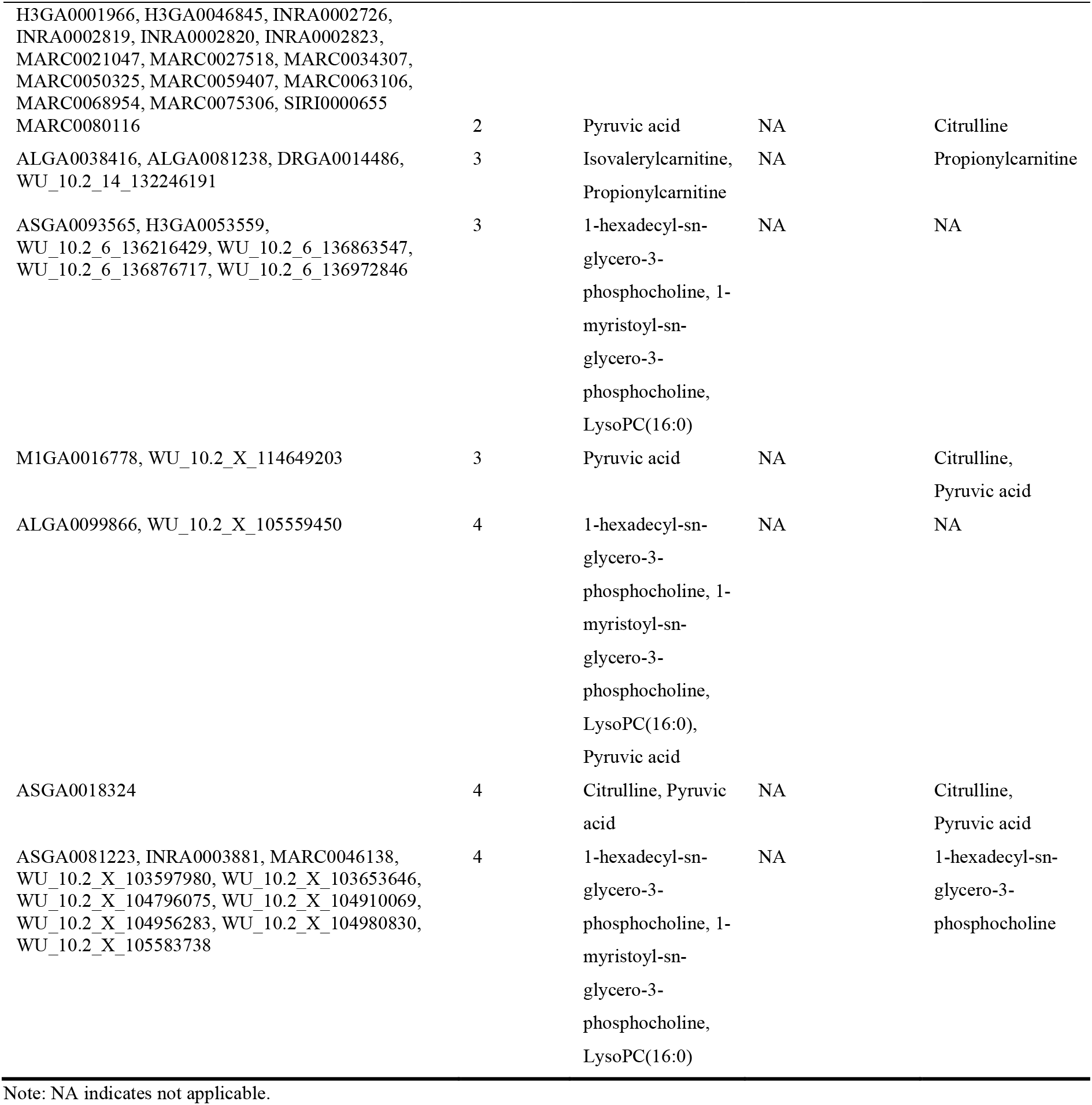
Common significant SNPs of genome-wide association for more than two different metabolites from first, second and combined two sampling times.

**Figure 2.**
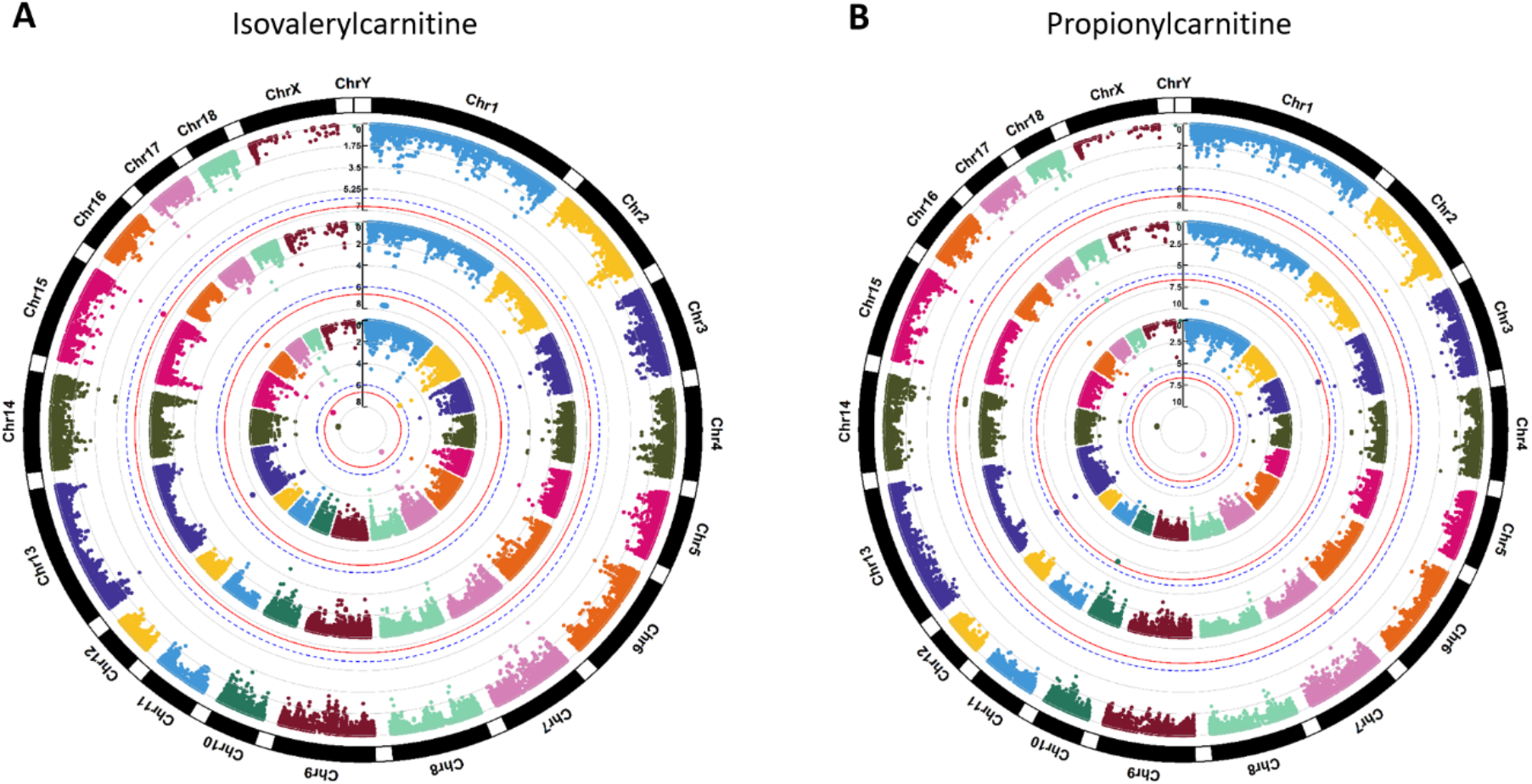
Manhattan plots of genome-wide association for (**A**) isovalerylcarnitine and (**B**) propionylcarnitine. Note: Y-axis indicates the log_10_(*P*-value). Blue dotted and red solid lines indicate the genome-wide threshold of 0.05 and 0.01 after Bonferroni multiple testing, respectively. The three tracks indicate the metabolites from first sampling time, second sampling time and combined two sampling times, respectively, from outside to inside.

After annotation of significant SNPs to the neighboring genes and gene components (Sscrofa10.2/susScr3), we found that 90 significant SNPs were within a 500kb window of 52 neighboring genes (Supplementary table 1), and 6 significant SNPs were directly located in the gene components of 5 genes (Table 3). For example, if we only consider the SNPs on chromosome one, we found 29 significant SNPs near 9 genes (Supplementary table 1), whereas ALGA0004000, ALGA0004041 and ALGA0004042 were located in the introns of *FBXL4* and *CCNC*, respectively (Table 3). These results show that these genes may be involved in regulating abundance of the metabolites that are significantly different between high and low RFI pigs.

**Table 3.**
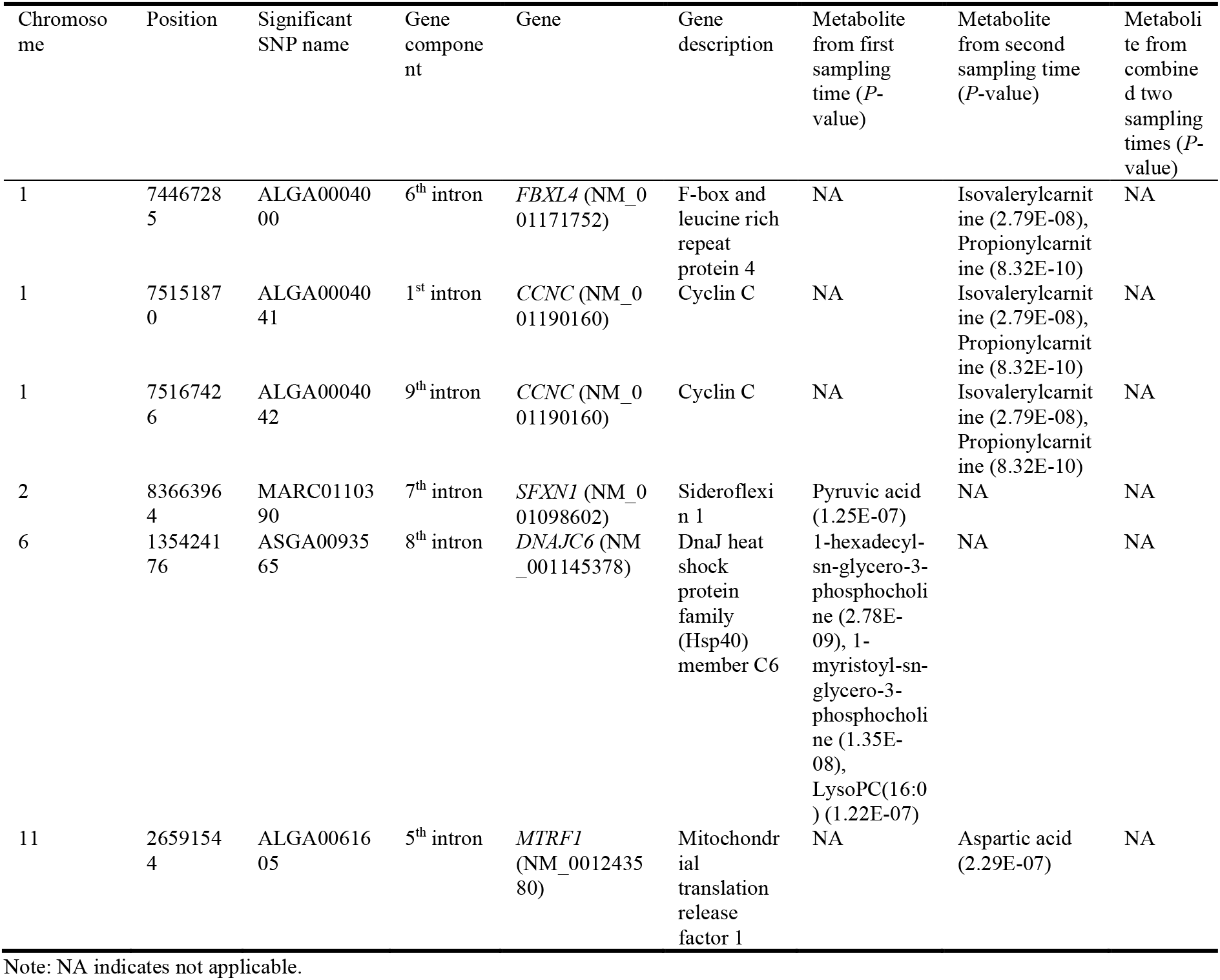
Gene component annotation for genome-wide significant SNPs.

The linkage disequilibrium (LD) pattern for all significant SNPs is shown in the Supplementary figure From the LD results for 58 significant SNPs on chromosome one, we found that 51 significant SNPs associated with isovalerylcarnitine (*P*-value = 2.79E-08) and propionylcarnitine (*P*-value = 8.32E-10) from second sampling time were in strong LD (Figure 3). Among the 58 significant SNPs, five of them were not in LD with the other 53 significant SNPs (Supplementary figure 2), so they were excluded in the haplotype visualization in the Figure 3. In detail, SNPs annotated to *LOC780435* (NM_001078684), *FHL5* (NM_001243314), *FBXL4* (NM_001171752), *CCNC* (NM_001190160)/*MCHR2* (NM_001044609), *SIM1* (NM_001172585) were in block 2, block 4, block 6, block 8, block 9/10, respectively. Furthermore, ALGA0004000 in the 6^th^ intron of *FBXL4* was in LD of block 6, together with another five SNPs (INRA0002726, MARC0075306, ALGA0003995, ALGA0004002 and ALGA0004005) that were located in the intergenic regions of *FBXL4*. Especially, three SNPs in strong LD consisted of block 8 that two SNPs (ALGA0004041 and ALGA0004042) were located in the second and ninth intron of *CCNC* (Figure 3, Table 3 and Supplementary table 1). The number of significant SNPs in strong LDs of the other chromosomes was less than the significant SNP number on chromosome one (Supplementary figure 2). Notably, MARC0110390 in the 7^th^ intron region of *SFXN1* (NM_001098602) on chromosome two, and ALGA0061605 in the 5^th^ intron region of *MTRF1* (NM_001243580) on chromosome eleven were not in the LD with other SNPs. However, ASGA0093565 in the 8^th^ intron region of *DNAJC6* (NM_001145378) was in strong LD with WU_10.2_6_135312468 that was annotated to *LEPROT* (NM_001145388) (Supplementary table 1 and Supplementary figure 2).

**Figure 3.**
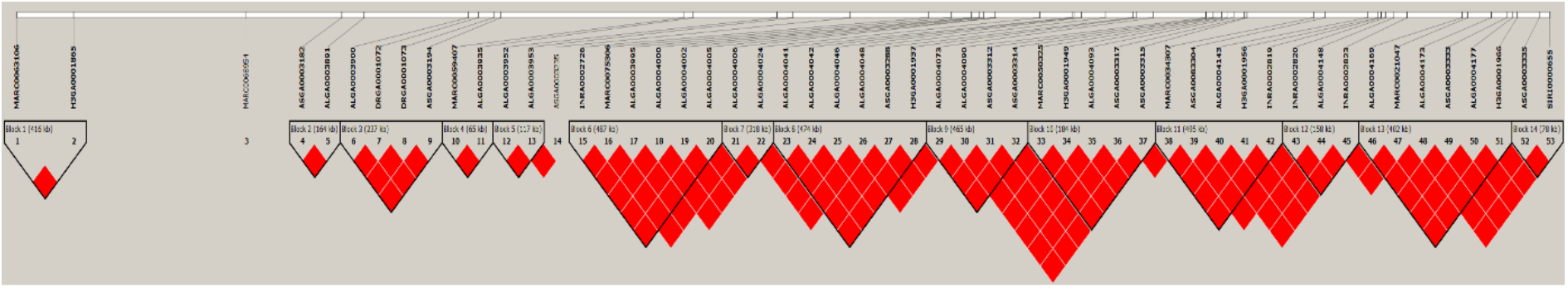
Linkage disequilibrium (LD) pattern for 53 significant SNPs on chromosome one. Note: Solid line triangle indicates LD. One square indicates LD level (r^2^) between two SNPs and the squares are colored by D’/LLOR standard scheme. D’/LLOR standard scheme is that red indicates LLOR > 2, D’ = 1; pink indicates LLOR > 2, D’ < 1; blue indicates LLOR < 2, D’ = 1; white indicates LLOR < 2, D’ < 1. LLOR is the logarithm of likelihood odds ratio and the reliable index to measure D’.

### 2.3. Gene and metabolite interaction network

The most significant enriched gene-based pathways were the human papillomavirus infection (ssc05165) with five genes (i.e., *CCND2*, *CTNNB1*, *JAK1*, *LAMC1* and *NFKB1*), followed by the PI3K-Akt signaling pathway (ssc04151) with five genes (i.e., *CCND2*, *F2R*, *JAK1*, *LAMC1* and *NFKB1*) and the hepatitis C (ssc05160) with four genes (i.e., *CLDN8*, *CTNNB1*, *JAK1* and *NFKB1*) (Figure 4A). Based on the gene-gene interaction network analysis, *CCNC* was in the good connections with *CDK8*, *CDK3* and *N6AMT1*, whereas *N6AMT1* linked to *FBXL4* (Figure 4B). Unfortunately, no gene-metabolite interaction network was found in this study. After the clustering of the SNP-related gene component associated metabolites (Table 3), we found that aspartic acid, 1-hexadecyl-sn-glycero-3-phosphocholine, 1-myristoyl-sn-glycero-3-phosphocholine and lysoPC(16:0) were clustered in the lower cluster, while the upper cluster included the metabolites of isovalerylcarnitine, propionylcarnitine and pyruvic acid (Figure 4C). Results show that metabolites from Duroc pigs have higher values in the upper cluster than those from lower cluster, but the metabolite values of Landrace pigs are higher in the lower cluster (Figure 4C). Afterwards, we investigated the metabolite values of aspartic acid, isovalerylcarnitine, propionylcarnitine and pyruvic acid that the associated significant SNPs were in the introns of *MTRF1*, *FBXL4*/*CCNC*, *SFXN1* (Table 3). Generally, propionylcarnitine from low RFI group had higher values while other three metabolite values in high RFI group seemed higher, but they are not significantly different between low and high RFI groups (*P*-value > 0.05) (Figure 4D).

**Figure 4.**
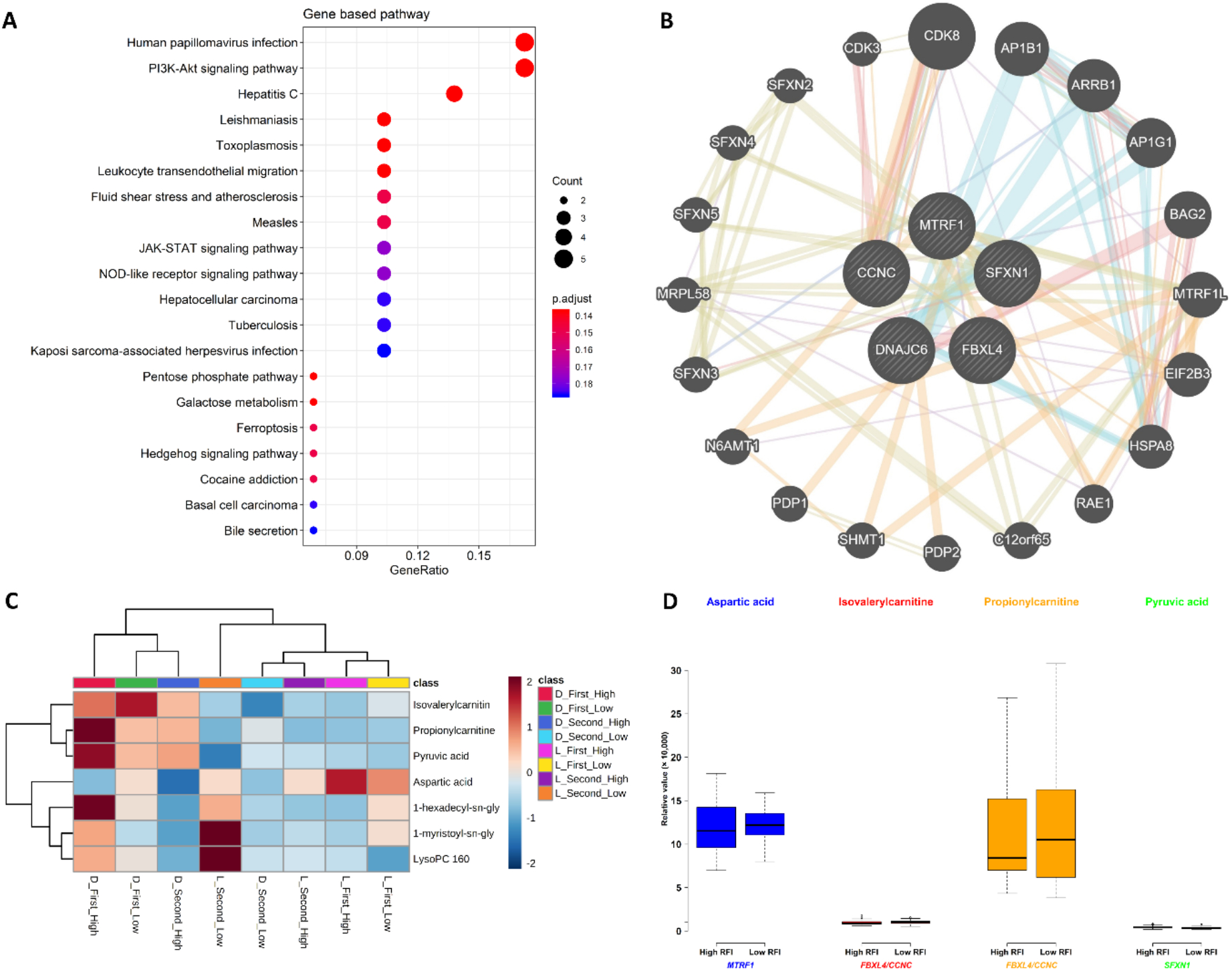
Gene pathway, metabolite cluster and the interaction network. (**A**) Pathway for significant SNP-related genes. (**B**) Network for the genes in which significant SNPs were annotate to gene components. (**C**) Heatmap cluster for the metabolites that were associated with significant SNPs annotated to gene components. (**D**) Metabolites (i.e., aspartic acid, isovalerylcarnitine, propionylcarnitine and pyruvic acid) values in high and low RFI pigs associated with the genes in which significant SNPs annotate to gene components. Note: The high RFI pigs and low RFI pigs were from left and right parts of all the pigs (n = 108) with one SD of actual RFI values.

## 3. Discussion

### 3.1 Metabolites in the PLS-DA and metabolic pathways of pigs

As breed types contributed to the RFI variations, RFI-related metabolomics could be breed-specific. Therefore, different breeds tend to exhibit different metabolite abundance values for example studies involving in the colostrum of pigs [25,26], in the milk and plasma of cattle [8,27], in the yolk and albumen of chickens [28,29], in the plasma of dogs [30] and in the fruit metabolite content of tomatoes [31]. In pigs, the heritability and genetic correlation of production traits of Duroc, Landrace and Yorkshire pigs vary. Duroc pigs showed lower heritability of feed efficiency but greater performance of growth traits [32,33]. The metabolomics of this study showed that metabolite values varies between two pig breeds and between the sampling times (Figure 1A and Figure 4C); hence, the breed differences between Duroc and Landrace pigs were driven both by genetic and metabolic factors.

The arginine biosynthesis pathway (ssc00220), where arginine, aspartic acid, citrulline, glutamic acid and ornithine were significantly enriched in our study (Table 1), plays a crucial role in amino acid metabolism, particularly in the synthesis of citrulline and proline in pigs [34]. By linking arginine, glutamate and proline in a bidirectional way, the arginine and proline metabolism pathway (ssc00330) biosynthesizes arginine and proline by glutamate. It is observed that proline metabolism associated with metastasis formation of breast cancer [35]. In the dairy cattle, the alanine, aspartate and glutamate metabolism (ssc00250) identified in the gene-based pathways of our study (Table 1) is the potential metabolic biomarker between the low and high feed efficient conditions [8].

### 3.2 Genome-wide significant SNP-related genes associated with metabolites

The previous GWAS for Duroc pigs identified two pleiotropic QTLs on chromosome one and seven for feed efficiency [33]. Do et al.(2014) [10] revealed 19 significant SNPs located on several chromosomes (e.g., one, three, seven, eight, nine, ten, fourteen and fifteen) that were highly associated with feed efficiency in Yorkshire pigs. In addition, other studies also found significant SNPs associated with RFI on other chromosomes, for example SNPs on chromosome five in the growing Piétrain–Large White pigs [36], on chromosome two in a crossed populations [37], on chromosome six in Large White pigs [38], etc. [39,40].

In this study, significant SNPs were mainly located on chromosome one (58/152), but the associated metabolites only mapped to 1-hexadecyl-sn-glycero-3-phosphocholine (1.7%), 1-myristoyl-sn-glycero-3-phosphocholine (1.7%), isovalerylcarnitine (47.0%), isoleucyl proline (0.9%), propionylcarnitine (47.0%) and lysoPC(16:0) (1.7%). Obviously, isovalerylcarnitine and propionylcarnitine primarily derived from amino acid catabolism were the major metabolites that associated with nine significant SNP-related genes (i.e., *CCNC*, *FBXL4*, *FHL5*, *LOC780435*, *MAT2B*, *MCHR2*, *PNISR*, *RRAGD* and *SIM1*) on chromosome one (Supplementary table 1). Previous study indicated that the amount of isovalerylcarnitine could decrease in the plasma and liver tissues but greatly increased in the muscle tissue, as a byproduct of leucine catabolism [41]. The isovalerylcarnitine compound was reported to be found in high amounts in the colostrum and milk of sows [42]. As a key role in the mitochondrial fatty acid transport and high-energy phosphate exchange, propionylcarnitine could improve cardiac dysfunction by reducing myocardial ischaemia [43].

### 3.3 Gene and metabolite interaction network

Based on the gene interaction node *N6AMT1*, one gene-gene interaction was found to connect *CCNC* with *FBXL4* (Figure 4B), in which significant SNPs were annotated to gene components and associated with isovalerylcarnitine and propionylcarnitine (Table 3). As the members of *CDK8* mediator complex that can regulate β-catenin-driven transcription, *CCNC* encodes the cell cycle regulatory protein cyclin C; in addition, results in the protein dysfunction due to the mutations of *CCNC* [44,45]. *CCNC* is also believed to increase the quiescent cells to maintain *CD34* expression after knocking down *CCNC* expression in human cord blood [46], while the amplification of *CCNC* was in an relationship with the unfavorable prognosis [47]. *FBXL4* is considered to participate in oxidative phosphorylation, mitochondrial dynamics, cell migration, prostate cancer metastasis, circadian GABAergic cyclic alteration, etc. [48–52]. The association results in pigs found that blood and immune traits were associated with the SNPs of *FBXL4* [53]. The node *N6AMT1* is responsible for DNA 6mA methylation modification, as the first glutamine-specific protein methyltransferase characterized in mammals, thus, glutamine could be regulated by the genes to promote the porcine growth performance [54,55].

## 4. Materials and methods

### 4.1. Animals, metabolites and genotypes

A total of 108 pigs were involved in this study including 59 Duroc and 49 Landrace pigs that were part of our own previous published studies [6,11]. The detailed description of the animal experiment and phenotype, metabolite and genotypes data collection are available from our previously published studies and all data used in this study were derived from our datasets that were made already public. Metabolite data [6] were accessed using MetaboLights accession ID MTBLS1384 with a link: https://www.ebi.ac.uk/metabolights/MTBLS1384. Genotype data [11] were accessed from National Center for Biotechnology Information (NCBI) GEO Accession Number: GSE144064 with the link: https://www.ncbi.nlm.nih.gov/geo/query/acc.cgi?acc=GSE144064.

As in Carmelo et al. [6], all the pigs were derived from sixteen-sire families in four generations and had RFI values calculated for each pig as the difference between the observed daily feed intake and the predicted feed intake. The range of actual RFI values of Duroc were from −10 to 14, while Landrace’s RFI value range was from −14 to 17 (Figure 5). In this study, we selectively chose the extreme left and extreme right tails of distribution of feed efficiency (RFI) distribution of all the pigs (n = 108) with one standard deviation (SD) from the mean [12] of actual RFI values. Then they were defined as high RFI pigs (RFI ≤ −5.23, n = 11) and low RFI pigs (RFI ≥ 5.23, n = 16), respectively (Figure 5). The overview of the analysis workflow is shown in Figure 6, and included five scenarios of phenotypes in the GWAS analysis based on different transformations of metabolites. The five types of phenotypes were (1) the metabolites from first sampling time, (2) the metabolites from second sampling time, (3) the metabolites from combined two sampling times, (4) the first component score (Component 1) from partial least squares-discriminant analysis (PLS-DA), and (5) the metabolites enriched in the significant metabolic pathways (Figure 6).

**Figure 5.**
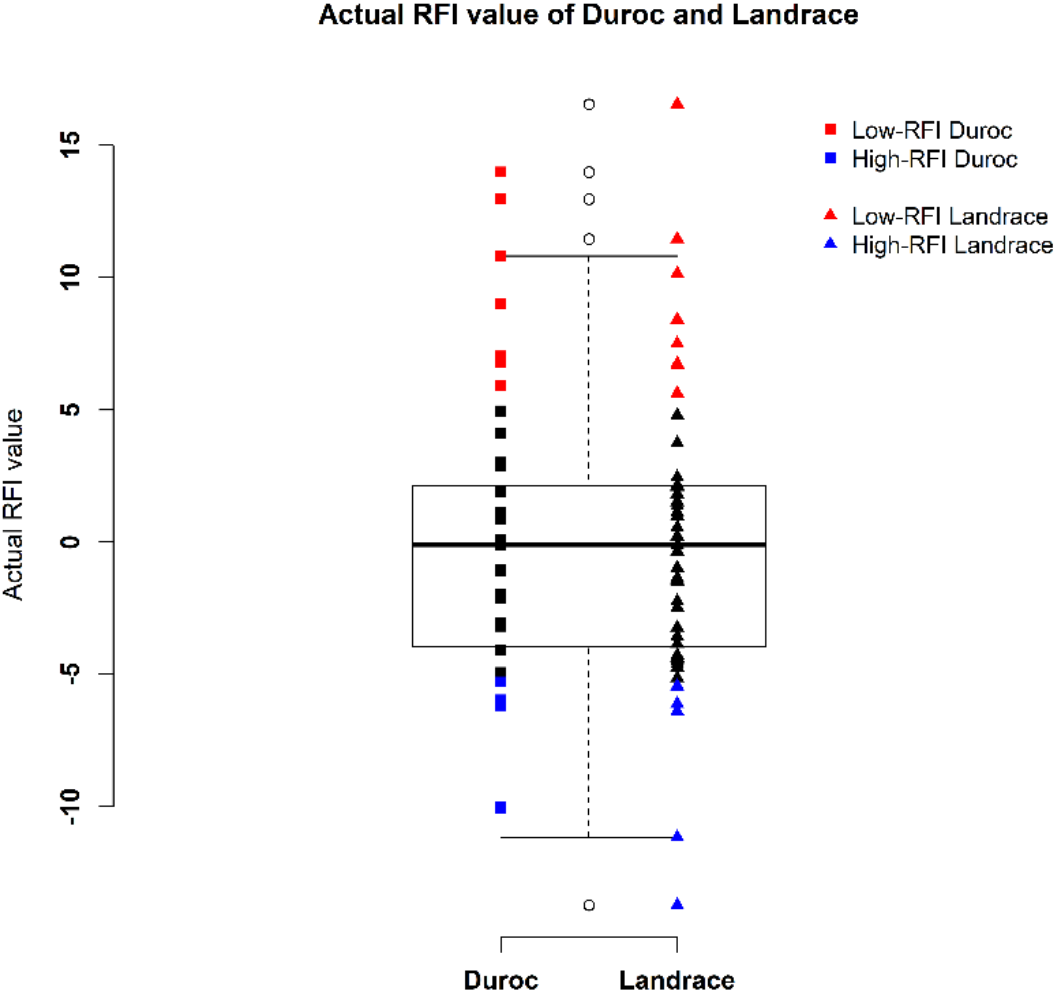
Distribution of actual RFI values of Duroc (n=59) and Landrace (n=49) pigs.

**Figure 6.**
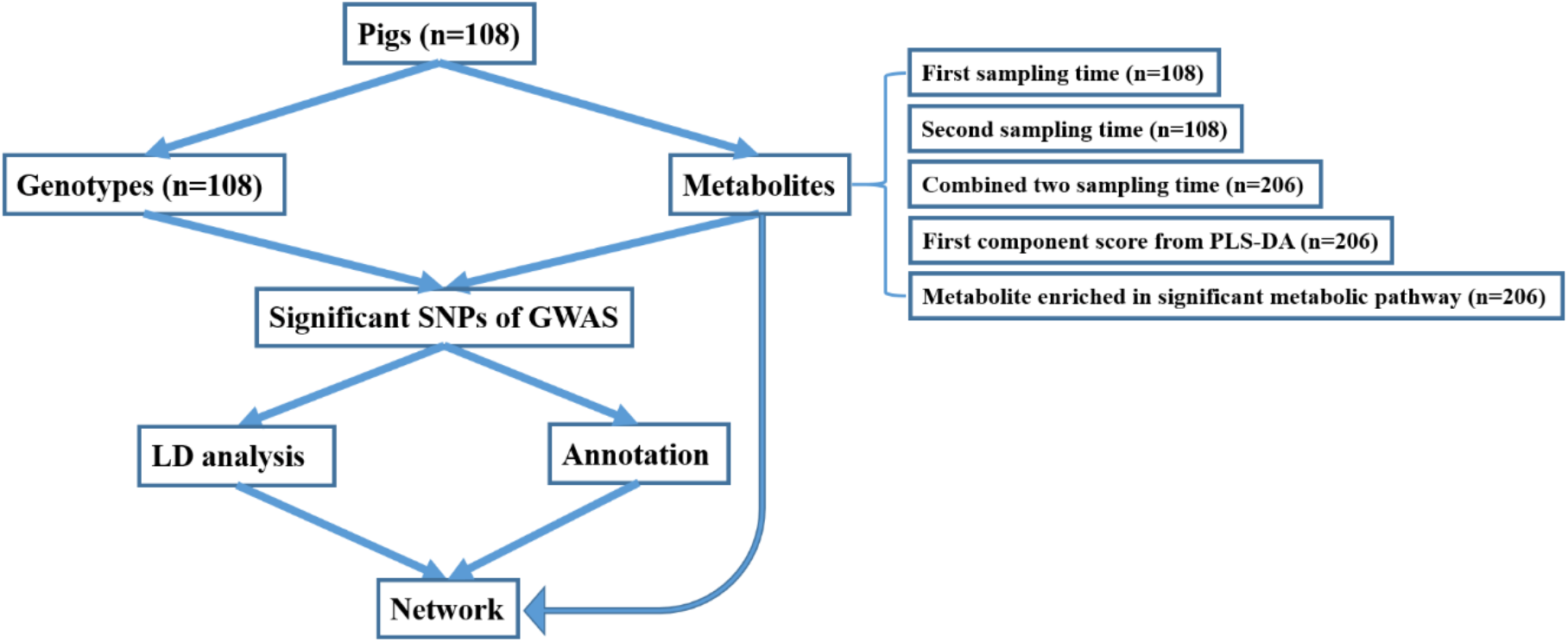
Overall analysis workflow.

Metabolite data was downloaded by accessing MetaboLights accession ID MTBLS1384 with a link: https://www.ebi.ac.uk/metabolights/MTBLS1384 that were collected in two sampling times for each pig by the previous study [6]. Finally, 45 metabolites were used in this study (Figure 7) including 16 annotated metabolites (i.e., 1-hexadecyl-sn-glycero-3-phosphocholine, 1-myristoyl-sn-glycero-3-phosphocholine, (3-Carboxypropyl)trimethylammonium, 5-methyl-5,6-dihydrouracils, acetaminophen, acetylcarnitine, benzoic acid, cotinine, creatinine, indoleacrylic acid, isoleucyl proline, isovalerylcarnitine, leucyl methionine, lysoPC(16:0), manNAc, propionylcarnitine) and 29 identified metabolites (i.e., 4-aminobenzoic acid, alanine, arginine, aspartic acid, carnitine, citrulline, cytidine, disaccharide, glutamic acid, guanine, guanosine, hypoxanthine, inosine, isoleucine, lactic acid, methionine, monosaccharide, nicotine amide, ornithine, phenylalanine, proline, pyruvic acid, riboflavine, sorbitol, thiamine, threonine, thymidine, uridine, xanthine).

**Figure 7.**
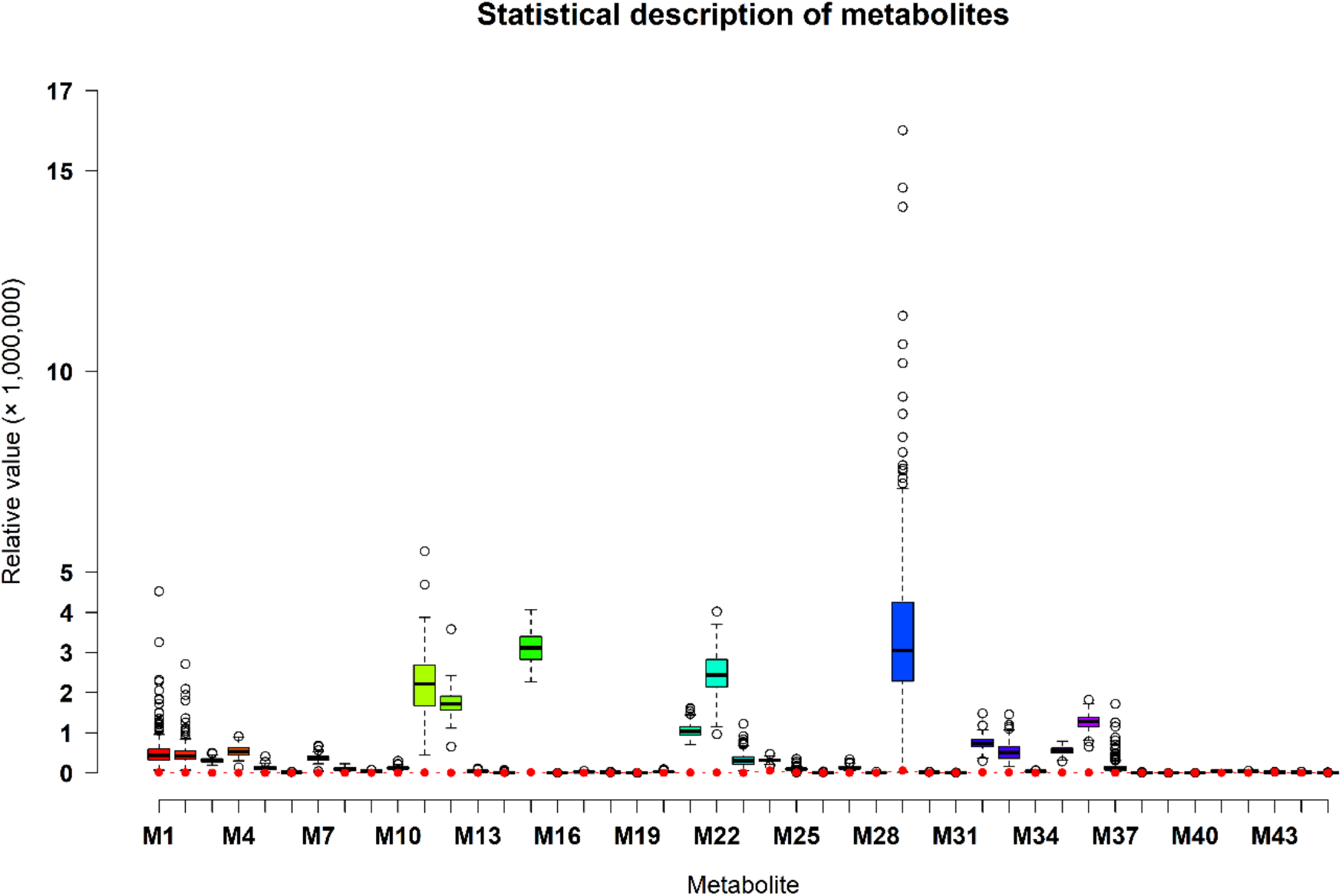
Statistical description of 45 metabolites from combined two sampling times. Note: Red solid circle indicates the limit of detection (LOD) relative value of each metabolite. LOD refers to the lowest value of a metabolite that the LC-MS method can detect. The M1 to M45 indicate the metabolites of 1-hexadecyl-sn-glycero-3-phosphocholine, 1-myristoyl-sn-glycero-3-phosphocholine, (3-Carboxypropyl)trimethylammonium, 4-aminobenzoic acid, 5-methyl-5,6-dihydrouracils, acetaminophen, acetylcarnitine, alanine, arginine, aspartic acid, benzoic acid, carnitine, citrulline, cotinine, creatinine, cytidine, disaccharide, glutamic acid, guanine, guanosine, hypoxanthine, indoleacrylic acid, inosine, isoleucine, isoleucyl proline, isovalerylcarnitine, lactic acid, leucyl methionine, lysoPC(16:0), manNAc, methionine, monosaccharide, nicotine amide, ornithine, phenylalanine, proline, propionylcarnitine, pyruvic acid, riboflavine, sorbitol, thiamine, threonine, thymidine, uridine, xanthine, respectively.

The genotype data was downloaded from NCBI GEO database with Accession Number: GSE144064 with the link: https://www.ncbi.nlm.nih.gov/geo/query/acc.cgi?acc=GSE144064 that were sequenced in the previous study [11]. After removing the markers with duplicated SNP positions (i.e., coordinates) (n = 274), unannotated SNP positions (n = 2618) and no genotypes (n = 3903), 61,721 SNP markers retained. Afterwards, we performed the imputation for missing markers using pedigree (i.e., all the pigs were derived from sixteen-sire families in four generations) by FImpute software (version 3) [13], as the closer relatives usually share longer haplotypes. Therefore, pedigree information could contribute FImpute to achieve more accurate imputation [13,14]. Quality control (QC) for the imputed genotypes was conducted based on the criteria of Hardy-Weinberg equilibrium (HWE > 10^−7^) and minor allele frequency (MAF ≥ 0.001) by PLINK software (version 1.9) [15].

In this study, we also combined the metabolite values from the first sampling time and the second sampling time as an integrated dataset, so each pig had two metabolic values for one metabolite. However, the genotypes of each pig for the metabolite values between the first sampling time and the second sampling time were the same. Finally, the genotypes for the first sampling time and the second sampling time retained 47,109 SNP markers after removing unqualified 9,337 (HWE ≤ 10^−7^) and 5,275 (MAF < 0.001) markers, while the genotypes for the combined two sampling times retained 42,393 SNP markers after removing unqualified 14,053 (HWE ≤ 10^−7^) and 5,275 (MAF < 0.001) markers.

### 4.2. Partial least squares-discriminant analysis and metabolic pathway enrichment for 45 metabolites

The partial least squares-discriminant analysis (PLS-DA) and metabolic pathway analysis for the 45 metabolites were performed by *MetaboAnalyst* software (version 4.0) [16] using *Homo sapiens* as the library. Fishers’ exact test and relative betweenness centrality were used for the over-represented analysis and the pathway impact value calculation (i.e., sum of importance measures of the matched metabolites divided by the sum of the importance measures of all the metabolites), respectively [17]. The first component scores (Component 1) and metabolites enriched in the significant metabolic pathways after false discovery rate (FDR) correction of multiple hypothesis testing (FDR < 0.1) were selected as phenotypes of the transformed metabolites for GWAS. The mean calculated for the metabolites enriched in each significant metabolic pathways was considered as transformed metabolite values, thus, each pathway had one transformed metabolite value (i.e., the mean).

### 4.3. Mixed linear model based association analysis

In our study, GWAS for 45 single metabolites and transformed metabolites (i.e., Component 1 and enriched metabolites) was conducted by mixed liner model based association analysis in GCTA software (version 1.93.0) [18]. The mixed linear model is:

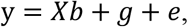

where y is the vector of phenotypes (i.e., metabolites from the first, second and combined two sampling times and the transformed metabolites), *b* is the vector of fixed effects including intercept, breed effects (i.e., Duroc and Landrace pigs), RFI effects (i.e., actual RFI values included as covariates) and SNP effects (i.e., candidate SNPs included as covariates) to be tested, *X* is the design matrix for fixed effects that includes SNP genotype indicators (i.e., 0, 1 or 2), *g* is the vector of polygenic effects as random effects that are the accumulated effects of all SNPs, *e* is the vector of residual effects. The polygenic and residual variances are 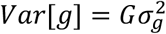 and 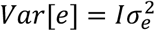, where *G* and *I* are the genetic relationship matrix (GRM) calculated using all SNPs and identity matrix, respectively.

### 4.4. Significant SNPs and their annotated genes

The significant SNPs for GWAS were defined when the *P*-values were less than the threshold after Bonferroni correction for multiple hypothesis testing on genome level. The threshold for metabolites from first and second sampling time was 1.06E-06 (i.e., 0.05/47109), while the threshold for combined two sampling times was 1.18E-06 (i.e., 0.05/42393). Then, the significant SNPs were annotated to the genes and gene components (i.e., promoters, exons and introns) of porcine RefSeq database (Sscrofa10.2/susScr3) downloaded from University of California Santa Cruz (UCSC) genome browser (https://genome.ucsc.edu/cgi-bin/hgTables), where a window of 500 kb was used for the annotation of intergenic regions to neighboring genes.

Linkage disequilibrium (LD) analysis to display the potential haplotypes for the significant SNPs was performed using Haploview software (version 4.2) [19]. SNPs with the distance larger than 500 kb were ignored in the pairwise comparisons for LD analysis.

### 4.5. Gene-based pathway enrichment analysis and gene-metabolite interaction network

We used R package *KEGG.db* (version 3.2.3) of *Sus scrofa* species to annotate SNP-related genes in the gene-based pathway enrichments. Based on the adjusted p-values (p.adjust) < 0.2 under FDR control, the gene-based pathways were finally realized by R package *clusterprofiler* (version 3.12.0) [20]. The gene-gene interaction networks were created by *GeneMANIA* server [21,22] with default settings using Homo sapiens as the library. Then, the gene-metabolite networks for interactions between SNP-related genes and phenotype-related metabolites were created by *MetaboAnalyst* tool [23] with default settings using the same library of Homo sapiens. Significant SNPs associated metabolites based on the low-high RFI pigs were hierarchically clustered by the Ward’s method in Euclidean distance [24]. Then, a heat map for averaged metabolite clustering was visualized by *MetaboAnalyst* tool [16].

## 5. Conclusions

We utilized metabolic and genomic datasets from a total of 108 pigs that were made available for this study from our own previously published studies [6,11] in publicly available data repositories. These studies involved 59 Duroc and 49 Landrace pigs and consisted of data on feed efficiency (RFI), genotype (PorcineSNP80 BeadChip) data and metabolomic data (45 final metabolite datasets derived from LC-MS system). Utilizing these datasets, our main aim was to identify genetic variants (SNPs) that affect 45 different metabolite concentrations in plasma collected at the start and end of the performance testing of pigs categorized as high or low in their feed efficiency, as measured by RFI values. Genome-wide significant genetic variants could be then used as potential genetic or biomarkers in breeding programs for feed efficiency. In order to achieve this main objective, we performed GWAS in the mixed linear model based association analysis and found 152 genome-wide significant SNPs (*P*-value < 1.06E-06) in association with 17 metabolites that included 90 significant SNPs annotated to 52 genes. On chromosome one alone, we found SNPs in strong LD that could be annotated to *FBXL4* and *CCNC*; it consisted of two haplotype blocks, where three SNPs (ALGA0004000, ALGA0004041 and ALGA0004042) were in the intron regions of *FBXL4* and *CCNC*. The interaction network analyses revealed that *CCNC* and *FBXL4* were linked to each other by *N6AMT1* gene and were associated with compounds isovalerylcarnitine and propionylcarnitine. The identified genetic variants and genes affecting important metabolites in high versus low feed efficient pigs could be considered as potential genetic or biomarkers but we recommend that these results are validated in much higher sample size.

## Abbreviations

Component 1: First component score
FC: Fold change
FDR: False discovery rate
LC-MS: Liquid chromatography-mass spectrometry
GRM: Genetic relationship matrix
GWAS: Genome-wide association study
HWE: Hardy-Weinberg equilibrium
LD: Linkage disequilibrium
LOD: Limit of detection
LLOR: Logarithm of likelihood odd ratio
MAF: Minor allele frequency
mGWAS: Metabolite GWAS
NCBI: National Center for Biotechnology Information
PLS-DA: Partial least squares-discriminant analysis
QC: Quality control
QTL: Quantitative trait locus
RFI: Residual feed intake
SNP: Single nucleotide polymorphism
UCSC: University of California Santa Cruz

## Availability of data and materials

All datasets used in this paper are from public repositories. Metabolite data were accessed using MetaboLights accession ID MTBLS1384 with a link: https://www.ebi.ac.uk/metabolights/MTBLS1384. Genotype data were accessed from NCBI GEO Accession Number: GSE144064 with the link: https://www.ncbi.nlm.nih.gov/geo/query/acc.cgi?acc=GSE144064). The details of these datasets can be found in Banerjee et al. (2020) [11] and Carmelo et al. (2020) [6].

## Acknowledgements

Authors thank open access platforms MetaboLights and NCBI-GEO from which we downloaded the datasets for research reported in this study and cited under the section “availability of data and materials”. Authors thank Prof. Claus Thorn Ekstrøm from Faculty of Health and Medical Sciences, University of Copenhagen for his advice in statistical modelling.

## Author information

### Affiliations

Quantitative Genomics, Bioinformatics and Computational Biology Group, Department of Applied Mathematics and Computer Science, Technical University of Denmark, Richard Peterson Plads, Building 324, 2800, Kongens Lyngby, Denmark Xiao Wang & Haja N. Kadarmideen

### Contributions

HNK was a grant holder and Lead PI for FeedOMICS project who conceived and designed the experiments and made the related datasets available in public repositories. XW analyzed the data under the supervision of HNK. XW and HNK wrote the first draft of this paper, which was improved by HNK. All authors have read, revised and approved the final manuscript.

### Corresponding author

Correspondence to Haja N. Kadarmideen.

## Ethics declarations

### Ethics approval and consent to participate

Not applicable as datasets were sourced from public repositories

### Consent for publication

Not applicable.

## Supplementary materials

Supplementary table 1. All the significant SNPs of genome-wide association with chromosome, position and *P*-value information for metabolites from first, second and combined two sampling times.

Supplementary table 2. All the metabolites in association with significant SNPs from first, second and combined two sampling times.

Supplementary figure 1. Manhattan plots of genome-wide association for 43 metabolites (i.e., 1- hexadecyl-sn-glycero-3-phosphocholine (M1), 1-myristoyl-sn-glycero-3-phosphocholine (M2), (3- Carboxypropyl)trimethylammonium (M3), 4-aminobenzoic acid (M4), 5-methyl-5,6-dihydrouracils (M5), acetaminophen (M6), acetylcarnitine (M7), alanine (M8), arginine (M9), aspartic acid (M10), benzoic acid (M11), carnitine (M12), citrulline (M13), cotinine (M14), creatinine (M15), cytidine (M16), disaccharide (M17), glutamic acid (M18), guanine (M19), guanosine (M20), hypoxanthine (M21), indoleacrylic acid (M22), inosine (M23), isoleucine (M24), isoleucyl proline (M25), lactic acid (M27), leucyl methionine (M28), lysoPC(16:0) (M29), manNAc (M30), methionine (M31), monosaccharide (M32), nicotine amide (M33), ornithine (M34), phenylalanine (M35), proline (M36), pyruvic acid (M38), riboflavin (M39), sorbitol (M40), thiamine (M41), threonine (M42), thymidine (M43), uridine (M44), xanthine (M45) from supplementary figure 1.1 to supplementary figure 1.43). Note: Y-axis indicates the log_10_(*P*-value). Blue dotted and red solid lines indicate the genome-wide threshold of 0.05 and 0.01 after Bonferroni multiple testing, respectively. The three tracks indicate the metabolites from first sampling time, second sampling time and combined two sampling times, respectively, from outside to inside.

Supplementary figure 2. Linkage disequilibrium (LD) pattern for all significant SNPs. Note: Solid line triangle indicates LD. One square indicates LD level (r^2^) between two SNPs and the squares are colored by D’/LLOR standard scheme. D’/LLOR standard scheme is that red indicates LLOR > 2, D’ = 1; pink indicates LLOR > 2, D’ < 1; blue indicates LLOR < 2, D’ = 1; white indicates LLOR < 2, D’ < 1. LLOR is the logarithm of likelihood odds ratio and the reliable index to measure D’.

## References

1. Burton, P.R.; Clayton, D.G.; Cardon, L.R.; Craddock, N.; Deloukas, P.; Duncanson, A.; Kwiatkowski, D.P.; McCarthy, M.I.; Ouwehand, W.H.; Samani, N.J.; et al. Genome-wide association study of 14,000 cases of seven common diseases and 3,000 shared controls. Nature 2007.

2. Suravajhala, P.; Kogelman, L.J.A.; Kadarmideen, H.N. Multi-omic data integration and analysis using systems genomics approaches: Methods and applications In animal production, health and welfare. Genet. Sel. Evol. 2016.

3. Beebe, K.; Kennedy, A.D. Sharpening Precision Medicine by a Thorough Interrogation of Metabolic Individuality. Comput. Struct. Biotechnol. J. 2016.

4. Fiehn, O. Metabolomics – The link between genotypes and phenotypes. Plant Mol. Biol. 2002.

5. Liu, H.; Chen, Y.; Ming, D.; Wang, J.; Li, Z.; Ma, X.; Wang, J.; van Milgen, J.; Wang, F. Integrative analysis of indirect calorimetry and metabolomics profiling reveals alterations in energy metabolism between fed and fasted pigs. J. Anim. Sci. Biotechnol. 2018.

6. Carmelo, V.A.O.; Banerjee, P.; da Silva Diniz, W.J.; Kadarmideen, H.N. Metabolomic networks and pathways associated with feed efficiency and related-traits in Duroc and Landrace pigs. Sci. Rep. 2020.

7. Kenéz, Á.; Dänicke, S.; Rolle-Kampczyk, U.; von Bergen, M.; Huber, K. A metabolomics approach to characterize phenotypes of metabolic transition from late pregnancy to early lactation in dairy cows. Metabolomics 2016.

8. Wang, X.; Kadarmideen, H.N. Metabolomics Analyses in High-Low Feed Efficient Dairy Cows Reveal Novel Biochemical Mechanisms and Predictive Biomarkers. Metabolites 2019, 9, 151.

9. Gieger, C.; Geistlinger, L.; Altmaier, E.; De Angelis, M.H.; Kronenberg, F.; Meitinger, T.; Mewes, H.W.; Wichmann, H.E.; Weinberger, K.M.; Adamski, J.; et al. Genetics meets metabolomics: A genome-wide association study of metabolite profiles in human serum. PLoS Genet. 2008.

10. Do, D.N.; Strathe, A.B.; Ostersen, T.; Pant, S.D.; Kadarmideen, H.N. Genome-wide association and pathway analysis of feed efficiency in pigs reveal candidate genes and pathways for residual feed intake. Front. Genet. 2014.

11. Banerjee, P.; Adriano, V.; Carmelo, O.; Kadarmideen, H.N. Genome-Wide Epistatic Interaction Networks Affecting Feed Efficiency in Duroc and Landrace Pigs. 2020, 11, 1–13.

12. Wang, X.; Ma, P.; Liu, J.; Zhang, Q.; Zhang, Y.; Ding, X.; Jiang, L.; Wang, Y.; Zhang, Y.; Sun, D.; et al. Genome-wide association study in Chinese Holstein cows reveal two candidate genes for somatic cell score as an indicator for mastitis susceptibility. BMC Genet. 2015.

13. Sargolzaei, M.; Chesnais, J.P.; Schenkel, F.S. A new approach for efficient genotype imputation using information from relatives. BMC Genomics 2014, 15, 478.

14. Wang, X.; Su, G.; Hao, D.; Lund, M.S.; Kadarmideen, H.N. Comparisons of improved genomic predictions generated by different imputation methods for genotyping by sequencing data in livestock populations. J. Anim. Sci. Biotechnol. 2020.

15. Purcell, S.; Neale, B.; Todd-Brown, K.; Thomas, L.; Ferreira, M.A.R.; Bender, D.; Maller, J.; Sklar, P.; De Bakker, P.I.W.; Daly, M.J.; et al. PLINK: A tool set for whole-genome association and population-based linkage analyses. Am. J. Hum. Genet. 2007.

16. Xia, J.; Wishart, D.S.; Valencia, A. MetPA: A web-based metabolomics tool for pathway analysis and visualization. In Proceedings of the Bioinformatics; 2011.

17. Xia, J.; Wishart, D.S. Web-based inference of biological patterns, functions and pathways from metabolomic data using MetaboAnalyst. Nat. Protoc. 2011.

18. Yang, J.; Lee, S.H.; Goddard, M.E.; Visscher, P.M. GCTA: A tool for genome-wide complex trait analysis. Am. J. Hum. Genet. 2011.

19. Barrett, J.C.; Fry, B.; Maller, J.; Daly, M.J. Haploview: Analysis and visualization of LD and haplotype maps. Bioinformatics 2005.

20. Yu, G.; Wang, L.-G.; Han, Y.; He, Q.-Y. clusterProfiler: an R Package for Comparing Biological Themes Among Gene Clusters. Omi. A J. Integr. Biol. 2012.

21. Mostafavi, S.; Ray, D.; Warde-Farley, D.; Grouios, C.; Morris, Q. GeneMANIA: A real-time multiple association network integration algorithm for predicting gene function. Genome Biol. 2008.

22. Warde-Farley, D.; Donaldson, S.L.; Comes, O.; Zuberi, K.; Badrawi, R.; Chao, P.; Franz, M.; Grouios, C.; Kazi, F.; Lopes, C.T.; et al. The GeneMANIA prediction server: Biological network integration for gene prioritization and predicting gene function. Nucleic Acids Res. 2010.

23. Chong, J.; Soufan, O.; Li, C.; Caraus, I.; Li, S.; Bourque, G.; Wishart, D.S.; Xia, J. MetaboAnalyst 4.0: Towards more transparent and integrative metabolomics analysis. Nucleic Acids Res. 2018.

24. Ward, J.H. Hierarchical Grouping to Optimize an Objective Function. J. Am. Stat. Assoc. 1963.

25. Helke, K.L.; Nelson, K.N.; Sargeant, A.M.; Jacob, B.; McKeag, S.; Haruna, J.; Vemireddi, V.; Greeley, M.; Brocksmith, D.; Navratil, N.; et al. Pigs in Toxicology: Breed Differences in Metabolism and Background Findings. Toxicol. Pathol. 2016.

26. Picone, G.; Zappaterra, M.; Luise, D.; Trimigno, A.; Capozzi, F.; Motta, V.; Davoli, R.; Nanni Costa, L.; Bosi, P.; Trevisi, P. Metabolomics characterization of colostrum in three sow breeds and its influences on piglets’ survival and litter growth rates. J. Anim. Sci. Biotechnol. 2018.

27. Sundekilde, U.K.; Frederiksen, P.D.; Clausen, M.R.; Larsen, L.B.; Bertram, H.C. Relationship between the metabolite profile and technological properties of bovine milk from two dairy breeds elucidated by NMR-based metabolomics. J. Agric. Food Chem. 2011.

28. Goto, T.; Mori, H.; Shiota, S.; Tomonaga, S. Metabolomics approach reveals the effects of breed and feed on the composition of chicken eggs. Metabolites 2019.

29. Yin, J.D.; Shang, X.G.; Li, D.F.; Wang, F.L.; Guan, Y.F.; Wang, Z.Y. Effects of dietary conjugated linoleic acid on the fatty acid profile and cholesterol content of egg yolks from different breeds of layers. Poult. Sci. 2008.

30. Middleton, R.P.; Lacroix, S.; Scott-Boyer, M.P.; Dordevic, N.; Kennedy, A.D.; Slusky, A.R.; Carayol, J.; Petzinger-Germain, C.; Beloshapka, A.; Kaput, J. Metabolic Differences between Dogs of Different Body Sizes. J. Nutr. Metab. 2017.

31. Zhu, G.; Wang, S.; Huang, Z.; Zhang, S.; Liao, Q.; Zhang, C.; Lin, T.; Qin, M.; Peng, M.; Yang, C.; et al. Rewiring of the Fruit Metabolome in Tomato Breeding. Cell 2018.

32. Do, D.N.; Strathe, A.B.; Jensen, J.; Mark, T.; Kadarmideen, H.N. Genetic parameters for different measures of feed efficiency and related traits in boars of three pig breeds. J. Anim. Sci. 2013.

33. Ding, R.; Yang, M.; Wang, X.; Quan, J.; Zhuang, Z.; Zhou, S.; Li, S.; Xu, Z.; Zheng, E.; Cai, G.; et al. Genetic architecture of feeding behavior and feed efficiency in a Duroc pig population. Front. Genet. 2018.

34. Dekaney, C.M.; Wu, G.; Jaeger, L.A. Gene expression and activity of enzymes in the arginine biosynthetic pathway in porcine fetal small intestine. Pediatr. Res. 2003.

35. Elia, I.; Broekaert, D.; Christen, S.; Boon, R.; Radaelli, E.; Orth, M.F.; Verfaillie, C.; Grünewald, T.G.P.; Fendt, S.M. Proline metabolism supports metastasis formation and could be inhibited to selectively target metastasizing cancer cells. Nat. Commun. 2017.

36. Gilbert, H.; Riquet, J.; Gruand, J.; Billon, Y.; Fève, K.; Sellier, P.; Noblet, J.; Bidanel, J.P. Detecting QTL for feed intake traits and other performance traits in growing pigs in a Piétrain-Large White backcross. Animal 2010.

37. Shirali, M.; Duthie, C.A.; Doeschl-Wilson, A.; Knap, P.W.; Kanis, E.; van Arendonk, J.A.M.; Roehe, R. Novel insight into the genomic architecture of feed and nitrogen efficiency measured by residual energy intake and nitrogen excretion in growing pigs. BMC Genet. 2013.

38. Sanchez, M.P.; Tribout, T.; Iannuccelli, N.; Bouffaud, M.; Servin, B.; Tenghe, A.; Dehais, P.; Muller, N.; Del Schneider, M.P.; Mercat, M.J.; et al. A genome-wide association study of production traits in a commercial population of Large White pigs: Evidence of haplotypes affecting meat quality. Genet. Sel. Evol. 2014.

39. Onteru, S.K.; Gorbach, D.M.; Young, J.M.; Garrick, D.J.; Dekkers, J.C.M.; Rothschild, M.F. Whole Genome Association Studies of Residual Feed Intake and Related Traits in the Pig. PLoS One 2013.

40. Do, D.N.; Ostersen, T.; Strathe, A.B.; Mark, T.; Jensen, J.; Kadarmideen, H.N. Genome-wide association and systems genetic analyses of residual feed intake, daily feed consumption, backfat and weight gain in pigs. BMC Genet. 2014.

41. Makowski, L.; Noland, R.C.; Koves, T.R.; Xing, W.; Ilkayeva, O.R.; Muehlbauer, M.J.; Stevens, R.D.; Muoio, D.M. Metabolic profiling of PPARα-/- mice reveals defects in carnitine and amino acid homeostasis that are partially reversed by oral carnitine supplementation. FASEB J. 2009.

42. Kerner, J.; Froseth, J.A.; Miller, E.R.; Bieber, L.L. A study of the acylcarnitine content of sows’ colostrum, milk and newborn piglet tissues: Demonstration of high amounts of isovaleryl-carnitine in colostrum and milk. J. Nutr. 1984.

43. Bartels, G.L.; Remme, W.J.; Holwerda, K.J.; Kruijssen, D.A.C.M. Anti-ischaemic efficacy of L-propionylcarnitine – a promising novel metabolic approach to ischaemia? Eur. Heart J. 1996.

44. Lew, D.J.; Dulić, V.; Reed, S.I. Isolation of three novel human cyclins by rescue of G1 cyclin (cln) function in yeast. Cell 1991.

45. Arai, E.; Sakamoto, H.; Ichikawa, H.; Totsuka, H.; Chiku, S.; Gotoh, M.; Mori, T.; Nakatani, T.; Ohnami, S.; Nakagawa, T.; et al. Multilayer-omics analysis of renal cell carcinoma, including the whole exome, methylome and transcriptome. Int. J. Cancer 2014.

46. Miyata, Y.; Liu, Y.; Jankovic, V.; Sashida, G.; Lee, J.M.; Shieh, J.H.; Naoe, T.; Moore, M.; Nimer, S.D. Cyclin C regulates human hematopoietic stem/progenitor sell quiescence. Stem Cells 2010.

47. Bondi, J.; Husdal, A.; Bukholm, G.; Nesland, J.M.; Bakka, A.; Bukholm, I.R.K. Expression and gene amplification of primary (A, B1, D1, D3, and E) and secondary (C and H) cyclins in colon adenocarcinomas and correlation with patient outcome. J. Clin. Pathol. 2005.

48. El-Hattab, A.W.; Dai, H.; Almannai, M.; Wang, J.; Faqeih, E.A.; Al Asmari, A.; Saleh, M.A.M.; Elamin, M.A.O.; Alfadhel, M.; Alkuraya, F.S.; et al. Molecular and clinical spectra of FBXL4 deficiency. Hum. Mutat. 2017.

49. Ballout, R.A.; Alam, C. Al; Bonnen, P.E.; Huemer, M.; El-Hattab, A.W.; Shbarou, R. FBXL4-related mitochondrial DNA depletion syndrome 13 (MTDPS13): A case report with a comprehensive mutation review. Front. Genet. 2019.

50. Bonnen, P.E.; Yarham, J.W.; Besse, A.; Wu, P.; Faqeih, E.A.; Al-Asmari, A.M.; Saleh, M.A.M.; Eyaid, W.; Hadeel, A.; He, L.; et al. Mutations in FBXL4 cause mitochondrial encephalopathy and a disorder of mitochondrial DNA maintenance. Am. J. Hum. Genet. 2013.

51. Stankiewicz, E.; Mao, X.; Mangham, D.C.; Xu, L.; Yeste-Velasco, M.; Fisher, G.; North, B.; Chaplin, T.; Young, B.; Wang, Y.; et al. Identification of FBXL4 as a Metastasis Associated Gene in Prostate Cancer. Sci. Rep. 2017.

52. Li, Q.; Li, Y.; Wang, X.; Qi, J.; Jin, X.; Tong, H.; Zhou, Z.; Zhang, Z.C.; Han, J. Fbxl4 Serves as a Clock Output Molecule that Regulates Sleep through Promotion of Rhythmic Degradation of the GABAA Receptor. Curr. Biol. 2017.

53. Li, Y.; Yang, S.L.; Tang, Z.L.; Cui, W.T.; Mu, Y.L.; Chu, M.X.; Zhao, S.H.; Wu, Z.F.; Li, K.; Peng, K.M. Expression and SNP association analysis of porcine FBXL4 gene. Mol. Biol. Rep. 2010.

54. Cabrera, R.A.; Usry, J.L.; Arrellano, C.; Nogueira, E.T.; Kutschenko, M.; Moeser, A.J.; Odle, J. Effects of creep feeding and supplemental glutamine or glutamine plus glutamate (Aminogut) on pre- and post-weaning growth performance and intestinal health of piglets. J. Anim. Sci. Biotechnol. 2013.

55. Hsu, C.B.; Lee, J.W.; Huang, H.J.; Wang, C.H.; Lee, T.T.; Yen, H.T.; Yu, B. Effects of supplemental glutamine on growth performance, plasma parameters and LPS-induced immune response of weaned barrows after castration. Asian-Australasian J. Anim. Sci. 2012.

