## Supplementary File for "Metabolite genome-wide association study (mGWAS) and gene-metabolite interaction network analysis reveal potential biomarkers for feed efficiency in pigs"

Supplementary table 1. All the significant SNPs of genome-wide association with chromosome, position and *P*-value information for metabolites from first, second and combined two sampling times.

| Significant SNP name | Chromosome | Position | Nearest gene | Distance | Associated metabolite number | Metabolite from first sampling time (*P*-value) | Metabolite from second sampling time (*P*-value) | Metabolite from combined two sampling times (*P*-value) |
| --- | --- | --- | --- | --- | --- | --- | --- | --- |
| 18_2566820 | 18 | 2566820 | *SHH* (NM_001244513) | 97973 | 1 |  | Aspartic acid (2.29E-07) |  |
| 2_29041727 | 2 | 29041727 | *CAT* (NM_214301) | 140893 | 1 |  | Nicotine amide (4.71E-07) |  |
| 3_118102178 | 3 | 118102178 | *RBKS* (NM_001315776) | 172300 | 1 |  | Aspartic acid (2.29E-07) |  |
| ALGA0000375 | 1 | 5395131 |  |  | 1 |  | Isoleucyl proline (3.57E-07) |  |
| ALGA0014130 | 2 | 86940572 | *F2R* (NM_001244372) | 339212 | 1 | Pyruvic acid (1.25E-07) |  |  |
| ALGA0043837 | 7 | 100079759 | *SYNJ2BP* (NM_001244991) | 149115 | 1 |  | Isoleucyl proline (3.57E-07) |  |
| ALGA0046501 | 8 | 12844147 | *LCORL* (NM_001195345) | 78106 | 1 |  | Isoleucyl proline (3.57E-07) |  |
| ALGA0046512 | 8 | 13111033 | *LCORL* (NM_001195345) | 344992 | 1 |  | Isoleucyl proline (3.57E-07) |  |
| ALGA0049375 | 8 | 127824811 | *NFKB1* (NM_001048232) | 160810 | 1 | Pyruvic acid (7.57E-07) |  |  |
| ALGA0049385 | 8 | 127976364 | *NFKB1* (NM_001048232) | 312363 | 1 | Pyruvic acid (7.57E-07) |  |  |
| ALGA0061605 | 11 | 26591544 | *MTRF1* (NM_001243580) | 5^th^ intron | 1 |  | Aspartic acid (2.29E-07) |  |
| ALGA0072779 | 13 | 177981732 | *EPHA3* (NM_001195335) | 171617 | 1 | Cotinine (4.18E-09) |  |  |
| ALGA0083708 | 15 | 2069154 |  |  | 1 |  | Guanine (1.63E-07) |  |
| ALGA0104701 | 17 | 8537481 |  |  | 1 |  | Guanine (8.32E-08) |  |
| ASGA0040302 | 8 | 143127778 | *CDS1* (NM_001044534) | 312027 | 1 | 1-Hexadecyl-sn-glycero-3-phosphocholine (7.02E-07) |  |  |
| ASGA0042433 | 9 | 34828091 |  |  | 1 | Pyruvic acid (6.93E-07) |  |  |
| ASGA0042436 | 9 | 34883824 |  |  | 1 | Pyruvic acid (6.93E-07) |  |  |
| ASGA0054868 | 12 | 49919795 | *MIR212* (NR_128427) | 96355 | 1 |  | Guanine (8.32E-08) |  |
| ASGA0078611 | 18 | 2676317 | *SHH* (NM_001244513) | 2512 | 1 |  | Propionylcarnitine (2.98E-07) |  |
| ASGA0083287 | 13 | 3868357 | *DPH3* (NM_001243616) | 282233 | 1 | Creatinine (4.79E-07) |  |  |
| ASGA0094610 | 17 | 48307754 |  |  | 1 | LysoPC(16:0) (5.29E-07) |  |  |
| ASGA0095299 | 5 | 83455803 | *TXNRD1* (NM_214154) | 202024 | 1 |  |  | Monosaccharide (9.09E-07) |
| DRGA0004012 | 3 | 71708951 | *DOK1* (NM_001143705) | 39335 | 1 |  | Propionylcarnitine (7.66E-08) |  |
| DRGA0008955 | 8 | 143055109 | *CDS1* (NM_001044534) | 384696 | 1 | 1-Hexadecyl-sn-glycero-3-phosphocholine (7.02E-07) |  |  |
| H3GA0007606 | 2 | 132433332 |  |  | 1 | Isovalerylcarnitine (7.84E-07) |  |  |
| H3GA0015652 | 5 | 10709671 | *MIR9834* (NR_128521) | 331400 | 1 |  | 1-Myristoyl-sn-glycero-3-phosphocholine (5.74E-09) |  |
| H3GA0026947 | 9 | 34841808 |  |  | 1 | Pyruvic acid (6.93E-07) |  |  |
| H3GA0028115 | 9 | 128659759 |  |  | 1 |  | Isoleucyl proline (3.57E-07) |  |
| H3GA0035928 | 13 | 27911595 | *CTNNB1* (NM_214367) | 244338 | 1 |  | Propionylcarnitine (5.78E-07) |  |
| H3GA0055935 | 9 | 2349391 |  |  | 1 |  | Aspartic acid (2.29E-07) |  |
| INRA0031485 | 9 | 34920723 |  |  | 1 | Pyruvic acid (6.93E-07) |  |  |
| M1GA0003090 | 2 | 132449420 |  |  | 1 | Isovalerylcarnitine (7.84E-07) |  |  |
| M1GA0015335 | 11 | 80613144 |  |  | 1 | Cotinine (4.18E-09) |  |  |
| MARC0001720 | 9 | 0 |  |  | 1 | Pyruvic acid (6.93E-07) |  |  |
| MARC0010841 | 5 | 10759678 | *MIR9834* (NR_128521) | 381407 | 1 |  | 1-Myristoyl-sn-glycero-3-phosphocholine (5.74E-09) |  |
| MARC0023109 | 14 | 149350120 |  |  | 1 |  | Pyruvic acid (1.07E-07) |  |
| MARC0027232 | 17 | 48876667 | *ZHX3* (NM_001258405) | 371390 | 1 | LysoPC(16:0) (5.29E-07) |  |  |
| MARC0040505 | 9 | 50293805 | *FXYD2* (NM_001014427) | 44876 | 1 |  |  | Citrulline (4.75E-07) |
| MARC0060063 | 8 | 142707308 |  |  | 1 | 1-Hexadecyl-sn-glycero-3-phosphocholine (7.02E-07) |  |  |
| MARC0083382 | 15 | 140091862 |  |  | 1 |  | Carnitine (2.40E-07) |  |
| MARC0092124 | 9 | 34900287 |  |  | 1 | Pyruvic acid (6.93E-07) |  |  |
| MARC0110390 | 2 | 83663964 | *SFXN1* (NM_001098602) | 7^th^ intron | 1 | Pyruvic acid (1.25E-07) |  |  |
| WU_10.2_10_71866847 | 10 | 71866847 | *AKR1C2* (NM_001044570) | 16292 | 1 |  | Propionylcarnitine (1.28E-07) |  |
| WU_10.2_12_49599992 | 12 | 49599992 | *INPP5K* (NM_001190292) | 3831 | 1 |  | Guanine (8.32E-08) |  |
| WU_10.2_12_49979240 | 12 | 49979240 | *MIR212* (NR_128427) | 36910 | 1 |  | Guanine (8.32E-08) |  |
| WU_10.2_13_204594763 | 13 | 204594763 | *CLDN8* (NM_001161646) | 428133 | 1 |  | Isoleucyl proline (4.20E-07) |  |
| WU_10.2_13_204608919 | 13 | 204608919 | *CLDN8* (NM_001161646) | 442289 | 1 |  | Isoleucyl proline (4.20E-07) |  |
| WU_10.2_13_204631833 | 13 | 204631833 | *CLDN8* (NM_001161646) | 465203 | 1 |  | Isoleucyl proline (4.20E-07) |  |
| WU_10.2_17_48295280 | 17 | 48295280 |  |  | 1 | LysoPC(16:0) (5.29E-07) |  |  |
| WU_10.2_2_132463311 | 2 | 132463311 |  |  | 1 | Isovalerylcarnitine (7.84E-07) |  |  |
| WU_10.2_2_132503341 | 2 | 132503341 |  |  | 1 | Isovalerylcarnitine (7.84E-07) |  |  |
| WU_10.2_2_132570224 | 2 | 132570224 |  |  | 1 | Isovalerylcarnitine (7.84E-07) |  |  |
| WU_10.2_6_135312468 | 6 | 135312468 | *LEPROT* (NM_001145388) | 67384 | 1 | 1-Hexadecyl-sn-glycero-3-phosphocholine (1.05E-06) |  |  |
| WU_10.2_7_100175430 | 7 | 100175430 | *SYNJ2BP* (NM_001244991) | 53444 | 1 |  | Isoleucyl proline (3.57E-07) |  |
| WU_10.2_8_12998407 | 8 | 12998407 | *LOCRL* (NM_001195345) | 232366 | 1 |  | Isoleucyl proline (3.57E-07) |  |
| WU_10.2_8_142992366 | 8 | 142992366 | *CDS1* (NM_001044534) | 447439 | 1 | 1-Hexadecyl-sn-glycero-3-phosphocholine (7.02E-07) |  |  |
| WU_10.2_9_128572226 | 9 | 128572226 |  |  | 1 |  | Isoleucyl proline (3.57E-07) |  |
| WU_10.2_X_104031384 | X | 104031384 |  |  | 1 |  | Lactic acid (3.14E-07) |  |
| WU_10.2_X_37114281 | X | 37114281 | *GP91-PHOX* (NM_214043) | 253546 | 1 |  | 1-Hexadecyl-sn-glycero-3-phosphocholine (4.85E-09) |  |
| ALGA0003891 | 1 | 69946212 | *LOC780435* (NM_001078684) | 226447 | 2 |  | Isovalerylcarnitine (2.79E-08), Propionylcarnitine (8.32E-10) |  |
| ALGA0003900 | 1 | 70712755 |  |  | 2 |  | Isovalerylcarnitine (2.79E-08), Propionylcarnitine (8.32E-10) |  |
| ALGA0003935 | 1 | 72365650 | *FHL5* (NM_001243314) | 244726 | 2 |  | Isovalerylcarnitine (2.79E-08), Propionylcarnitine (8.32E-10) |  |
| ALGA0003952 | 1 | 72982828 |  |  | 2 |  | Isovalerylcarnitine (2.79E-08), Propionylcarnitine (8.32E-10) |  |
| ALGA0003953 | 1 | 73100657 |  |  | 2 |  | Isovalerylcarnitine (2.79E-08), Propionylcarnitine (8.32E-10) |  |
| ALGA0003995 | 1 | 74410201 | *FBXL4* (NM_001171752) | 12232 | 2 |  | Isovalerylcarnitine (2.79E-08), Propionylcarnitine (8.32E-10) |  |
| ALGA0004000 | 1 | 74467285 | *FBXL4* (NM_001171752) | 6^th^ intron | 2 |  | Isovalerylcarnitine (2.79E-08), Propionylcarnitine (8.32E-10) |  |
| ALGA0004002 | 1 | 74502751 | *FBXL4* (NM_001171752) | 32524 | 2 |  | Isovalerylcarnitine (2.79E-08), Propionylcarnitine (8.32E-10) |  |
| ALGA0004005 | 1 | 74583833 | *FBXL4* (NM_001171752) | 113606 | 2 |  | Isovalerylcarnitine (2.79E-08), Propionylcarnitine (8.32E-10) |  |
| ALGA0004006 | 1 | 74597318 | *FBXL4* (NM_001171752) | 127091 | 2 |  | Isovalerylcarnitine (2.79E-08), Propionylcarnitine (8.32E-10) |  |
| ALGA0004024 | 1 | 74916127 | *PNISR* (NM_001113439) | 79963 | 2 |  | Isovalerylcarnitine (2.79E-08), Propionylcarnitine (8.32E-10) |  |
| ALGA0004041 | 1 | 75151870 | *CCNC* (NM_001190160) | 1^st^ intron | 2 |  | Isovalerylcarnitine (2.79E-08), Propionylcarnitine (8.32E-10) |  |
| ALGA0004042 | 1 | 75167426 | *CCNC* (NM_001190160) | 9^th^ intron | 2 |  | Isovalerylcarnitine (2.79E-08), Propionylcarnitine (8.32E-10) |  |
| ALGA0004046 | 1 | 75219602 | *CCNC* (NM_001190160) | 44247 | 2 |  | Isovalerylcarnitine (2.79E-08), Propionylcarnitine (8.32E-10) |  |
| ALGA0004048 | 1 | 75398012 | *MCHR2* (NM_001044609) | 165694 | 2 |  | Isovalerylcarnitine (2.79E-08), Propionylcarnitine (8.32E-10) |  |
| ALGA0004073 | 1 | 75749595 | *MCHR2* (NM_001044609) | 167075 | 2 |  | Isovalerylcarnitine (2.79E-08), Propionylcarnitine (8.32E-10) |  |
| ALGA0004090 | 1 | 76136641 | *SIM1* (NM_001172585) | 65411 | 2 |  | Isovalerylcarnitine (2.79E-08), Propionylcarnitine (8.32E-10) |  |
| ALGA0004093 | 1 | 76340224 | *SIM1* (NM_001172585) | 268994 | 2 |  | Isovalerylcarnitine (2.79E-08), Propionylcarnitine (8.32E-10) |  |
| ALGA0004143 | 1 | 77325351 |  |  | 2 |  | Isovalerylcarnitine (2.79E-08), Propionylcarnitine (8.32E-10) |  |
| ALGA0004148 | 1 | 77518365 |  |  | 2 |  | Isovalerylcarnitine (2.79E-08), Propionylcarnitine (8.32E-10) |  |
| ALGA0004169 | 1 | 78021030 |  |  | 2 |  | Isovalerylcarnitine (2.79E-08), Propionylcarnitine (8.32E-10) |  |
| ALGA0004173 | 1 | 78233457 |  |  | 2 |  | Isovalerylcarnitine (2.79E-08), Propionylcarnitine (8.32E-10) |  |
| ALGA0004177 | 1 | 78389441 |  |  | 2 |  | Isovalerylcarnitine (2.79E-08), Propionylcarnitine (8.32E-10) |  |
| ALGA0028664 | 4 | 129539293 | *RTCA* (NM_001243470) | 35897 | 2 |  | Glutamic acid (3.57E-10) | Glutamic acid (4.80E-08) |
| ALGA0103368 | 15 | 150174862 |  |  | 2 | Isovalerylcarnitine (9.57E-08) |  | Isovalerylcarnitine (4.19E-07) |
| ASGA0003182 | 1 | 69781723 | *LOC780435* (NM_001078684) | 61958 | 2 |  | Isovalerylcarnitine (2.79E-08), Propionylcarnitine (8.32E-10) |  |
| ASGA0003194 | 1 | 70950692 |  |  | 2 |  | Isovalerylcarnitine (2.79E-08), Propionylcarnitine (8.32E-10) |  |
| ASGA0003235 | 1 | 73520354 |  |  | 2 |  | Isovalerylcarnitine (2.79E-08), Propionylcarnitine (8.32E-10) |  |
| ASGA0003288 | 1 | 75602201 | *MCHR2* (NM_001044609) | 19681 | 2 |  | Isovalerylcarnitine (2.79E-08), Propionylcarnitine (8.32E-10) |  |
| ASGA0003312 | 1 | 76189831 | *SIM1* (NM_001172585) | 118601 | 2 |  | Isovalerylcarnitine (2.79E-08), Propionylcarnitine (8.32E-10) |  |
| ASGA0003314 | 1 | 76215472 | *SIM1* (NM_001172585) | 144242 | 2 |  | Isovalerylcarnitine (2.79E-08), Propionylcarnitine (8.32E-10) |  |
| ASGA0003315 | 1 | 76438140 | *SIM1* (NM_001172585) | 366910 | 2 |  | Isovalerylcarnitine (2.79E-08), Propionylcarnitine (8.32E-10) |  |
| ASGA0003317 | 1 | 76416357 | *SIM1* (NM_001172585) | 345127 | 2 |  | Isovalerylcarnitine (2.79E-08), Propionylcarnitine (8.32E-10) |  |
| ASGA0003333 | 1 | 78347698 |  |  | 2 |  | Isovalerylcarnitine (2.79E-08), Propionylcarnitine (8.32E-10) |  |
| ASGA0003335 | 1 | 78587646 |  |  | 2 |  | Isovalerylcarnitine (2.79E-08), Propionylcarnitine (8.32E-10) |  |
| ASGA0023013 | 4 | 135677951 | *TMED5* (NM_001243695) | 96353 | 2 |  | Glutamic acid (3.57E-10) | Glutamic acid (4.80E-08) |
| ASGA0025965 | 5 | 68149464 | *C5H12orf4* (NM_001243412) | 4228 | 2 |  | Glutamic acid (3.57E-10) | Glutamic acid (4.80E-08) |
| ASGA0044546 | 9 | 133959774 | *TOR1AIP2* (NM_001243389) | 487430 | 2 |  | Carnitine (1.24E-08) | Carnitine (5.24E-08) |
| ASGA0057312 | 13 | 40238900 |  |  | 2 |  | Isovalerylcarnitine (2.79E-08), Propionylcarnitine (8.32E-10) |  |
| ASGA0083304 | 1 | 77011485 |  |  | 2 |  | Isovalerylcarnitine (2.79E-08), Propionylcarnitine (8.32E-10) |  |
| DRGA0000994 | 1 | 66455394 |  |  | 2 |  | Isovalerylcarnitine (2.79E-08), Propionylcarnitine (8.32E-10) |  |
| DRGA0001072 | 1 | 70784564 |  |  | 2 |  | Isovalerylcarnitine (2.79E-08), Propionylcarnitine (8.32E-10) |  |
| DRGA0001073 | 1 | 70898747 |  |  | 2 |  | Isovalerylcarnitine (2.79E-08), Propionylcarnitine (8.32E-10) |  |
| H3GA0001865 | 1 | 67809463 |  |  | 2 |  | Isovalerylcarnitine (2.79E-08), Propionylcarnitine (8.32E-10) |  |
| H3GA0001937 | 1 | 75626511 | *MCHR2* (NM_001044609) | 43991 | 2 |  | Isovalerylcarnitine (2.79E-08), Propionylcarnitine (8.32E-10) |  |
| H3GA0001949 | 1 | 76297353 | *SIM1* (NM_001172585) | 226123 | 2 |  | Isovalerylcarnitine (2.79E-08), Propionylcarnitine (8.32E-10) |  |
| H3GA0001956 | 1 | 77402476 |  |  | 2 |  | Isovalerylcarnitine (2.79E-08), Propionylcarnitine (8.32E-10) |  |
| H3GA0001966 | 1 | 78423848 |  |  | 2 |  | Isovalerylcarnitine (2.79E-08), Propionylcarnitine (8.32E-10) |  |
| H3GA0014734 | 4 | 135234038 | *DNTTIP2* (NM_001243674) | 180160 | 2 |  | Glutamic acid (3.57E-10) | Glutamic acid (4.80E-08) |
| H3GA0046845 | 16 | 65033887 | *MAT2B* (NM_001142832) | 62817 | 2 |  | Isovalerylcarnitine (2.79E-08), Propionylcarnitine (8.32E-10) |  |
| INRA0002726 | 1 | 74096746 | *FBXL4* (NM_001171752) | 325687 | 2 |  | Isovalerylcarnitine (2.79E-08), Propionylcarnitine (8.32E-10) |  |
| INRA0002819 | 1 | 77424826 |  |  | 2 |  | Isovalerylcarnitine (2.79E-08), Propionylcarnitine (8.32E-10) |  |
| INRA0002820 | 1 | 77457737 |  |  | 2 |  | Isovalerylcarnitine (2.79E-08), Propionylcarnitine (8.32E-10) |  |
| INRA0002823 | 1 | 77616272 |  |  | 2 |  | Isovalerylcarnitine (2.79E-08), Propionylcarnitine (8.32E-10) |  |
| MARC0021047 | 1 | 78061517 |  |  | 2 |  | Isovalerylcarnitine (2.79E-08), Propionylcarnitine (8.32E-10) |  |
| MARC0027518 | 1 | 64488649 | *RRAGD* (NM_001243623) | 289310 | 2 |  | Isovalerylcarnitine (2.79E-08), Propionylcarnitine (8.32E-10) |  |
| MARC0034307 | 1 | 76929255 |  |  | 2 |  | Isovalerylcarnitine (2.79E-08), Propionylcarnitine (8.32E-10) |  |
| MARC0050325 | 1 | 76254021 | *SIM1* (NM_001172585) | 182791 | 2 |  | Isovalerylcarnitine (2.79E-08), Propionylcarnitine (8.32E-10) |  |
| MARC0059407 | 1 | 72300056 | *FHL5* (NM_001243314) | 179132 | 2 |  | Isovalerylcarnitine (2.79E-08), Propionylcarnitine (8.32E-10) |  |
| MARC0063106 | 1 | 67392874 |  |  | 2 |  | Isovalerylcarnitine (2.79E-08), Propionylcarnitine (8.32E-10) |  |
| MARC0068954 | 1 | 69079272 |  |  | 2 |  | Isovalerylcarnitine (2.79E-08), Propionylcarnitine (8.32E-10) |  |
| MARC0075306 | 1 | 74264584 | *FBXL4* (NM_001171752) | 157849 | 2 |  | Isovalerylcarnitine (2.79E-08), Propionylcarnitine (8.32E-10) |  |
| MARC0075913 | 10 | 6389965 |  |  | 2 |  | Carnitine (1.24E-08) | Carnitine (5.24E-08) |
| MARC0080116 | 4 | 3550229 |  |  | 2 | Pyruvic acid (4.06E-08) |  | Citrulline (8.58E-07) |
| MARC0085569 | 5 | 68417760 | *CCND2* (NM_214088) | 85816 | 2 |  | Glutamic acid (3.57E-10) | Glutamic acid (4.80E-08) |
| SIRI0000655 | 1 | 78666093 |  |  | 2 |  | Isovalerylcarnitine (2.79E-08), Propionylcarnitine (8.32E-10) |  |
| WU_10.2_18_15516325 | 18 | 15516325 | *AKR1B1* (NM_001001539) | 80480 | 2 |  | Carnitine (1.24E-08) | Carnitine (5.24E-08) |
| WU_10.2_9_136780050 | 9 | 136780050 | *LAMC1* (NM_001271715) | 111754 | 2 |  | Carnitine (1.24E-08) | Carnitine (5.24E-08) |
| ALGA0038416 | 7 | 8540896 |  |  | 3 | Isovalerylcarnitine (3.47E-08), Propionylcarnitine (1.17E-09) |  | Propionylcarnitine (4.66E-07) |
| ALGA0081238 | 14 | 124416261 | *STN1* (NM_001243685) | 310170 | 3 | Isovalerylcarnitine (1.27E-08), Propionylcarnitine (3.01E-10) |  | Propionylcarnitine (3.27E-08) |
| ASGA0093565 | 6 | 135424176 | *DNAJC6* (NM_001145378) | 8^th^ intron | 3 | 1-hexadecyl-sn-glycero-3-phosphocholine (2.78E-09), 1-myristoyl-sn-glycero-3-phosphocholine (1.35E-08), LysoPC(16:0) (1.22E-07) |  |  |
| DRGA0014486 | 14 | 121391605 | *KAZALD1* (NM_001244551) | 312903 | 3 | Isovalerylcarnitine (1.27E-08), Propionylcarnitine (3.01E-10) |  | Propionylcarnitine (3.27E-08) |
| H3GA0053559 | 17 | 46932020 | *LBP* (NM_001128435) | 63443 | 3 | 1-hexadecyl-sn-glycero-3-phosphocholine (3.05E-10), 1-myristoyl-sn-glycero-3-phosphocholine (2.76E-08), LysoPC(16:0) (4.69E-09) |  |  |
| M1GA0016778 | 12 | 44192225 | *CDK5R1* (NM_001101816) | 60431 | 3 | Pyruvic acid (1.30E-09) |  | Citrulline (3.94E-07), Pyruvic acid (1.57E-08) |
| WU_10.2_14_132246191 | 14 | 132246191 | *ADRA2A* (NM_214400) | 157702 | 3 | Isovalerylcarnitine (1.27E-08), Propionylcarnitine (3.01E-10) |  | Propionylcarnitine (3.27E-08) |
| WU_10.2_6_136216429 | 6 | 136216429 | *JAK1* (NM_214114) | 298537 | 3 | 1-hexadecyl-sn-glycero-3-phosphocholine (2.78E-09), 1-myristoyl-sn-glycero-3-phosphocholine (1.35E-08), LysoPC(16:0) (1.22E-07) |  |  |
| WU_10.2_6_136863547 | 6 | 136863547 | *PGM1* (NM_001246318) | 307725 | 3 | 1-hexadecyl-sn-glycero-3-phosphocholine (2.78E-09), 1-myristoyl-sn-glycero-3-phosphocholine (1.35E-08), LysoPC(16:0) (1.22E-07) |  |  |
| WU_10.2_6_136876717 | 6 | 136876717 | *PGM1* (NM_001246318) | 294555 | 3 | 1-hexadecyl-sn-glycero-3-phosphocholine (2.78E-09), 1-myristoyl-sn-glycero-3-phosphocholine (1.35E-08), LysoPC(16:0) (1.22E-07) |  |  |
| WU_10.2_6_136972846 | 6 | 136972846 | *PGM1* (NM_001246318) | 198426 | 3 | 1-hexadecyl-sn-glycero-3-phosphocholine (2.78E-09), 1-myristoyl-sn-glycero-3-phosphocholine (1.35E-08), LysoPC(16:0) (1.22E-07) |  |  |
| ALGA0099866 | X | 105147853 | *ACSL4* (NM_001038694) | 228041 | 4 | 1-hexadecyl-sn-glycero-3-phosphocholine (3.35E-10), 1-myristoyl-sn-glycero-3-phosphocholine (2.77E-09), LysoPC(16:0) (1.16E-07), Pyruvic acid (8.03E-09) |  |  |
| ASGA0018324 | 4 | 12174119 |  |  | 4 | Citrulline (2.75E-09), Pyruvic acid (1.24E-16) |  | Citrulline (4.22E-11), Pyruvic acid (1.64E-13) |
| ASGA0081223 | X | 103657659 |  |  | 4 | 1-hexadecyl-sn-glycero-3-phosphocholine (2.63E-15), 1-myristoyl-sn-glycero-3-phosphocholine (1.49E-13), LysoPC(16:0) (7.03E-11) |  | 1-hexadecyl-sn-glycero-3-phosphocholine (4.05E-09) |
| INRA0003881 | 1 | 126310419 | *AQP9* (NM_001112684) | 357302 | 4 | 1-hexadecyl-sn-glycero-3-phosphocholine (2.63E-15), 1-myristoyl-sn-glycero-3-phosphocholine (1.49E-13), LysoPC(16:0) (7.03E-11) |  | 1-hexadecyl-sn-glycero-3-phosphocholine (4.05E-09) |
| MARC0046138 | 1 | 228287595 |  |  | 4 | 1-hexadecyl-sn-glycero-3-phosphocholine (2.63E-15), 1-myristoyl-sn-glycero-3-phosphocholine (1.49E-13), LysoPC(16:0) (7.03E-11) |  | 1-hexadecyl-sn-glycero-3-phosphocholine (4.05E-09) |
| WU_10.2_X_103597980 | X | 103597980 |  |  | 4 | 1-hexadecyl-sn-glycero-3-phosphocholine (2.63E-15), 1-myristoyl-sn-glycero-3-phosphocholine (1.49E-13), LysoPC(16:0) (7.03E-11) |  | 1-hexadecyl-sn-glycero-3-phosphocholine (4.05E-09) |
| WU_10.2_X_103653646 | X | 103653646 |  |  | 4 | 1-hexadecyl-sn-glycero-3-phosphocholine (2.63E-15), 1-myristoyl-sn-glycero-3-phosphocholine (1.49E-13), LysoPC(16:0) (7.03E-11) |  | 1-hexadecyl-sn-glycero-3-phosphocholine (4.05E-09) |
| WU_10.2_X_104796075 | X | 104796075 |  |  | 4 | 1-hexadecyl-sn-glycero-3-phosphocholine (2.63E-15), 1-myristoyl-sn-glycero-3-phosphocholine (1.49E-13), LysoPC(16:0) (7.03E-11) |  | 1-hexadecyl-sn-glycero-3-phosphocholine (4.05E-09) |
| WU_10.2_X_104910069 | X | 104910069 | *ACSL4* (NM_001038694) | 465825 | 4 | 1-hexadecyl-sn-glycero-3-phosphocholine (2.63E-15), 1-myristoyl-sn-glycero-3-phosphocholine (1.49E-13), LysoPC(16:0) (7.03E-11) |  | 1-hexadecyl-sn-glycero-3-phosphocholine (4.05E-09) |
| WU_10.2_X_104956283 | X | 104956283 | *ACSL4* (NM_001038694) | 419611 | 4 | 1-hexadecyl-sn-glycero-3-phosphocholine (2.63E-15), 1-myristoyl-sn-glycero-3-phosphocholine (1.49E-13), LysoPC(16:0) (7.03E-11) |  | 1-hexadecyl-sn-glycero-3-phosphocholine (4.05E-09) |
| WU_10.2_X_104980830 | X | 104980830 | *ACSL4* (NM_001038694) | 395064 | 4 | 1-hexadecyl-sn-glycero-3-phosphocholine (2.63E-15), 1-myristoyl-sn-glycero-3-phosphocholine (1.49E-13), LysoPC(16:0) (7.03E-11) |  | 1-hexadecyl-sn-glycero-3-phosphocholine (4.05E-09) |
| WU_10.2_X_105559450 | X | 105559450 | *ACSL4* (NM_001038694) | 106806 | 4 | 1-hexadecyl-sn-glycero-3-phosphocholine (3.35E-10), 1-myristoyl-sn-glycero-3-phosphocholine (2.77E-09), LysoPC(16:0) (1.16E-07), Pyruvic acid (8.03E-09) |  |  |
| WU_10.2_X_105583738 | X | 105583738 | *ACSL4* (NM_001038694) | 131094 | 4 | 1-hexadecyl-sn-glycero-3-phosphocholine (2.63E-15), 1-myristoyl-sn-glycero-3-phosphocholine (1.49E-13), LysoPC(16:0) (7.03E-11) |  | 1-hexadecyl-sn-glycero-3-phosphocholine (4.05E-09) |
| WU_10.2_X_114649203 | X | 114649203 | *IL13RA1* (NM_214341) | 262774 | 4 | Citrulline (2.75E-09), Pyruvic acid (1.24E-16) |  | Citrulline (4.22E-11), Pyruvic acid (1.64E-13) |

Supplementary table 2. All the metabolites in association with significant SNPs from first, second and combined two sampling times.

| Metabolite | First sampling time | | Second sampling time | | Combined two sampling times | | Common Significant SNP |
| --- | --- | --- | --- | --- | --- | --- | --- |
|  | Significant SNP Number | Significant SNP name | Significant SNP Number | Significant SNP name | Significant SNP Number | Significant SNP name |  |
| 1-hexadecyl-sn-glycero-3-phosphocholine | 23 | INRA0003881, MARC0046138, WU_10.2_6_135312468, ASGA0093565, WU_10.2_6_136216429, WU_10.2_6_136863547, WU_10.2_6_136876717, WU_10.2_6_136972846, MARC0060063, WU_10.2_8_142992366, DRGA0008955, ASGA0040302, H3GA0053559, WU_10.2_X_103597980, WU_10.2_X_103653646, ASGA0081223, WU_10.2_X_104796075, WU_10.2_X_104910069, WU_10.2_X_104956283, WU_10.2_X_104980830, ALGA0099866, WU_10.2_X_105559450, WU_10.2_X_105583738 | 1 | WU_10.2_X_37114281 | 10 | INRA0003881, MARC0046138, WU_10.2_X_103597980, WU_10.2_X_103653646, ASGA0081223, WU_10.2_X_104796075, WU_10.2_X_104910069, WU_10.2_X_104956283, WU_10.2_X_104980830, WU_10.2_X_105583738 | 0 |
| 1-myristoyl-sn-glycero-3-phosphocholine | 18 | INRA0003881, MARC0046138, ASGA0093565, WU_10.2_6_136216429, WU_10.2_6_136863547, WU_10.2_6_136876717, WU_10.2_6_136972846, H3GA0053559, WU_10.2_X_103597980, WU_10.2_X_103653646, ASGA0081223, WU_10.2_X_104796075, WU_10.2_X_104910069, WU_10.2_X_104956283, WU_10.2_X_104980830, ALGA0099866, WU_10.2_X_105559450, WU_10.2_X_105583738 | 2 | H3GA0015652, MARC0010841 | 0 | 0 | 0 |
| Aspartic acid | 0 | 0 | 4 | 3_118102178, H3GA0055935, ALGA0061605, 18_2566820 | 0 | 0 | 0 |
| Carnitine | 0 | 0 | 5 | ASGA0044546, WU_10.2_9_136780050, MARC0075913, MARC0083382, WU_10.2_18_15516325 | 4 | ASGA0044546, WU_10.2_9_136780050, MARC0075913, WU_10.2_18_15516325 | 0 |
| Citrulline | 2 | ASGA0018324, WU_10.2_X_114649203 | 0 | 0 | 5 | MARC0080116, ASGA0018324, MARC0040505, M1GA0016778, WU_10.2_X_114649203 | 0 |
| Cotinine | 2 | M1GA0015335, ALGA0072779 | 0 | 0 | 0 | 0 | 0 |
| Creatinine | 1 | ASGA0083287 | 0 | 0 | 0 | 0 | 0 |
| Glutamic acid | 0 | 0 | 5 | ALGA0028664, H3GA0014734, ASGA0023013, ASGA0025965, MARC0085569 | 5 | ALGA0028664, H3GA0014734, ASGA0023013, ASGA0025965, MARC0085569 | 0 |
| Guanine | 0 | 0 | 5 | WU_10.2_12_49599992, ASGA0054868, WU_10.2_12_49979240, ALGA0083708, ALGA0104701 | 0 | 0 | 0 |
| Isoleucyl proline | 0 | 0 | 11 | ALGA0000375, ALGA0043837, WU_10.2_7_100175430, ALGA0046501, WU_10.2_8_12998407, ALGA0046512, WU_10.2_9_128572226, H3GA0028115, WU_10.2_13_204594763, WU_10.2_13_204608919, WU_10.2_13_204631833 | 0 | 0 | 0 |
| Isovalerylcarnitine | 10 | H3GA0007606, M1GA0003090, WU_10.2_2_132463311, WU_10.2_2_132503341, WU_10.2_2_132570224, ALGA0038416, DRGA0014486, ALGA0081238, WU_10.2_14_132246191, ALGA0103368 | 57 | MARC0027518, DRGA0000994, MARC0063106, H3GA0001865, MARC0068954, ASGA0003182, ALGA0003891, ALGA0003900, DRGA0001072, DRGA0001073, ASGA0003194, MARC0059407, ALGA0003935, ALGA0003952, ALGA0003953, ASGA0003235, INRA0002726, MARC0075306, ALGA0003995, ALGA0004000, ALGA0004002, ALGA0004005, ALGA0004006, ALGA0004024, ALGA0004041, ALGA0004042, ALGA0004046, ALGA0004048, ASGA0003288, H3GA0001937, ALGA0004073, ALGA0004090, ASGA0003312, ASGA0003314, MARC0050325, H3GA0001949, ALGA0004093, ASGA0003317, ASGA0003315, MARC0034307, ASGA0083304, ALGA0004143, H3GA0001956, INRA0002819, INRA0002820, ALGA0004148, INRA0002823, ALGA0004169, MARC0021047, ALGA0004173, ASGA0003333, ALGA0004177, H3GA0001966, ASGA0003335, SIRI0000655, ASGA0057312, H3GA0046845 | 1 | ALGA0103368 | 0 |
| Lactic acid | 0 | 0 | 1 | WU_10.2_X_104031384 | 0 | 0 | 0 |
| LysoPC(16:0) | 21 | INRA0003881, MARC0046138, ASGA0093565, WU_10.2_6_136216429, WU_10.2_6_136863547, WU_10.2_6_136876717, WU_10.2_6_136972846, H3GA0053559, WU_10.2_17_48295280, ASGA0094610, MARC0027232, WU_10.2_X_103597980, WU_10.2_X_103653646, ASGA0081223, WU_10.2_X_104796075, WU_10.2_X_104910069, WU_10.2_X_104956283, WU_10.2_X_104980830, ALGA0099866, WU_10.2_X_105559450, WU_10.2_X_105583738 | 0 | 0 | 0 | 0 | 0 |
| Monosaccharide | 0 | 0 | 0 | 0 | 1 | ASGA0095299 | 0 |
| Nicotine amide | 0 | 0 | 1 | 2_29041727 | 0 | 0 | 0 |
| Propionylcarnitine | 4 | ALGA0038416, DRGA0014486, ALGA0081238, WU_10.2_14_132246191 | 61 | MARC0027518, DRGA0000994, MARC0063106, H3GA0001865, MARC0068954, ASGA0003182, ALGA0003891, ALGA0003900, DRGA0001072, DRGA0001073, ASGA0003194, MARC0059407, ALGA0003935, ALGA0003952, ALGA0003953, ASGA0003235, INRA0002726, MARC0075306, ALGA0003995, ALGA0004000, ALGA0004002, ALGA0004005, ALGA0004006, ALGA0004024, ALGA0004041, ALGA0004042, ALGA0004046, ALGA0004048, ASGA0003288, H3GA0001937, ALGA0004073, ALGA0004090, ASGA0003312, ASGA0003314, MARC0050325, H3GA0001949, ALGA0004093, ASGA0003317, ASGA0003315, MARC0034307, ASGA0083304, ALGA0004143, H3GA0001956, INRA0002819, INRA0002820, ALGA0004148, INRA0002823, ALGA0004169, MARC0021047, ALGA0004173, ASGA0003333, ALGA0004177, H3GA0001966, ASGA0003335, SIRI0000655, DRGA0004012, WU_10.2_10_71866847, H3GA0035928, ASGA0057312, H3GA0046845, ASGA0078611 | 4 | ALGA0038416, DRGA0014486, ALGA0081238, WU_10.2_14_132246191 |  |
| Pyruvic acid | 16 | MARC0110390, ALGA0014130, MARC0080116, ASGA0018324, ALGA0049375, ALGA0049385, MARC0001720, ASGA0042433, H3GA0026947, ASGA0042436, MARC0092124, INRA0031485, M1GA0016778, ALGA0099866, WU_10.2_X_105559450, WU_10.2_X_114649203 | 1 | MARC0023109 | 3 | ASGA0018324, M1GA0016778, WU_10.2_X_114649203 |  |
| Total | 97 |  | 154 |  | 33 |  | 0 |

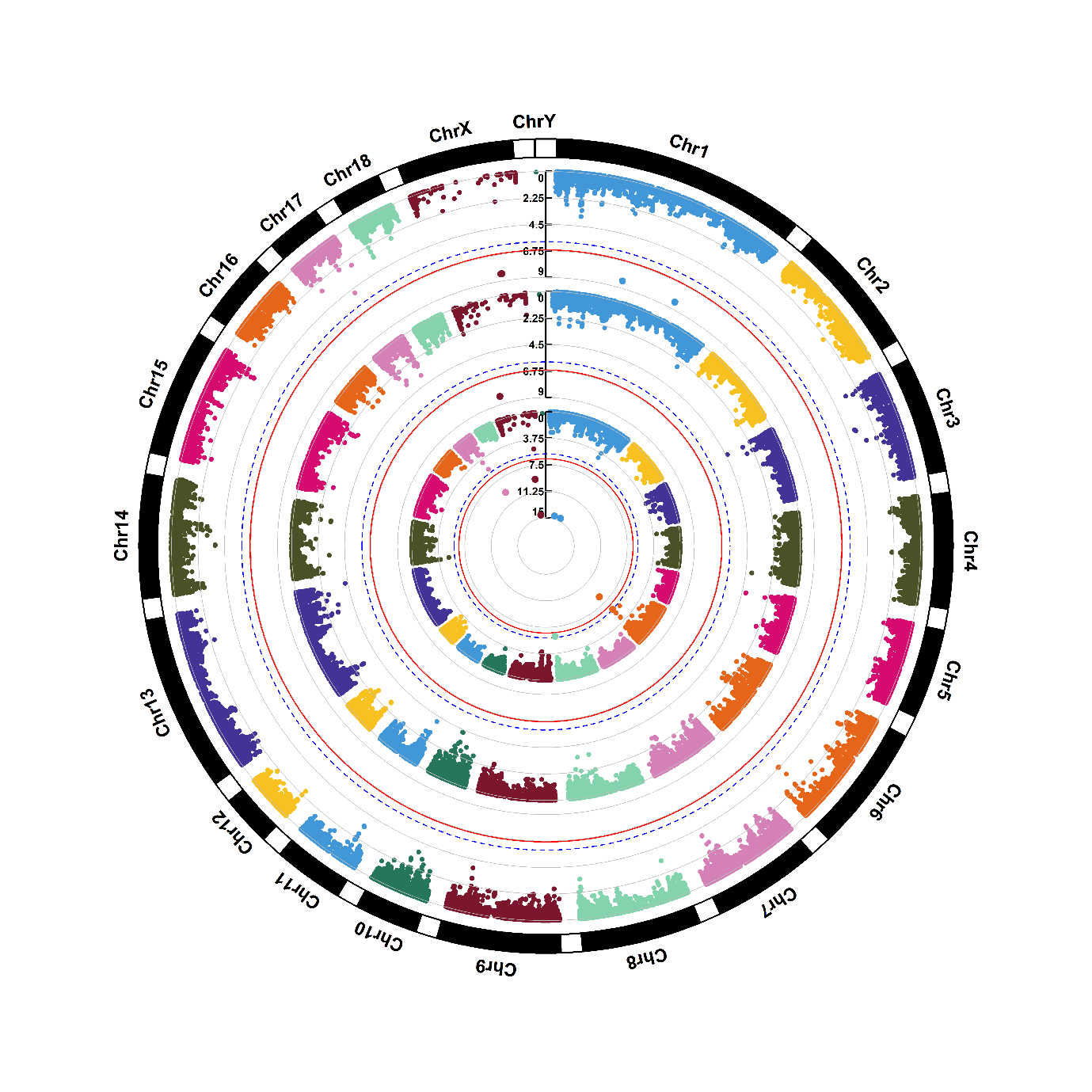

Supplementary figure 1.1. Manhattan plots of genome-wide association for 1-hexadecyl-sn-glycero-3-phosphocholine (M1). Note: Y-axis indicates the log_10_(*P*-value). Blue dotted and red solid lines indicate the genome-wide threshold of 0.05 and 0.01 after Bonferroni multiple testing, respectively. The three tracks indicate the metabolites from first sampling time, second sampling time and combined two sampling times, respectively, from outside to inside.

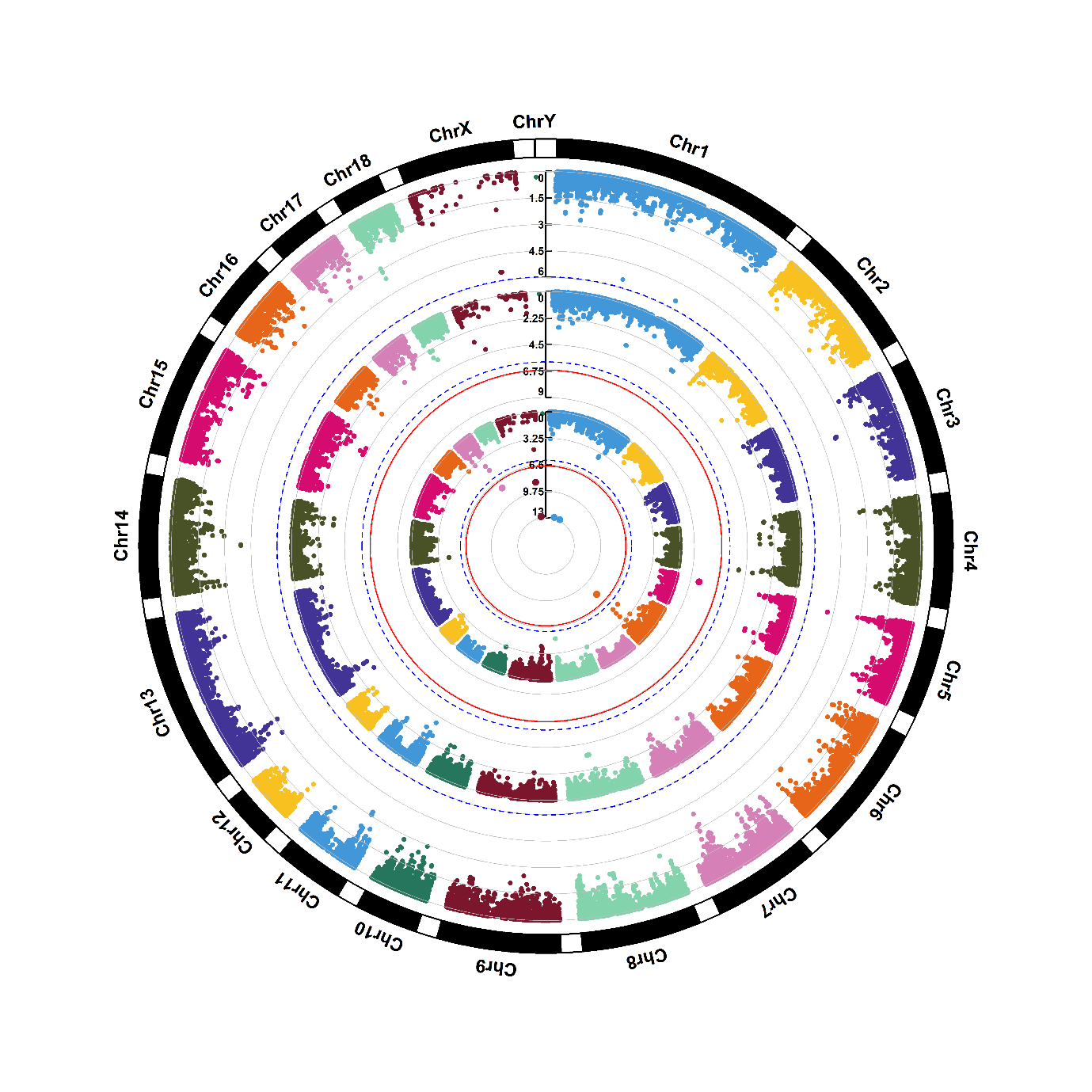

Supplementary figure 1.2. Manhattan plots of genome-wide association for 1-myristoyl-sn-glycero-3-phosphocholine (M2). Note: Y-axis indicates the log_10_(*P*-value). Blue dotted and red solid lines indicate the genome-wide threshold of 0.05 and 0.01 after Bonferroni multiple testing, respectively. The three tracks indicate the metabolites from first sampling time, second sampling time and combined two sampling times, respectively, from outside to inside.

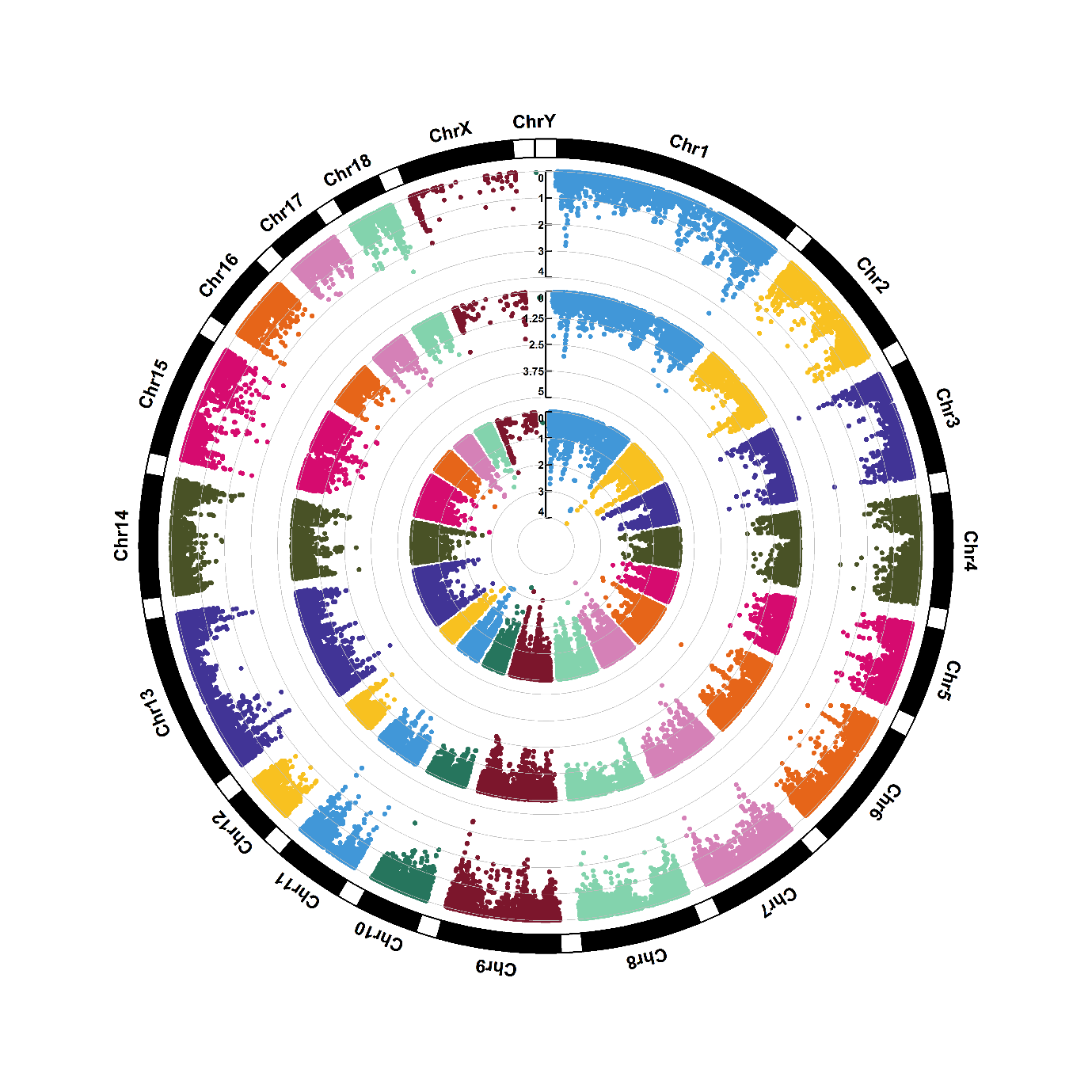

Supplementary figure 1.3. Manhattan plots of genome-wide association for trimethylammonium (M3). Note: Y-axis indicates the log_10_(*P*-value). Blue dotted and red solid lines indicate the genome-wide threshold of 0.05 and 0.01 after Bonferroni multiple testing, respectively. The three tracks indicate the metabolites from first sampling time, second sampling time and combined two sampling times, respectively, from outside to inside.

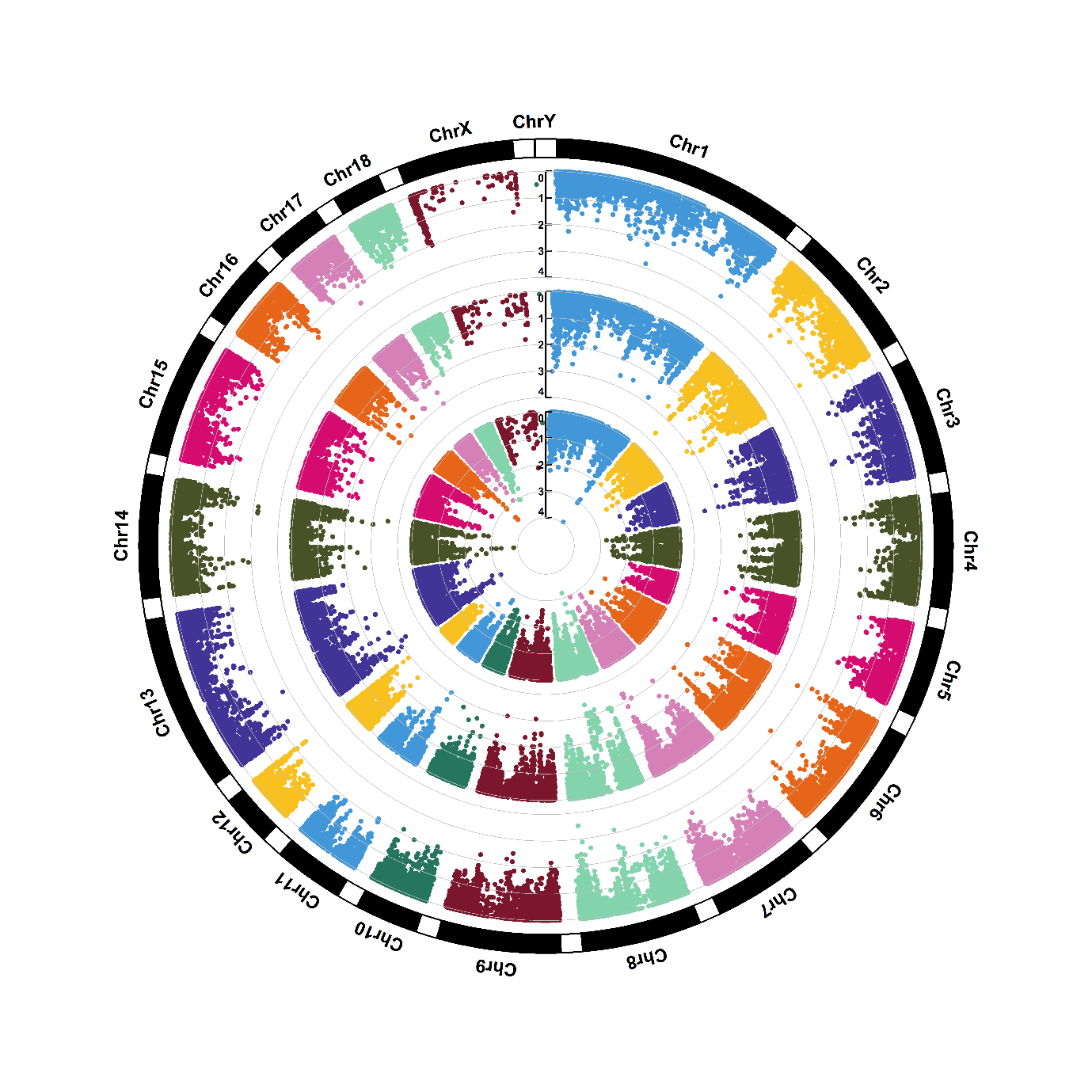

Supplementary figure 1.4. Manhattan plots of genome-wide association for 4-aminobenzoic acid (M4). Note: Y-axis indicates the log_10_(*P*-value). Blue dotted and red solid lines indicate the genome-wide threshold of 0.05 and 0.01 after Bonferroni multiple testing, respectively. The three tracks indicate the metabolites from first sampling time, second sampling time and combined two sampling times, respectively, from outside to inside.

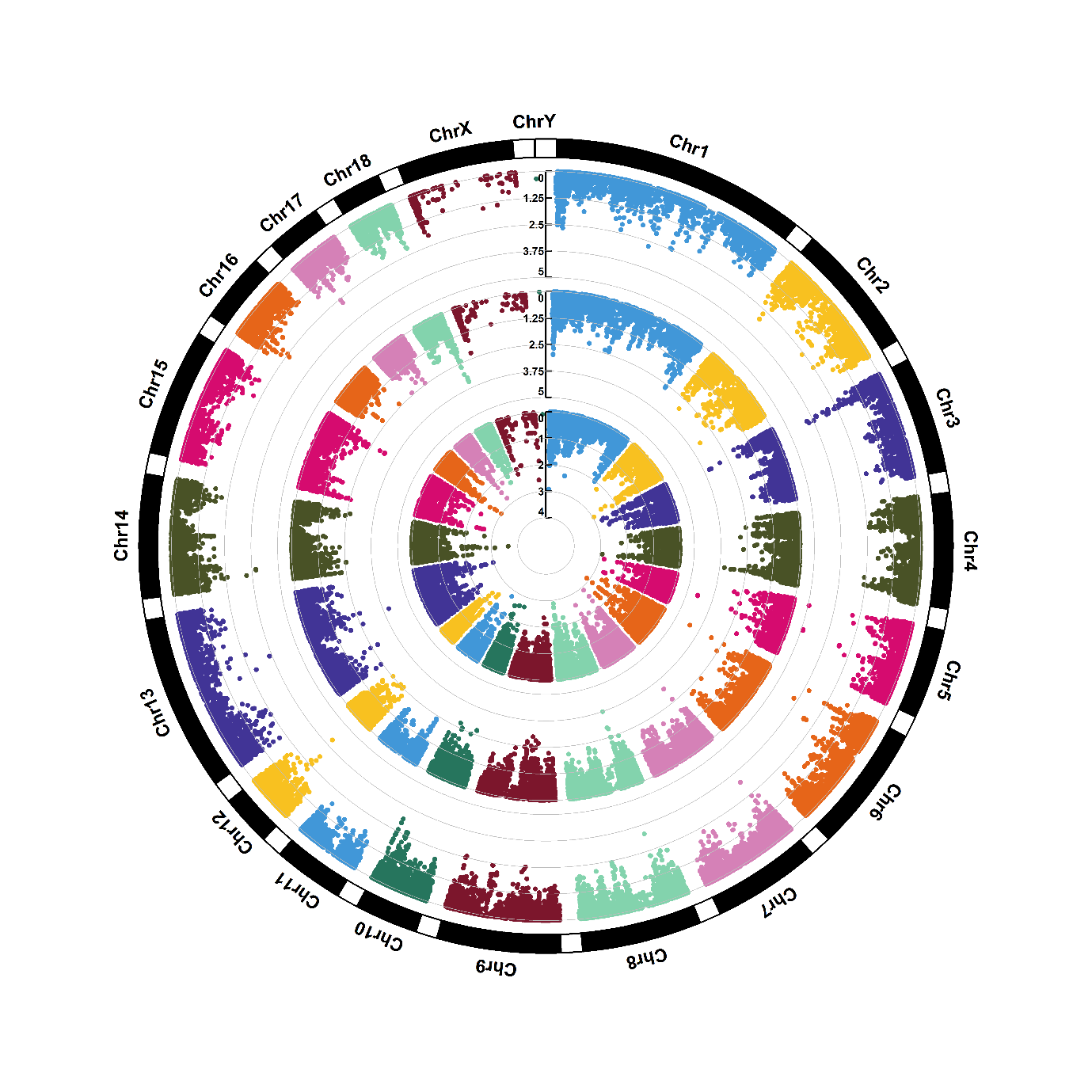

Supplementary figure 1.5. Manhattan plots of genome-wide association for 5-methyl-5,6-dihydrouracils (M5). Note: Y-axis indicates the log_10_(*P*-value). Blue dotted and red solid lines indicate the genome-wide threshold of 0.05 and 0.01 after Bonferroni multiple testing, respectively. The three tracks indicate the metabolites from first sampling time, second sampling time and combined two sampling times, respectively, from outside to inside.

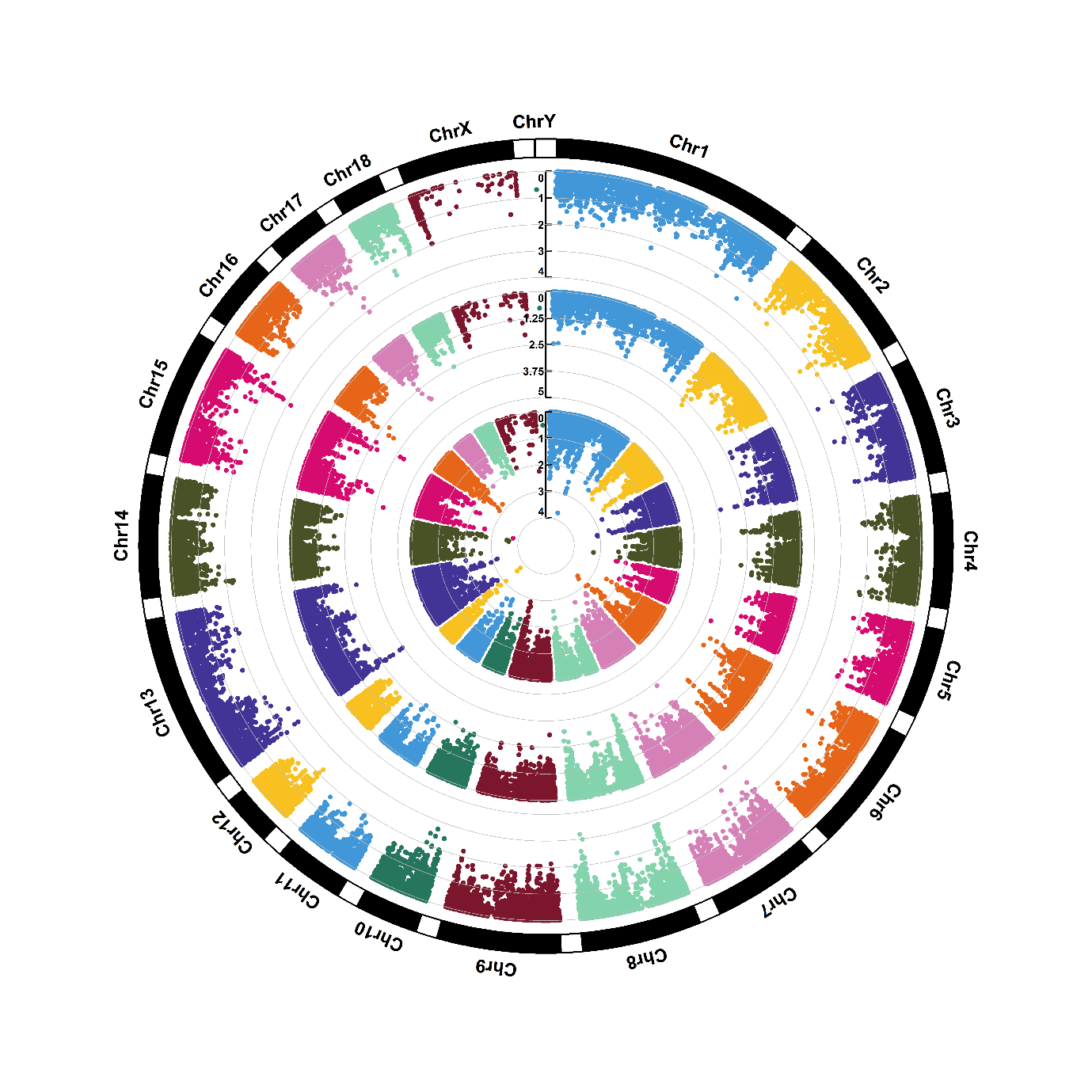

Supplementary figure 1.6. Manhattan plots of genome-wide association for acetaminophen (M6). Note: Y-axis indicates the log_10_(*P*-value). Blue dotted and red solid lines indicate the genome-wide threshold of 0.05 and 0.01 after Bonferroni multiple testing, respectively. The three tracks indicate the metabolites from first sampling time, second sampling time and combined two sampling times, respectively, from outside to inside.

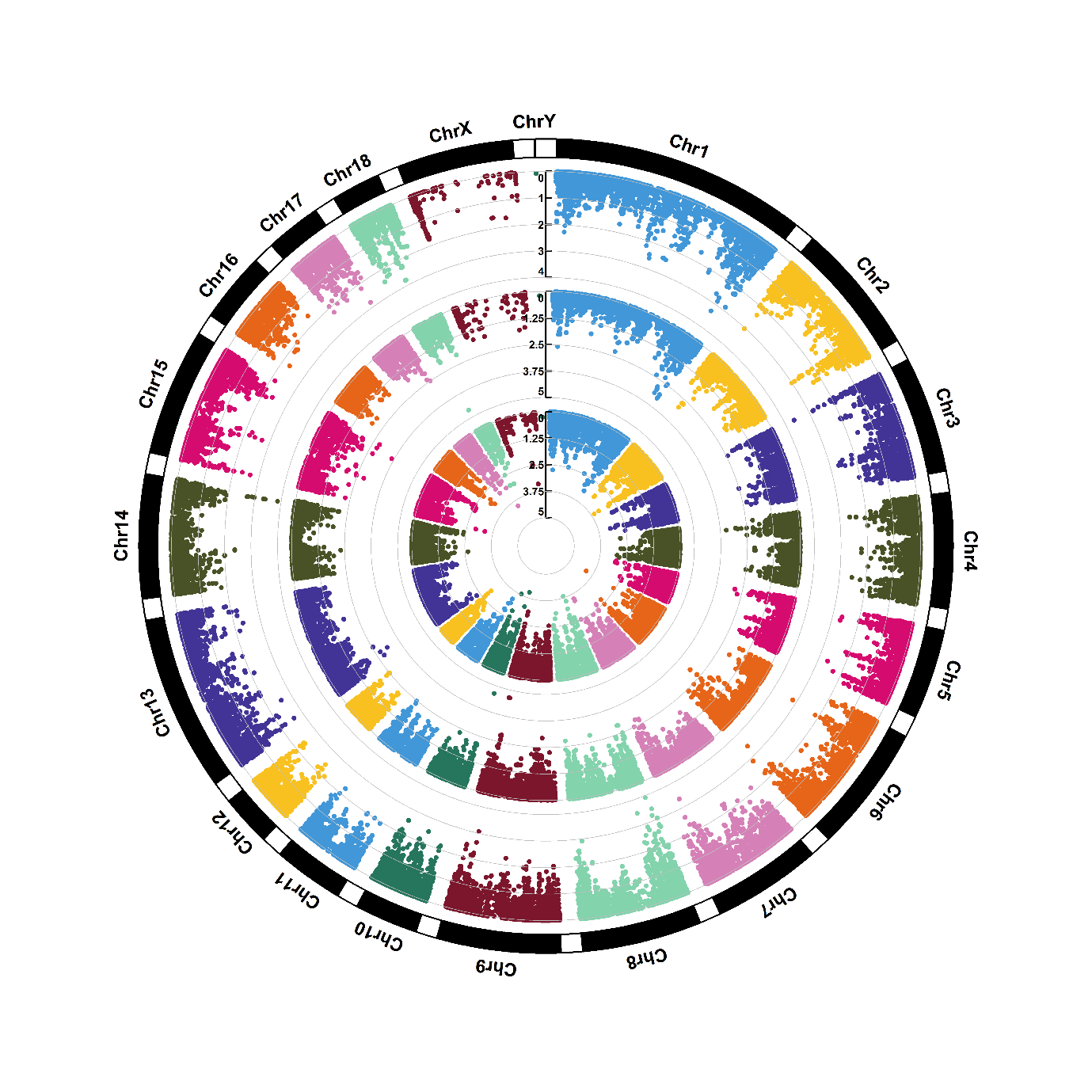

Supplementary figure 1.7. Manhattan plots of genome-wide association for acetylcarnitine (M7). Note: Y-axis indicates the log_10_(*P*-value). Blue dotted and red solid lines indicate the genome-wide threshold of 0.05 and 0.01 after Bonferroni multiple testing, respectively. The three tracks indicate the metabolites from first sampling time, second sampling time and combined two sampling times, respectively, from outside to inside.

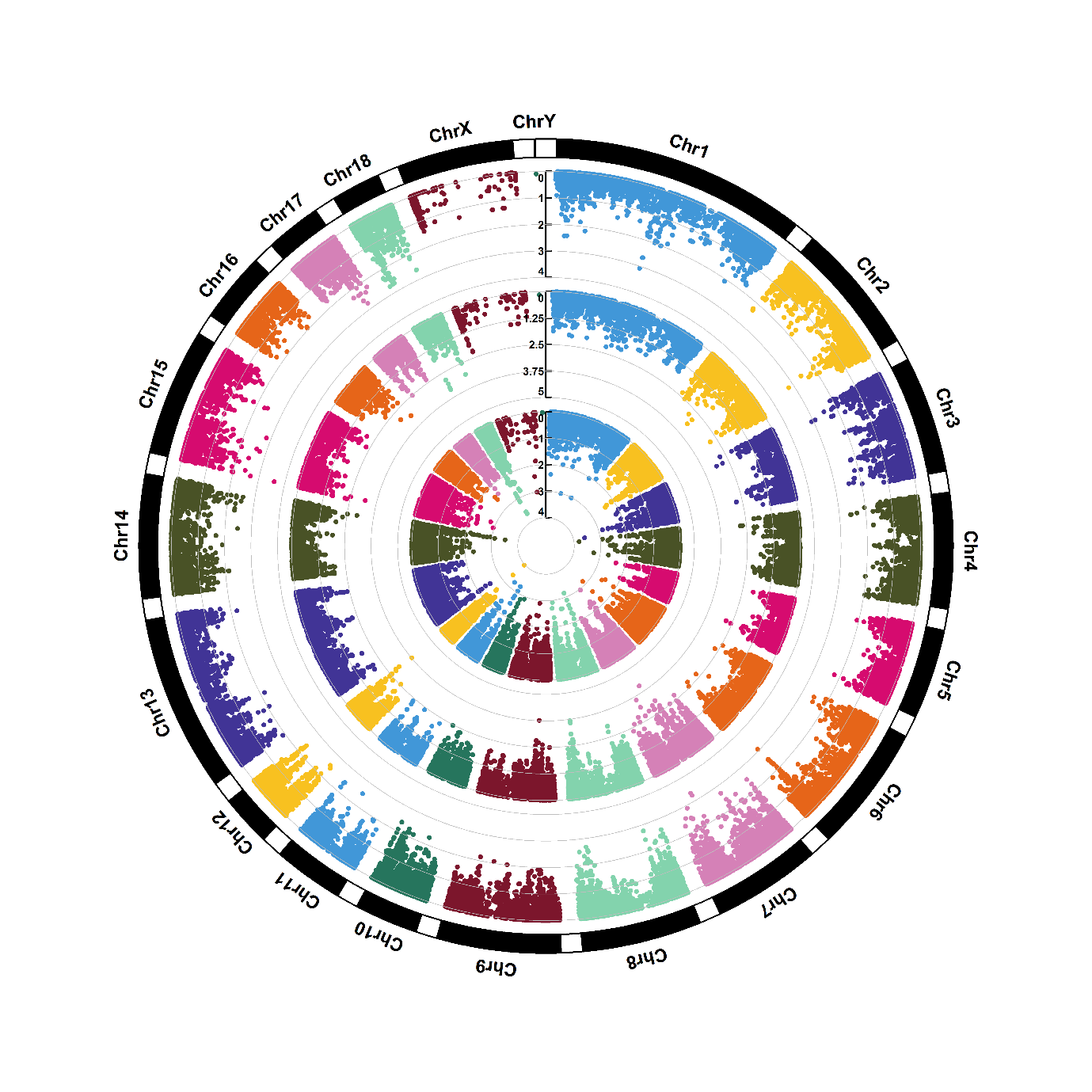

Supplementary figure 1.8. Manhattan plots of genome-wide association for alanine (M8). Note: Y-axis indicates the log_10_(*P*-value). Blue dotted and red solid lines indicate the genome-wide threshold of 0.05 and 0.01 after Bonferroni multiple testing, respectively. The three tracks indicate the metabolites from first sampling time, second sampling time and combined two sampling times, respectively, from outside to inside.

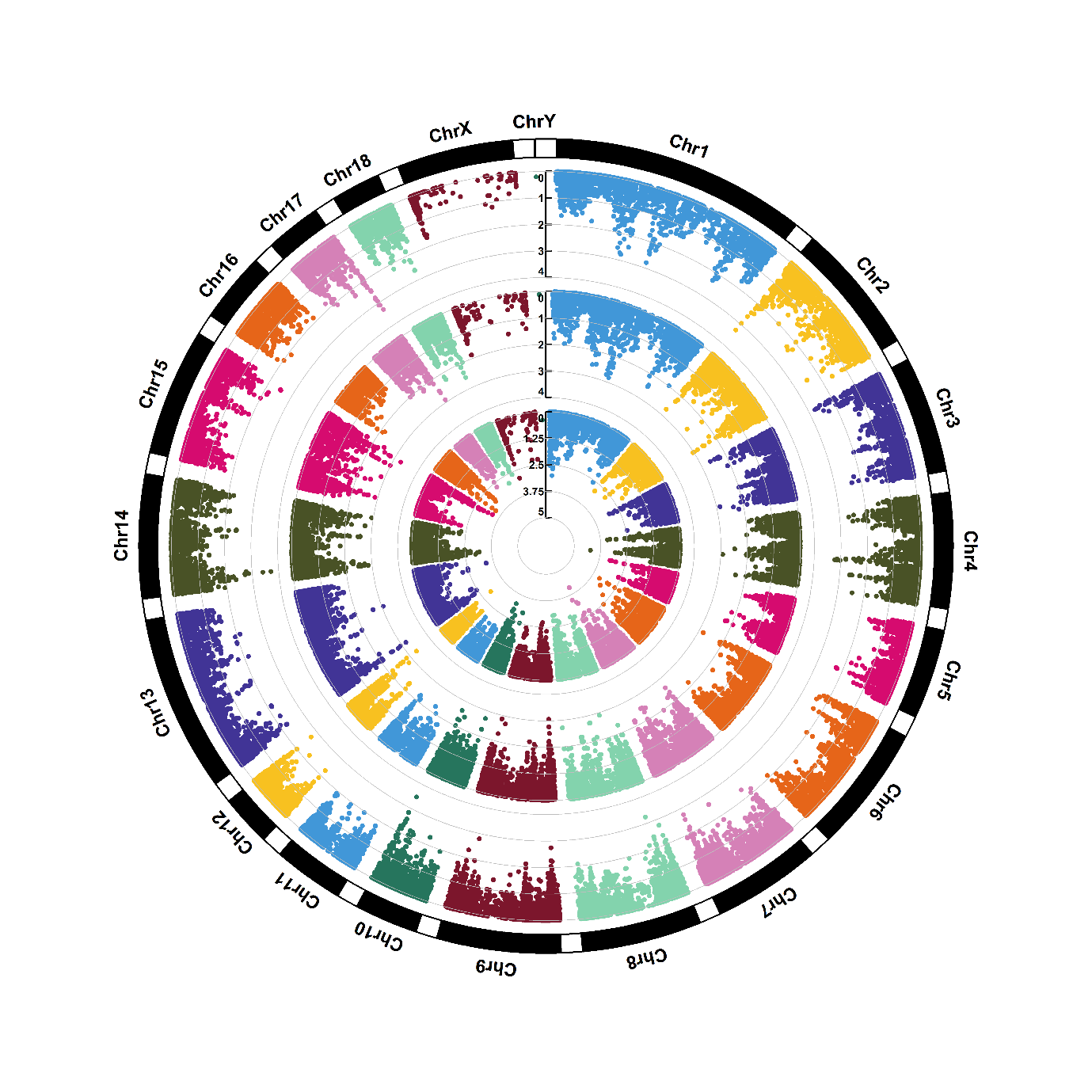

Supplementary figure 1.9. Manhattan plots of genome-wide association for arginine (M9). Note: Y-axis indicates the log_10_(*P*-value). Blue dotted and red solid lines indicate the genome-wide threshold of 0.05 and 0.01 after Bonferroni multiple testing, respectively. The three tracks indicate the metabolites from first sampling time, second sampling time and combined two sampling times, respectively, from outside to inside.

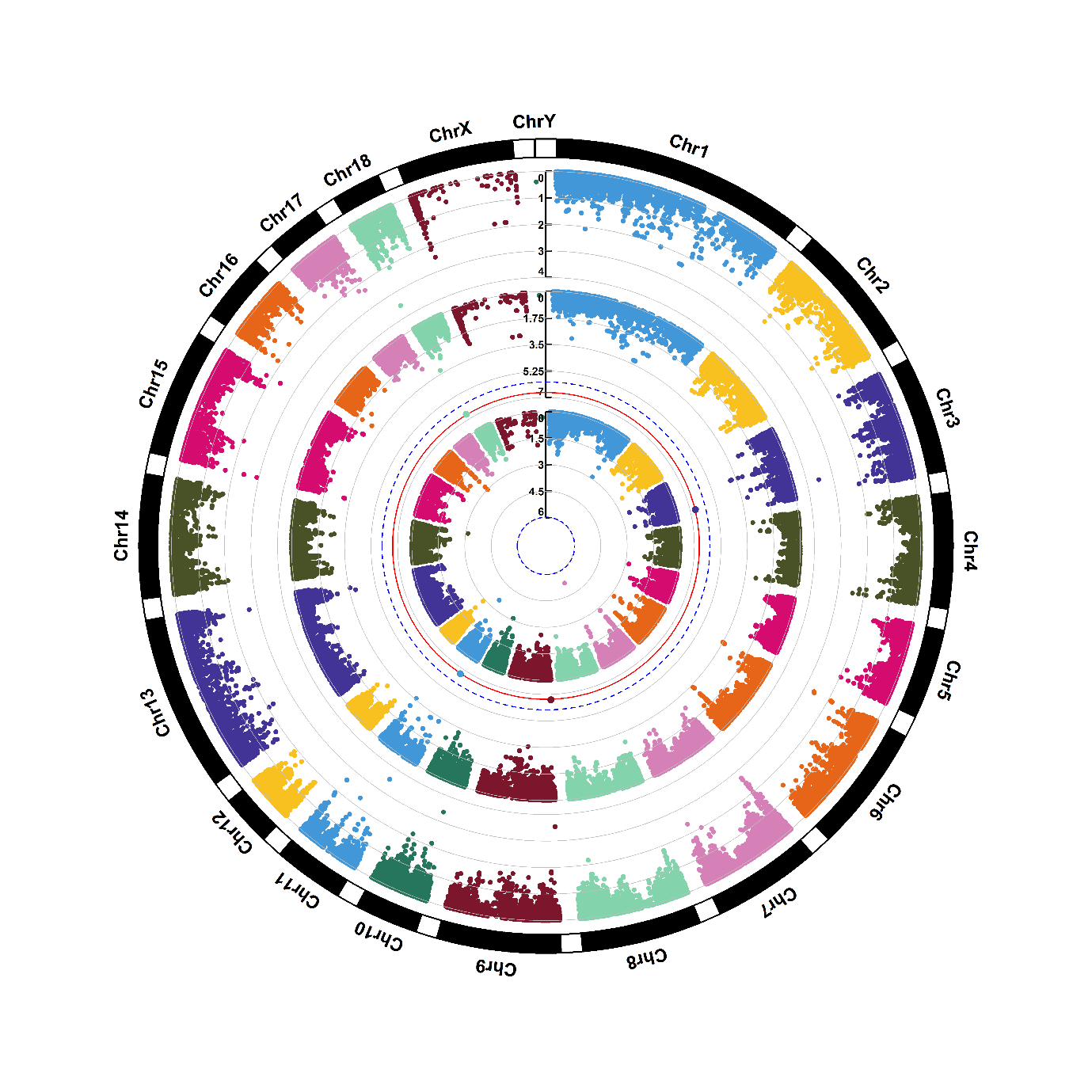

Supplementary figure 1.10. Manhattan plots of genome-wide association for aspartic acid (M10). Note: Y-axis indicates the log_10_(*P*-value). Blue dotted and red solid lines indicate the genome-wide threshold of 0.05 and 0.01 after Bonferroni multiple testing, respectively. The three tracks indicate the metabolites from first sampling time, second sampling time and combined two sampling times, respectively, from outside to inside.

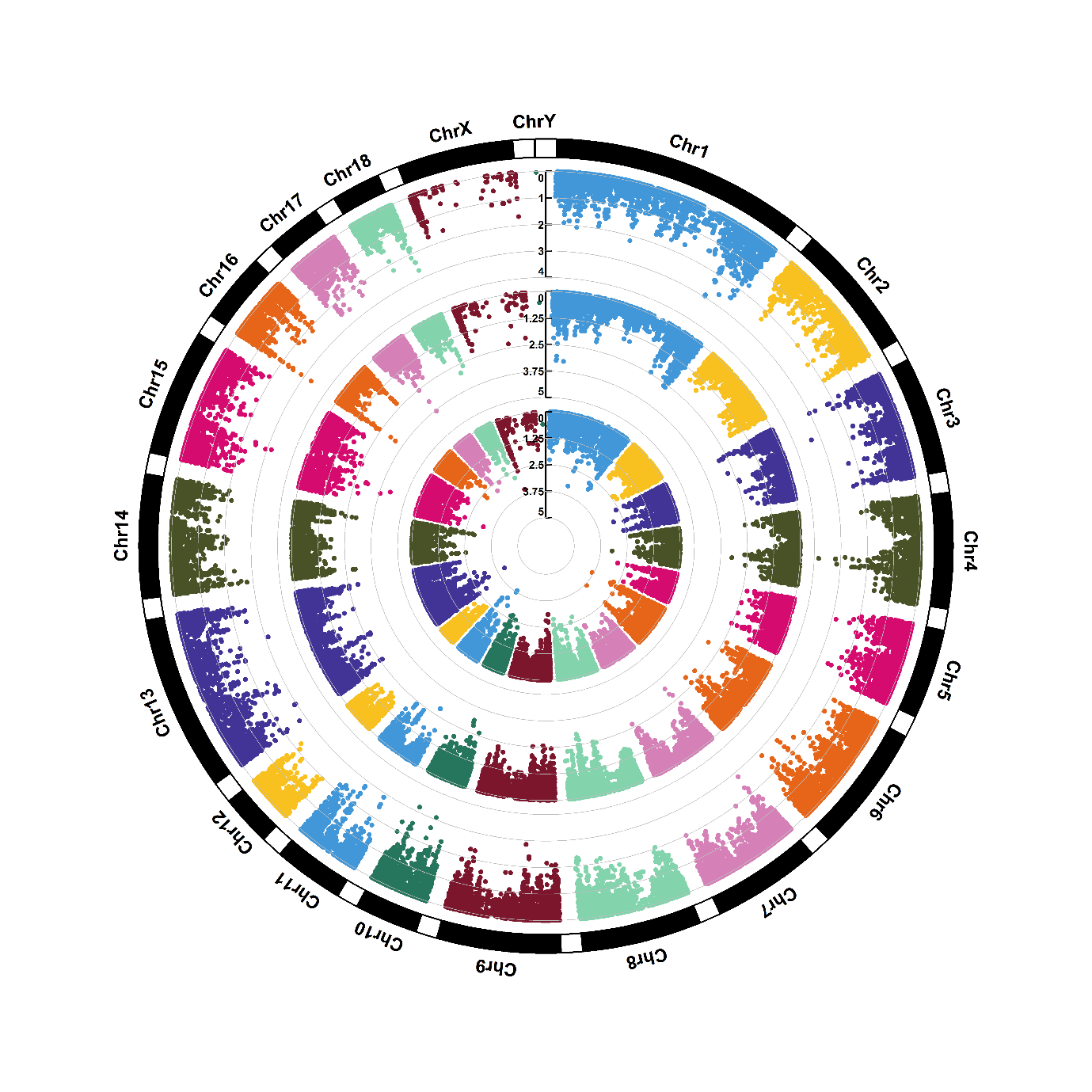

Supplementary figure 1.11. Manhattan plots of genome-wide association for benzoic acid (M11). Note: Y-axis indicates the log_10_(*P*-value). Blue dotted and red solid lines indicate the genome-wide threshold of 0.05 and 0.01 after Bonferroni multiple testing, respectively. The three tracks indicate the metabolites from first sampling time, second sampling time and combined two sampling times, respectively, from outside to inside.

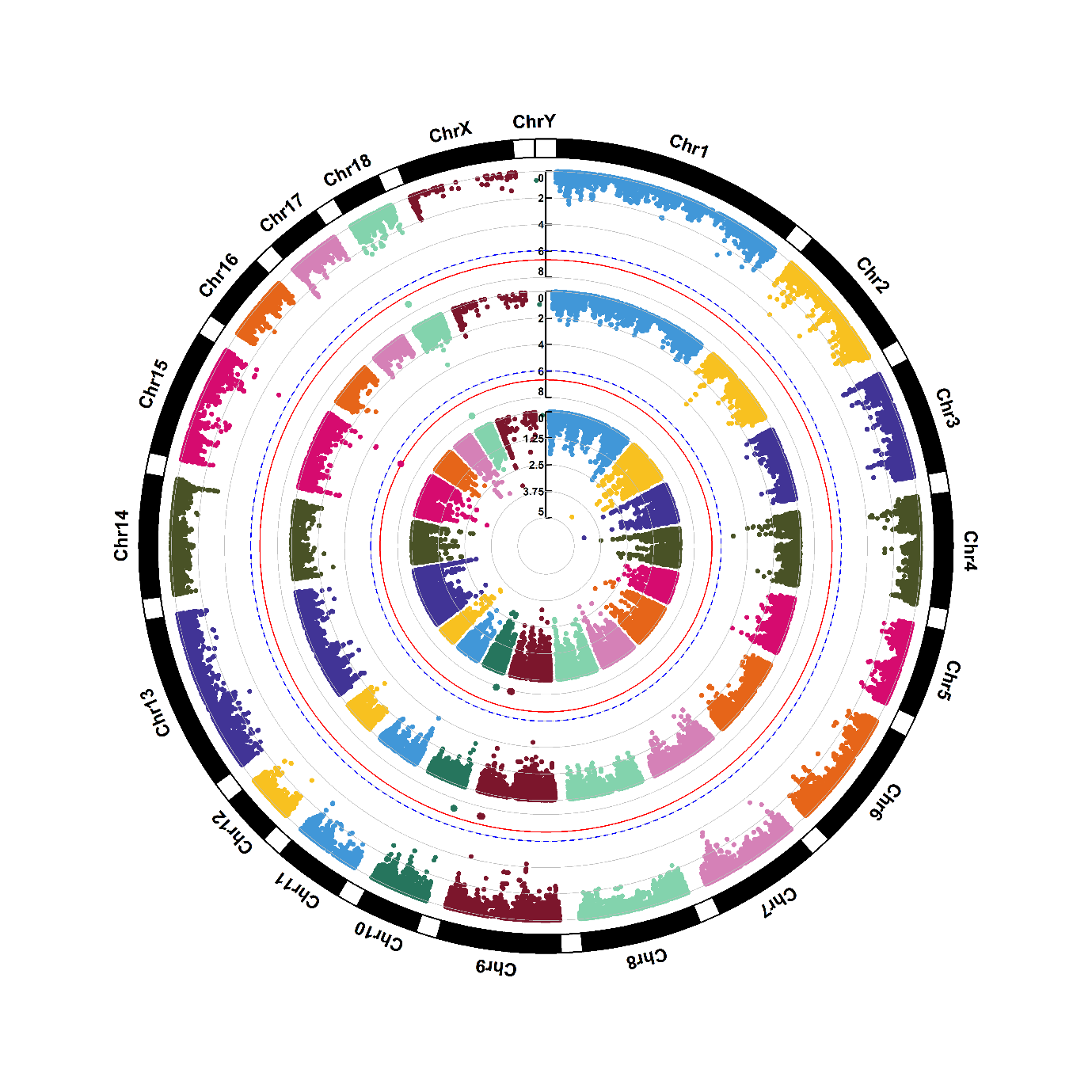

Supplementary figure 1.12. Manhattan plots of genome-wide association for carnitine (M12). Note: Y-axis indicates the log_10_(*P*-value). Blue dotted and red solid lines indicate the genome-wide threshold of 0.05 and 0.01 after Bonferroni multiple testing, respectively. The three tracks indicate the metabolites from first sampling time, second sampling time and combined two sampling times, respectively, from outside to inside.

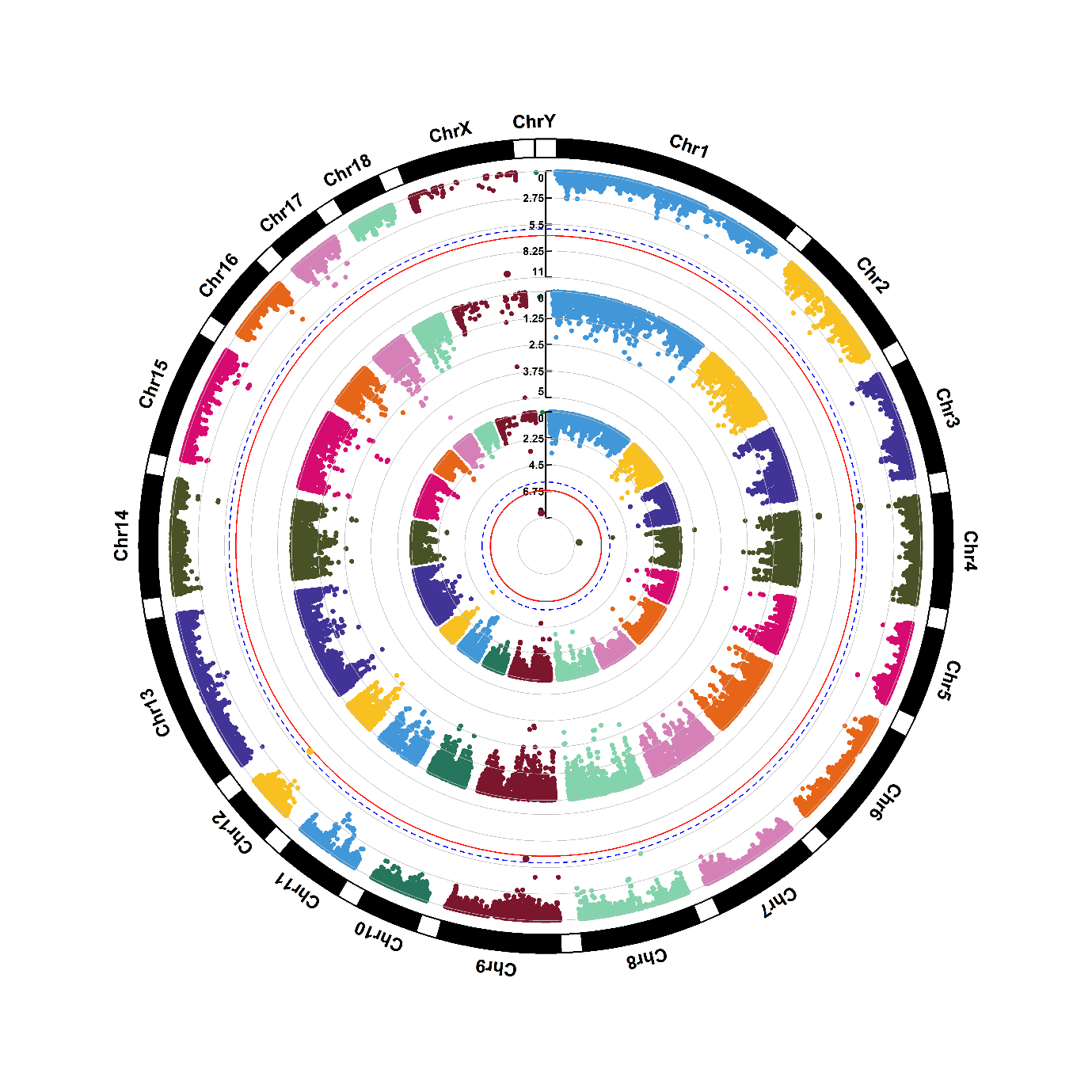

Supplementary figure 1.13. Manhattan plots of genome-wide association for citrulline (M13). Note: Y-axis indicates the log_10_(*P*-value). Blue dotted and red solid lines indicate the genome-wide threshold of 0.05 and 0.01 after Bonferroni multiple testing, respectively. The three tracks indicate the metabolites from first sampling time, second sampling time and combined two sampling times, respectively, from outside to inside.

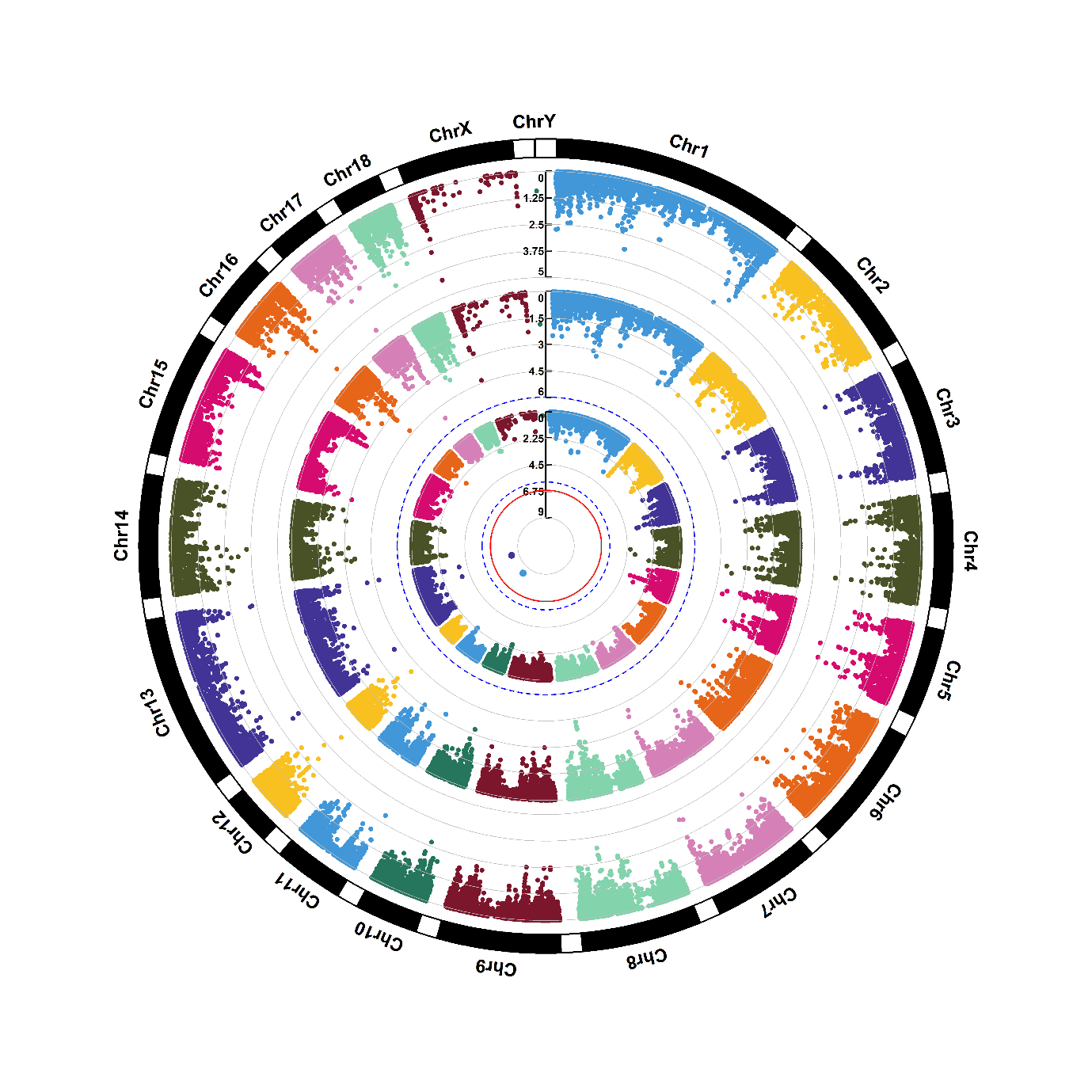

Supplementary figure 1.14. Manhattan plots of genome-wide association for cotinine (M14). Note: Y-axis indicates the log_10_(*P*-value). Blue dotted and red solid lines indicate the genome-wide threshold of 0.05 and 0.01 after Bonferroni multiple testing, respectively. The three tracks indicate the metabolites from first sampling time, second sampling time and combined two sampling times, respectively, from outside to inside.

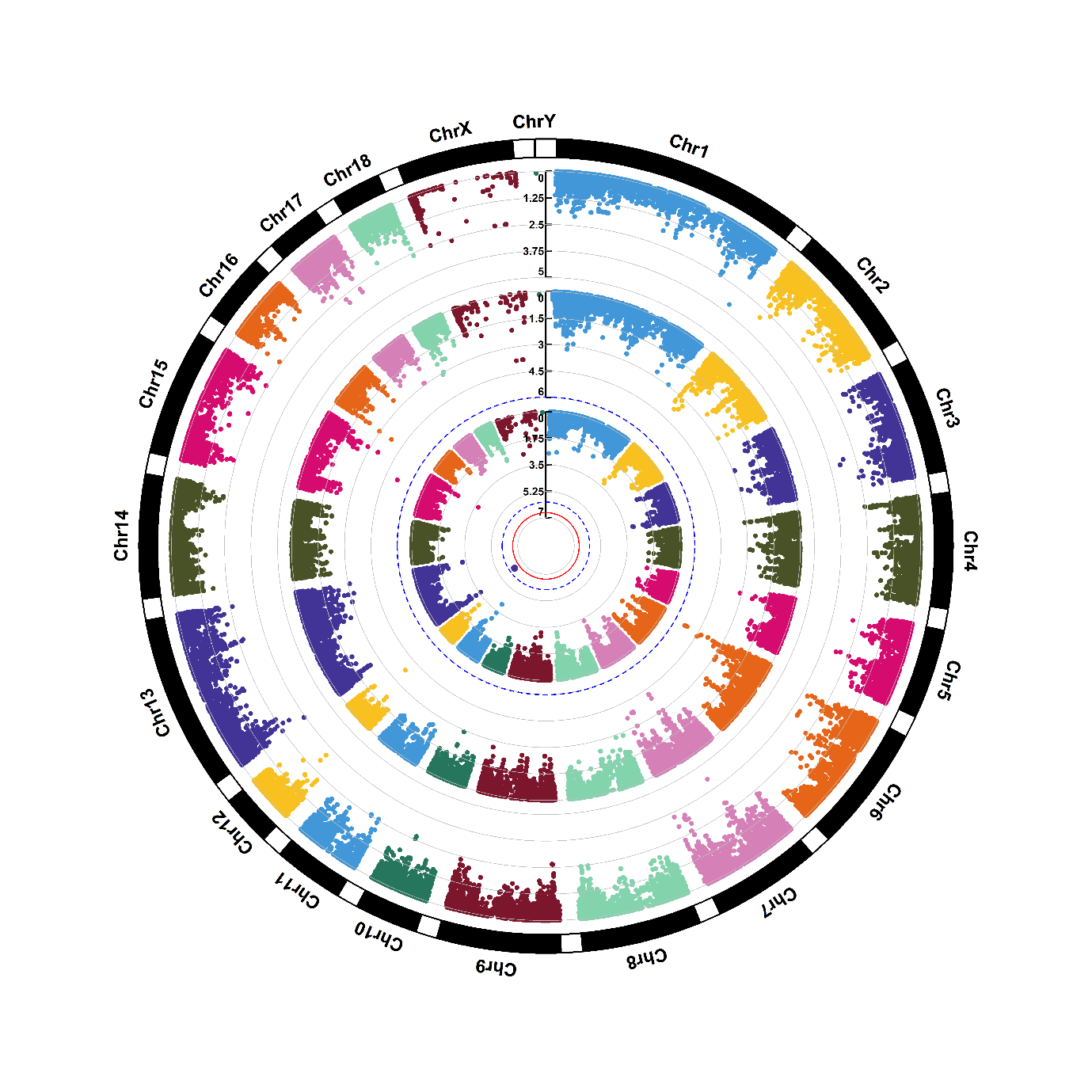

Supplementary figure 1.15. Manhattan plots of genome-wide association for creatinine (M15). Note: Y-axis indicates the log_10_(*P*-value). Blue dotted and red solid lines indicate the genome-wide threshold of 0.05 and 0.01 after Bonferroni multiple testing, respectively. The three tracks indicate the metabolites from first sampling time, second sampling time and combined two sampling times, respectively, from outside to inside.

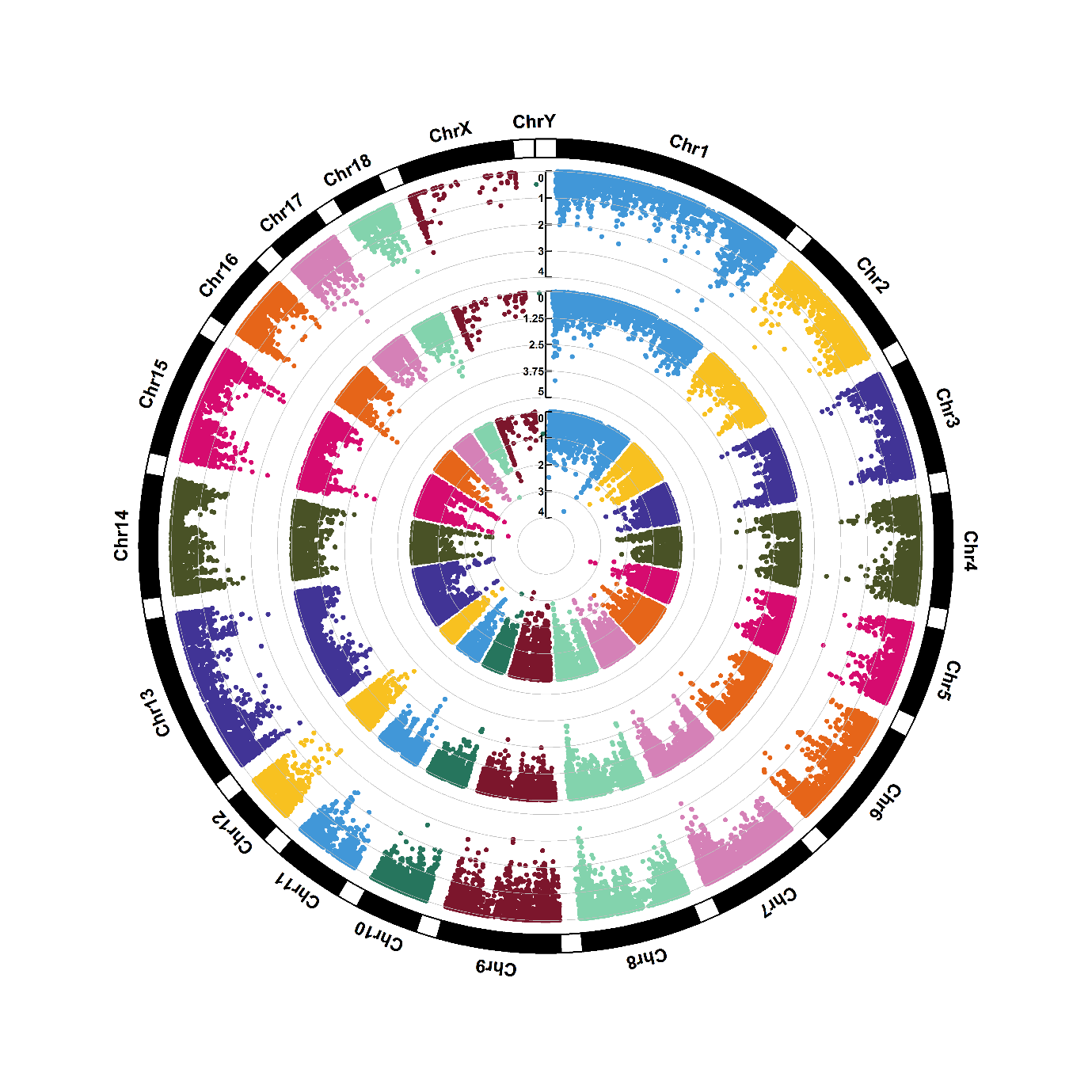

Supplementary figure 1.16. Manhattan plots of genome-wide association for cytidine (M16). Note: Y-axis indicates the log_10_(*P*-value). Blue dotted and red solid lines indicate the genome-wide threshold of 0.05 and 0.01 after Bonferroni multiple testing, respectively. The three tracks indicate the metabolites from first sampling time, second sampling time and combined two sampling times, respectively, from outside to inside.

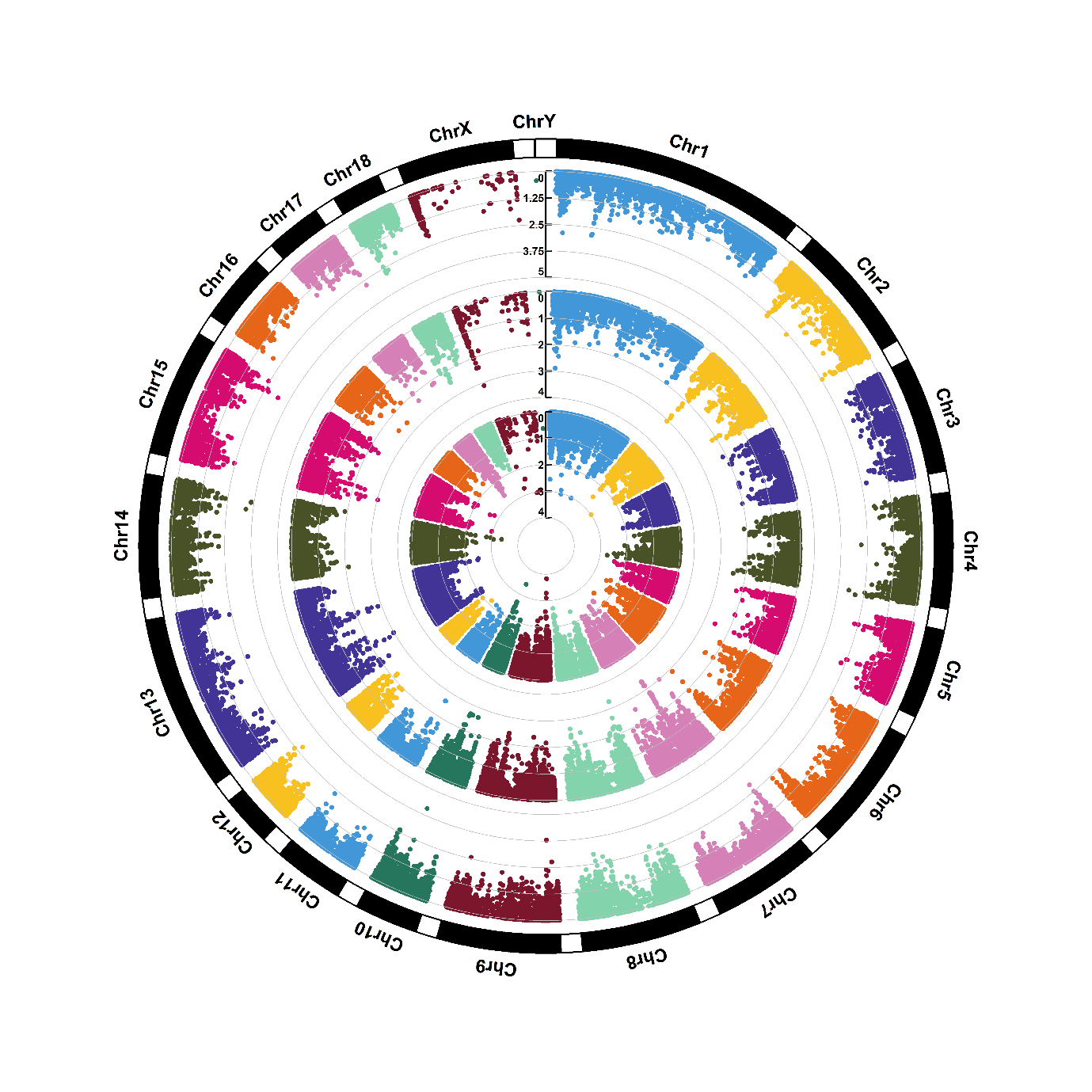

Supplementary figure 1.17. Manhattan plots of genome-wide association for disaccharide (M17). Note: Y-axis indicates the log_10_(*P*-value). Blue dotted and red solid lines indicate the genome-wide threshold of 0.05 and 0.01 after Bonferroni multiple testing, respectively. The three tracks indicate the metabolites from first sampling time, second sampling time and combined two sampling times, respectively, from outside to inside.

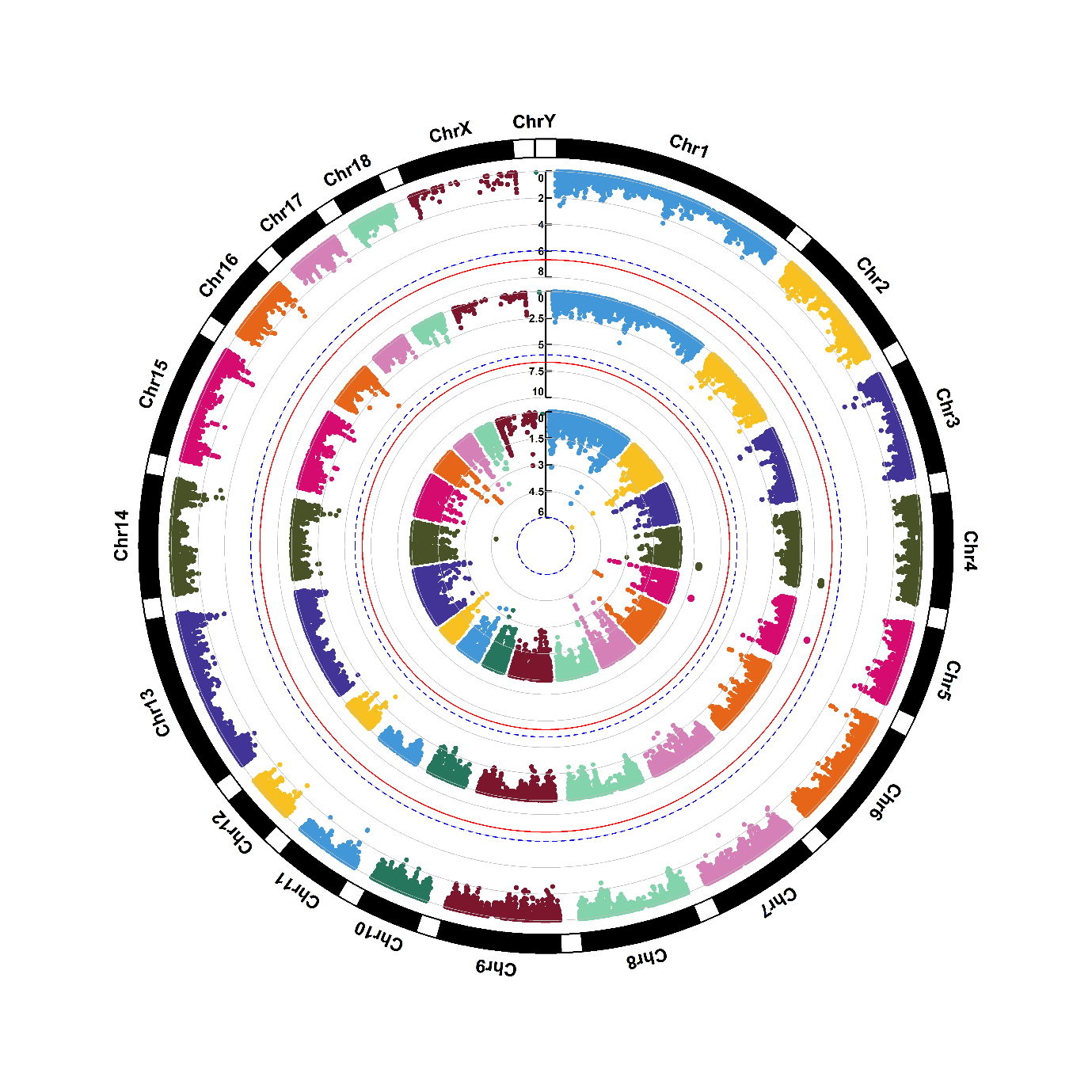

Supplementary figure 1.18. Manhattan plots of genome-wide association for glutamic acid (M18). Note: Y-axis indicates the log_10_(*P*-value). Blue dotted and red solid lines indicate the genome-wide threshold of 0.05 and 0.01 after Bonferroni multiple testing, respectively. The three tracks indicate the metabolites from first sampling time, second sampling time and combined two sampling times, respectively, from outside to inside.

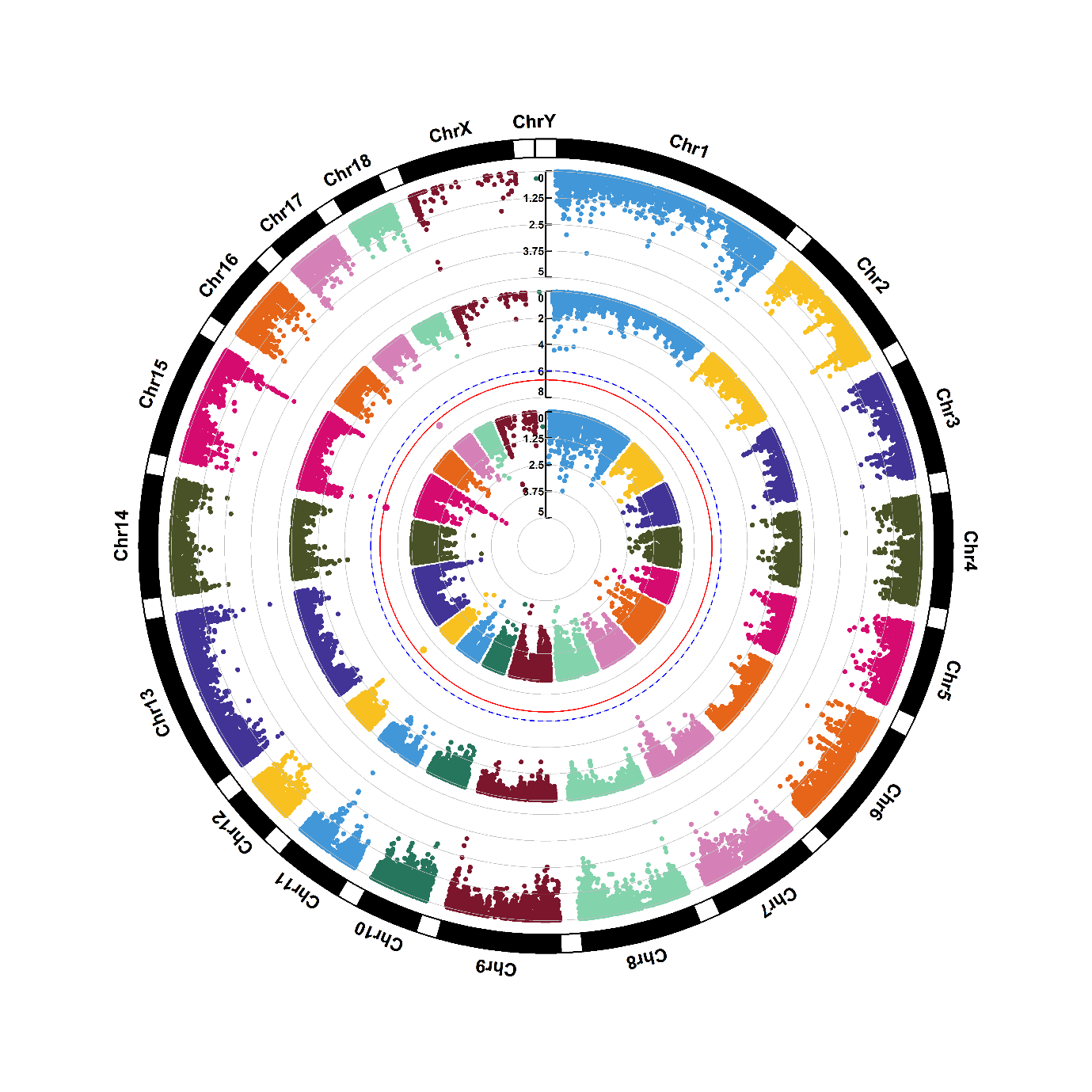

Supplementary figure 1.19. Manhattan plots of genome-wide association for guanine (M19). Note: Y-axis indicates the log_10_(*P*-value). Blue dotted and red solid lines indicate the genome-wide threshold of 0.05 and 0.01 after Bonferroni multiple testing, respectively. The three tracks indicate the metabolites from first sampling time, second sampling time and combined two sampling times, respectively, from outside to inside.

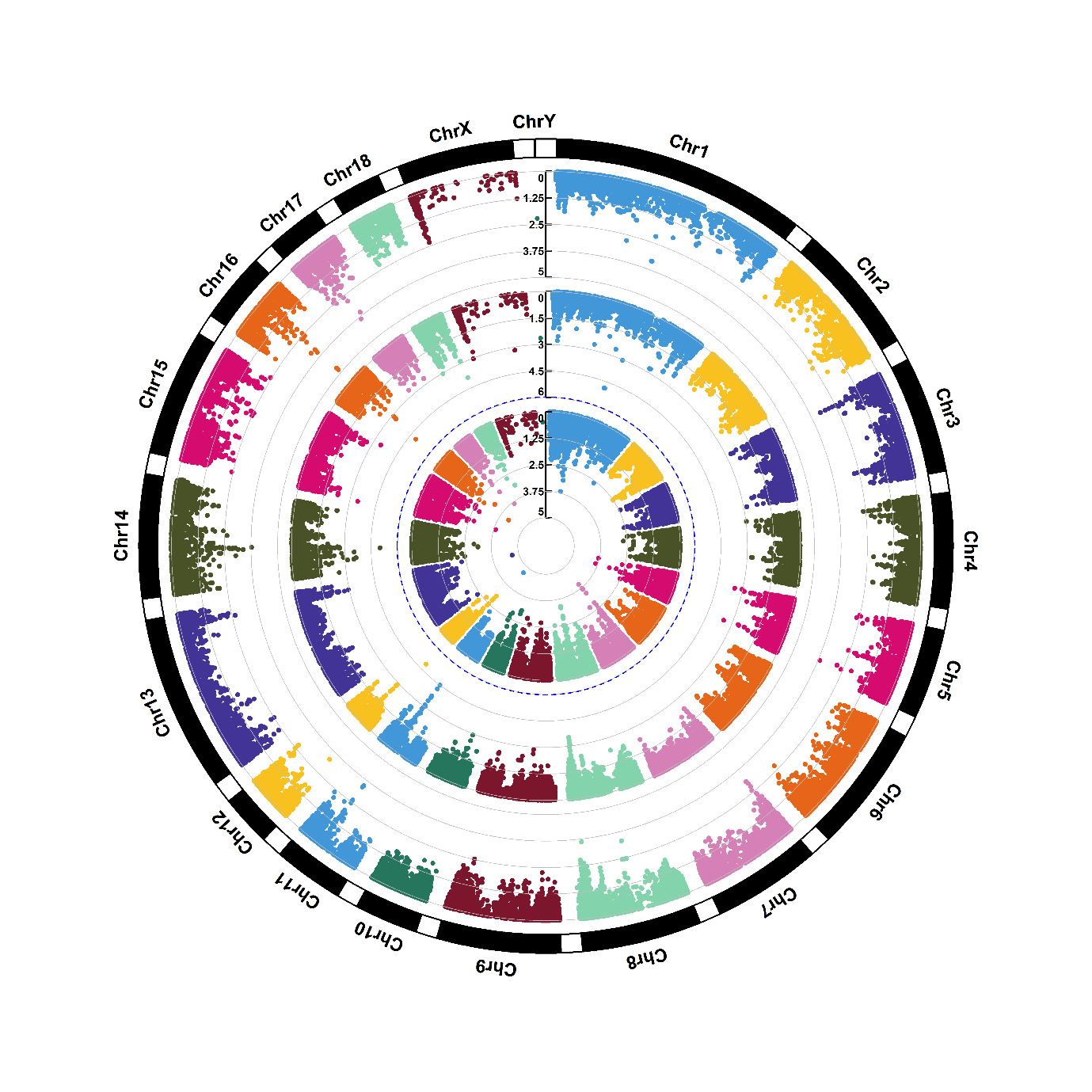

Supplementary figure 1.20. Manhattan plots of genome-wide association for guanosine (M20). Note: Y-axis indicates the log_10_(*P*-value). Blue dotted and red solid lines indicate the genome-wide threshold of 0.05 and 0.01 after Bonferroni multiple testing, respectively. The three tracks indicate the metabolites from first sampling time, second sampling time and combined two sampling times, respectively, from outside to inside.

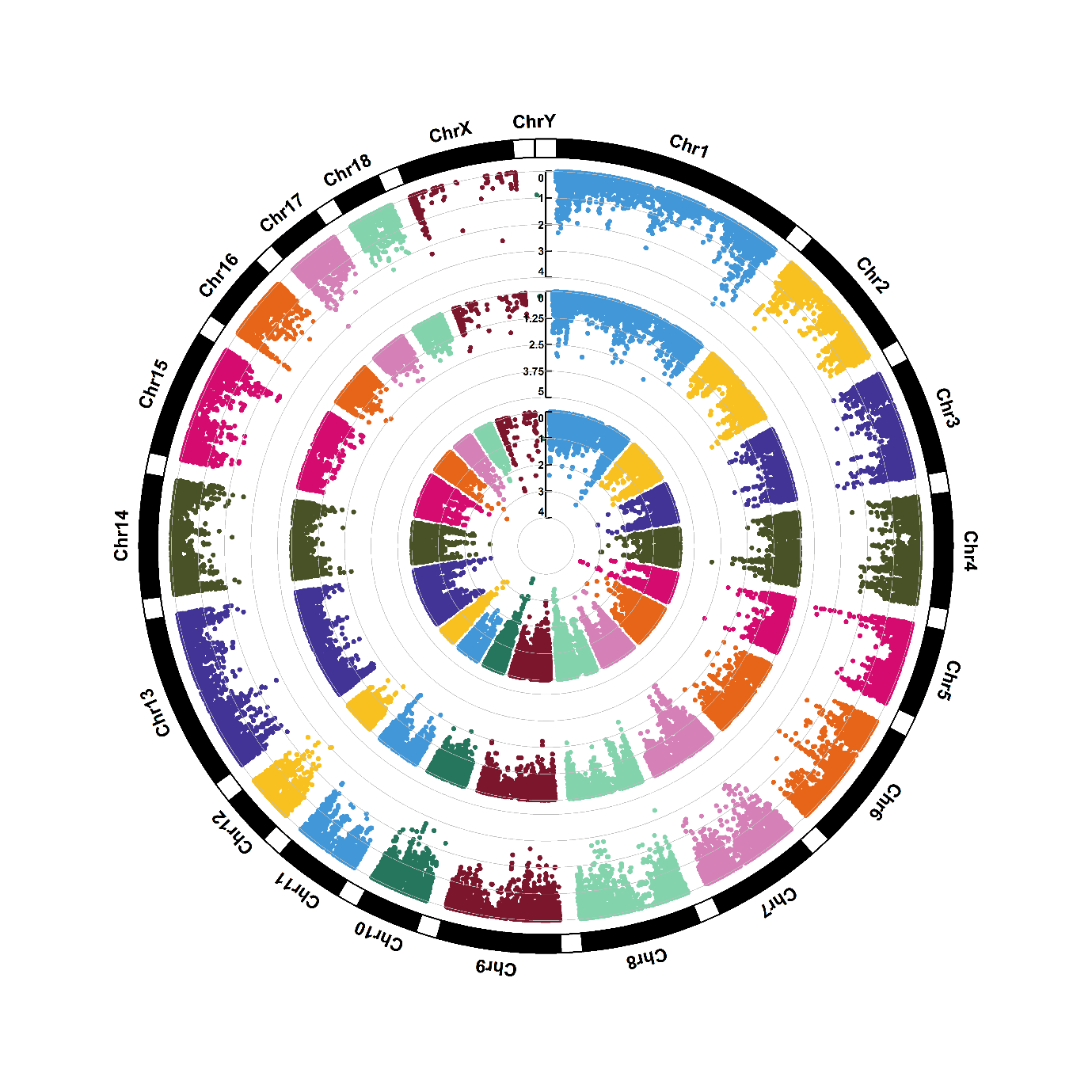

Supplementary figure 1.21. Manhattan plots of genome-wide association for hypoxanthine (M21). Note: Y-axis indicates the log_10_(*P*-value). Blue dotted and red solid lines indicate the genome-wide threshold of 0.05 and 0.01 after Bonferroni multiple testing, respectively. The three tracks indicate the metabolites from first sampling time, second sampling time and combined two sampling times, respectively, from outside to inside.

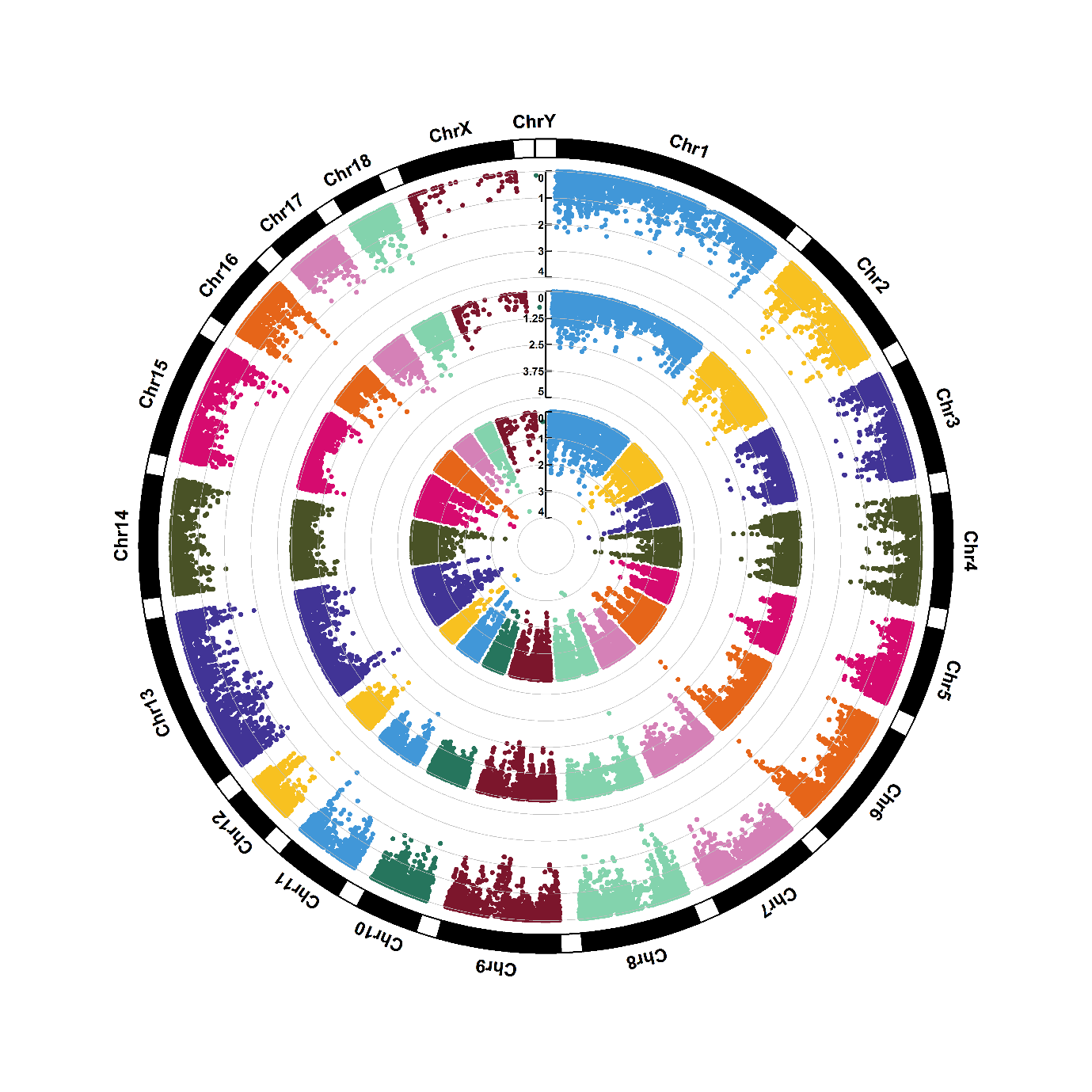

Supplementary figure 1.22. Manhattan plots of genome-wide association for indoleacrylic acid (M22). Note: Y-axis indicates the log_10_(*P*-value). Blue dotted and red solid lines indicate the genome-wide threshold of 0.05 and 0.01 after Bonferroni multiple testing, respectively. The three tracks indicate the metabolites from first sampling time, second sampling time and combined two sampling times, respectively, from outside to inside.

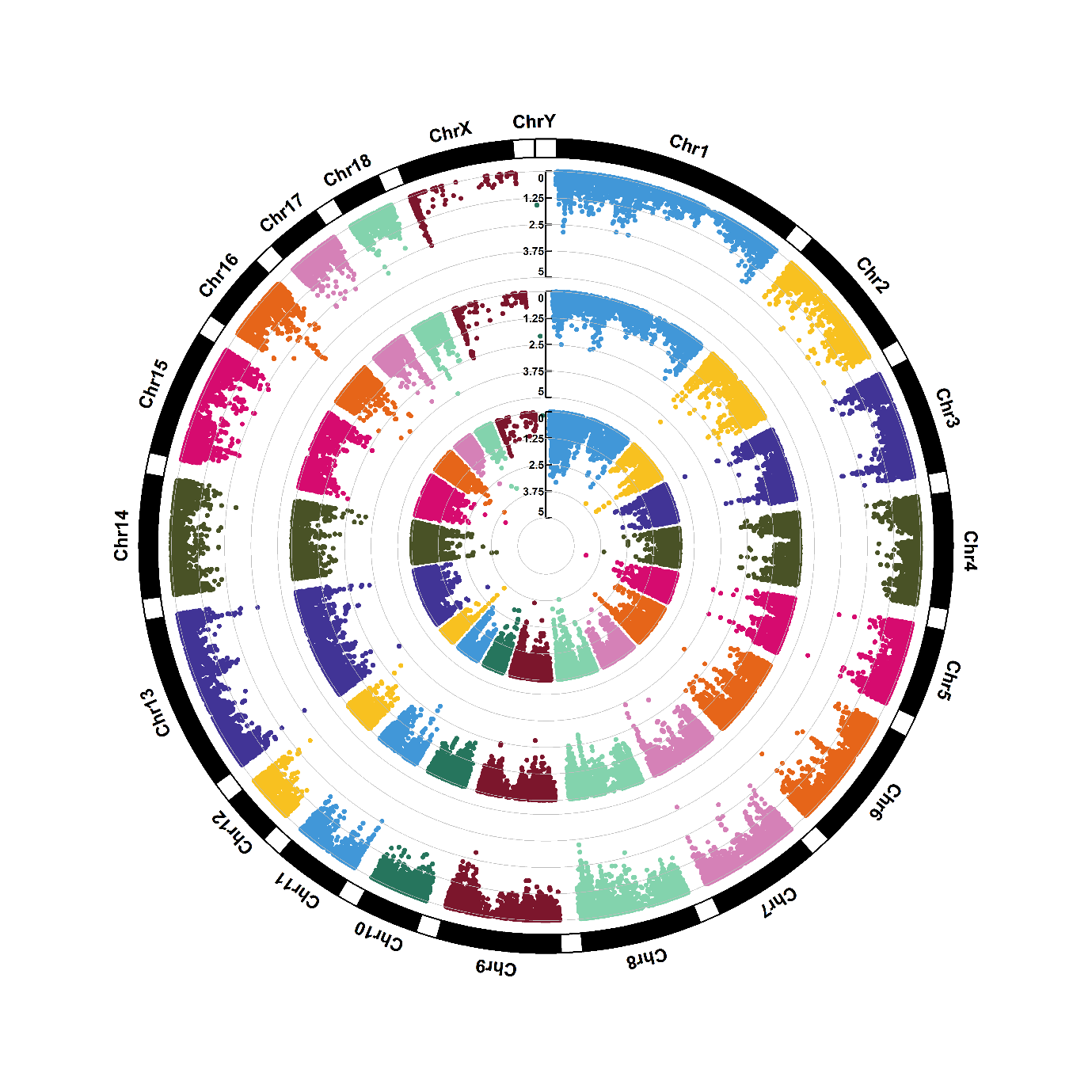

Supplementary figure 1.23. Manhattan plots of genome-wide association for inosine (M23). Note: Y-axis indicates the log_10_(*P*-value). Blue dotted and red solid lines indicate the genome-wide threshold of 0.05 and 0.01 after Bonferroni multiple testing, respectively. The three tracks indicate the metabolites from first sampling time, second sampling time and combined two sampling times, respectively, from outside to inside.

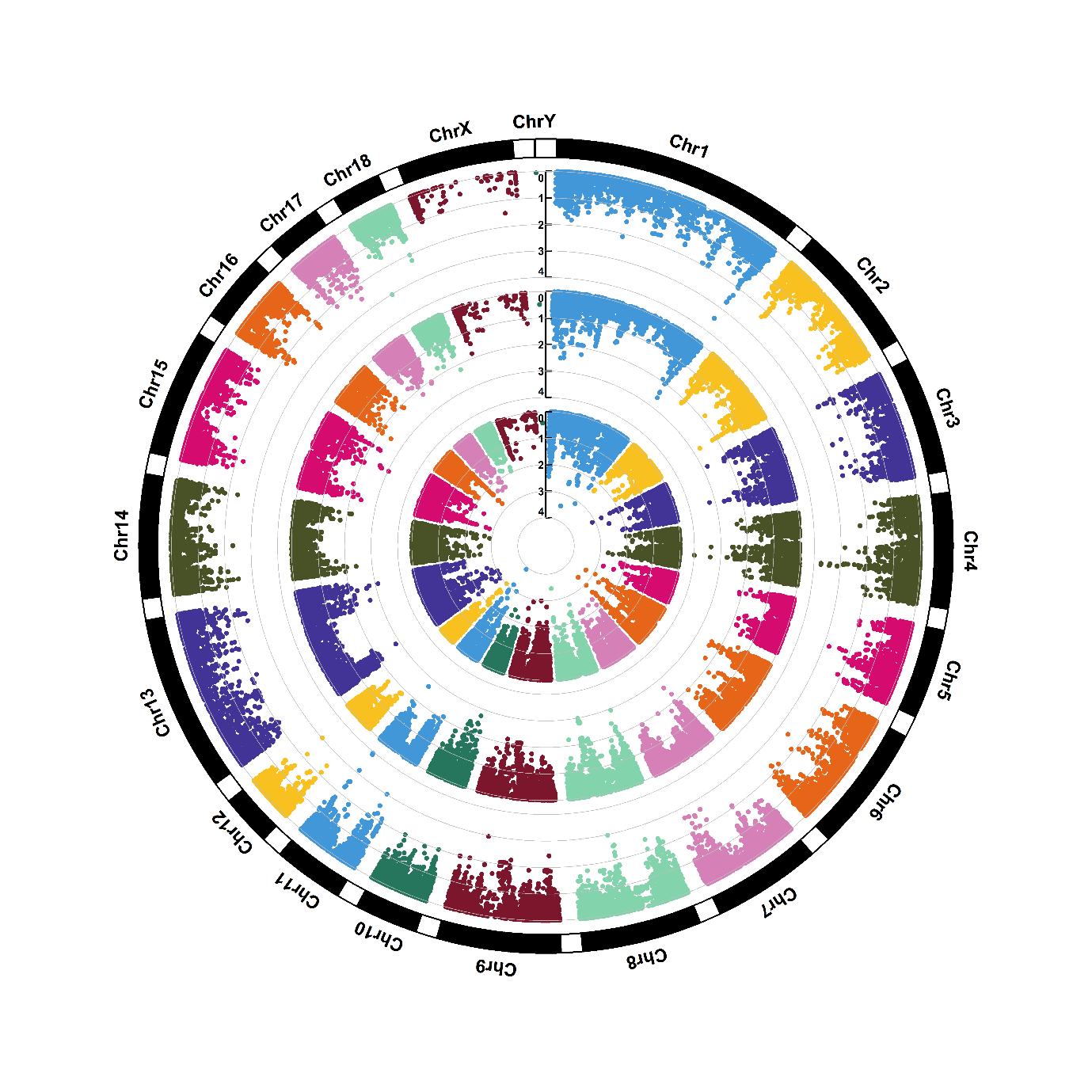

Supplementary figure 1.24. Manhattan plots of genome-wide association for isoleucine (M24). Note: Y-axis indicates the log_10_(*P*-value). Blue dotted and red solid lines indicate the genome-wide threshold of 0.05 and 0.01 after Bonferroni multiple testing, respectively. The three tracks indicate the metabolites from first sampling time, second sampling time and combined two sampling times, respectively, from outside to inside.

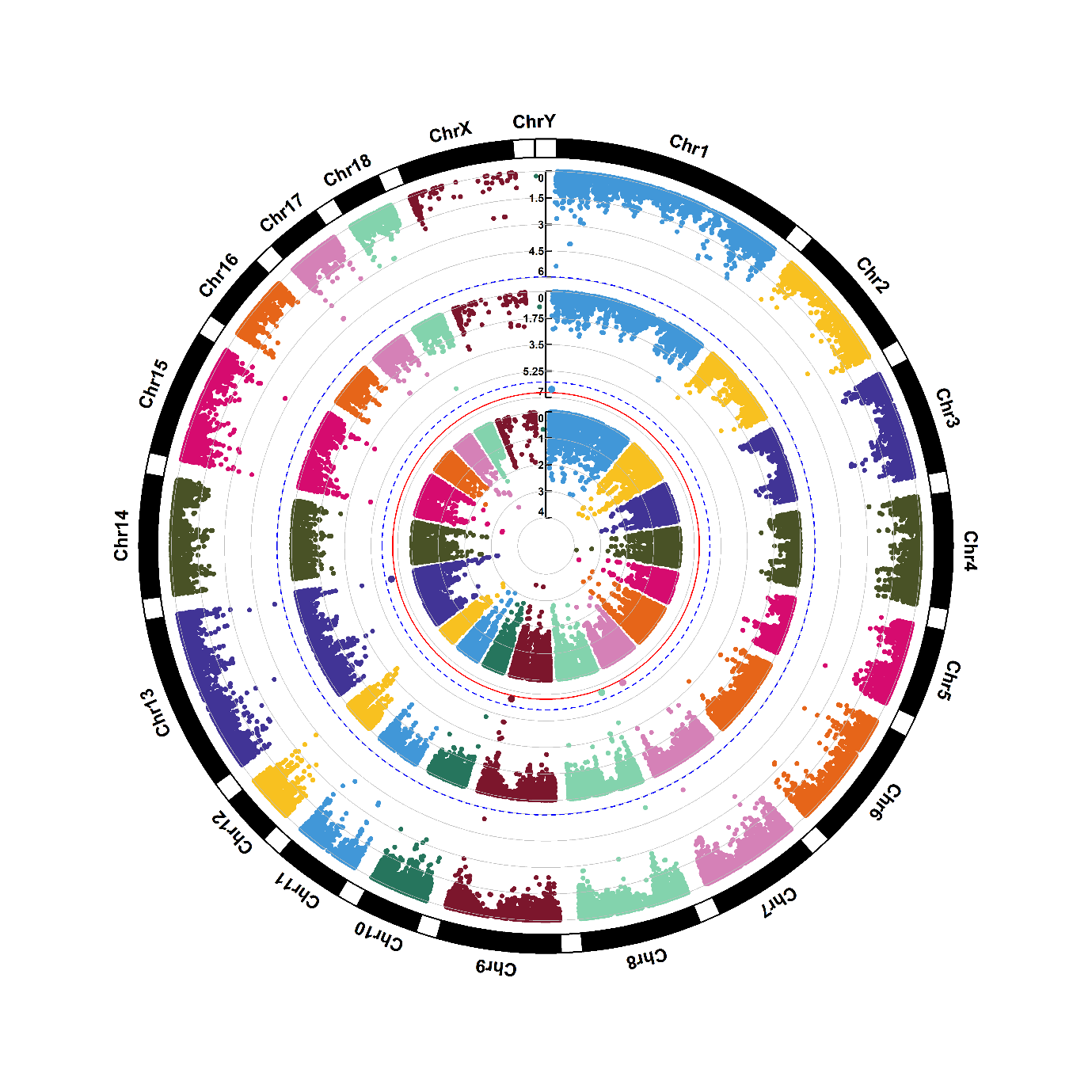

Supplementary figure 1.25. Manhattan plots of genome-wide association for isoleucyl proline (M25). Note: Y-axis indicates the log_10_(*P*-value). Blue dotted and red solid lines indicate the genome-wide threshold of 0.05 and 0.01 after Bonferroni multiple testing, respectively. The three tracks indicate the metabolites from first sampling time, second sampling time and combined two sampling times, respectively, from outside to inside.

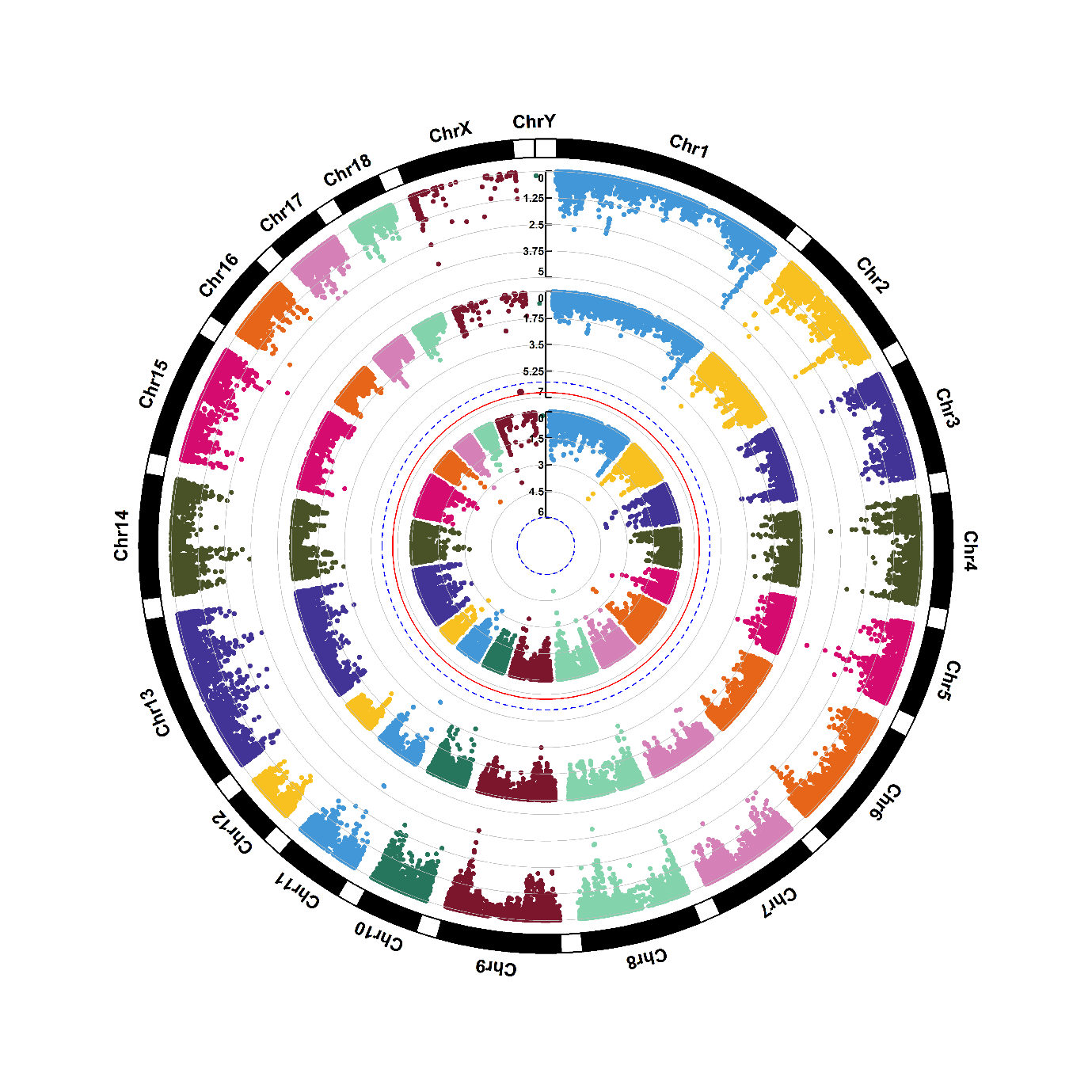

Supplementary figure 1.26. Manhattan plots of genome-wide association for lactic acid (M27). Note: Y-axis indicates the log_10_(*P*-value). Blue dotted and red solid lines indicate the genome-wide threshold of 0.05 and 0.01 after Bonferroni multiple testing, respectively. The three tracks indicate the metabolites from first sampling time, second sampling time and combined two sampling times, respectively, from outside to inside.

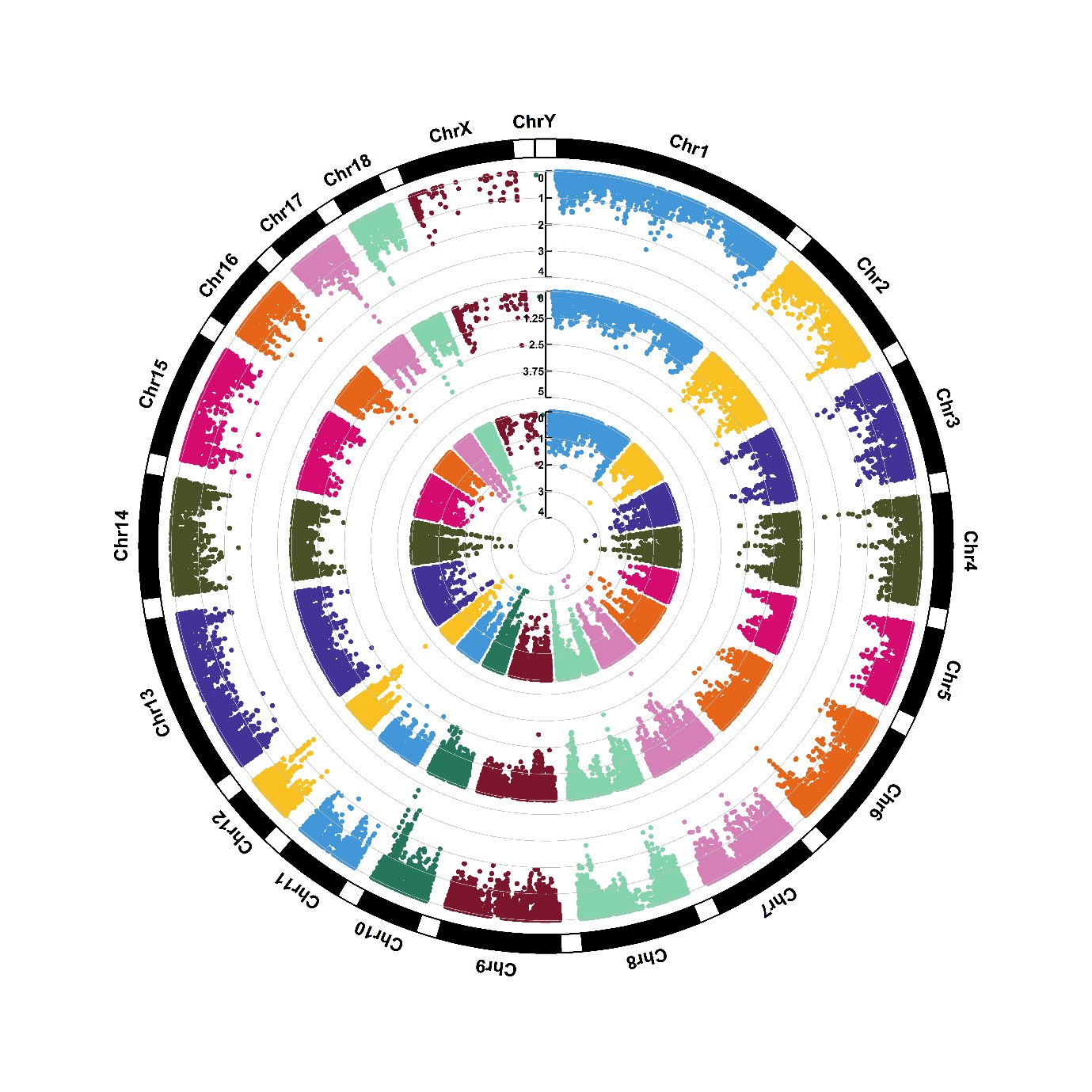

Supplementary figure 1.27. Manhattan plots of genome-wide association for leucyl methionine (M28). Note: Y-axis indicates the log_10_(*P*-value). Blue dotted and red solid lines indicate the genome-wide threshold of 0.05 and 0.01 after Bonferroni multiple testing, respectively. The three tracks indicate the metabolites from first sampling time, second sampling time and combined two sampling times, respectively, from outside to inside.

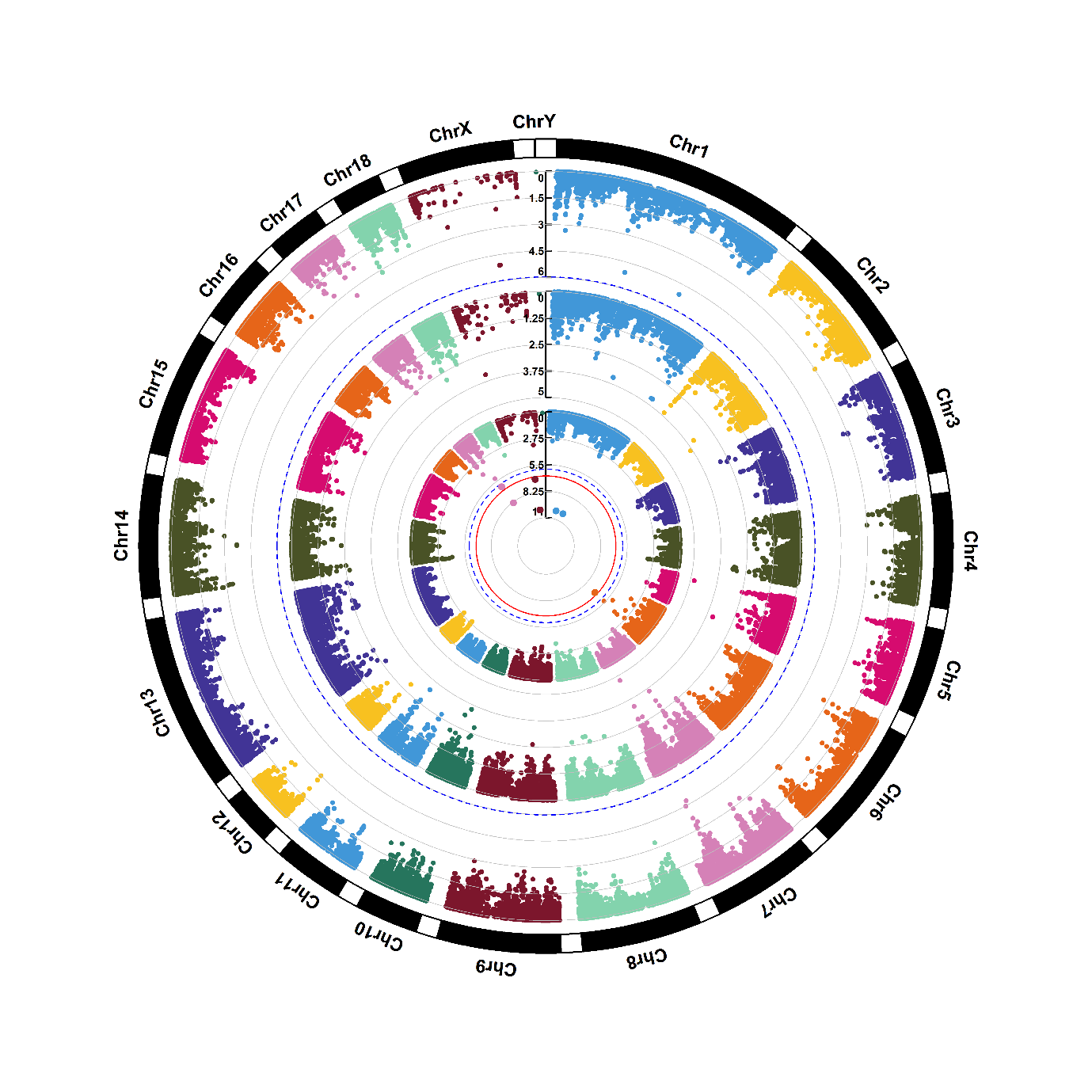

Supplementary figure 1.28. Manhattan plots of genome-wide association for lysoPC(16:0) (M29). Note: Y-axis indicates the log_10_(*P*-value). Blue dotted and red solid lines indicate the genome-wide threshold of 0.05 and 0.01 after Bonferroni multiple testing, respectively. The three tracks indicate the metabolites from first sampling time, second sampling time and combined two sampling times, respectively, from outside to inside.

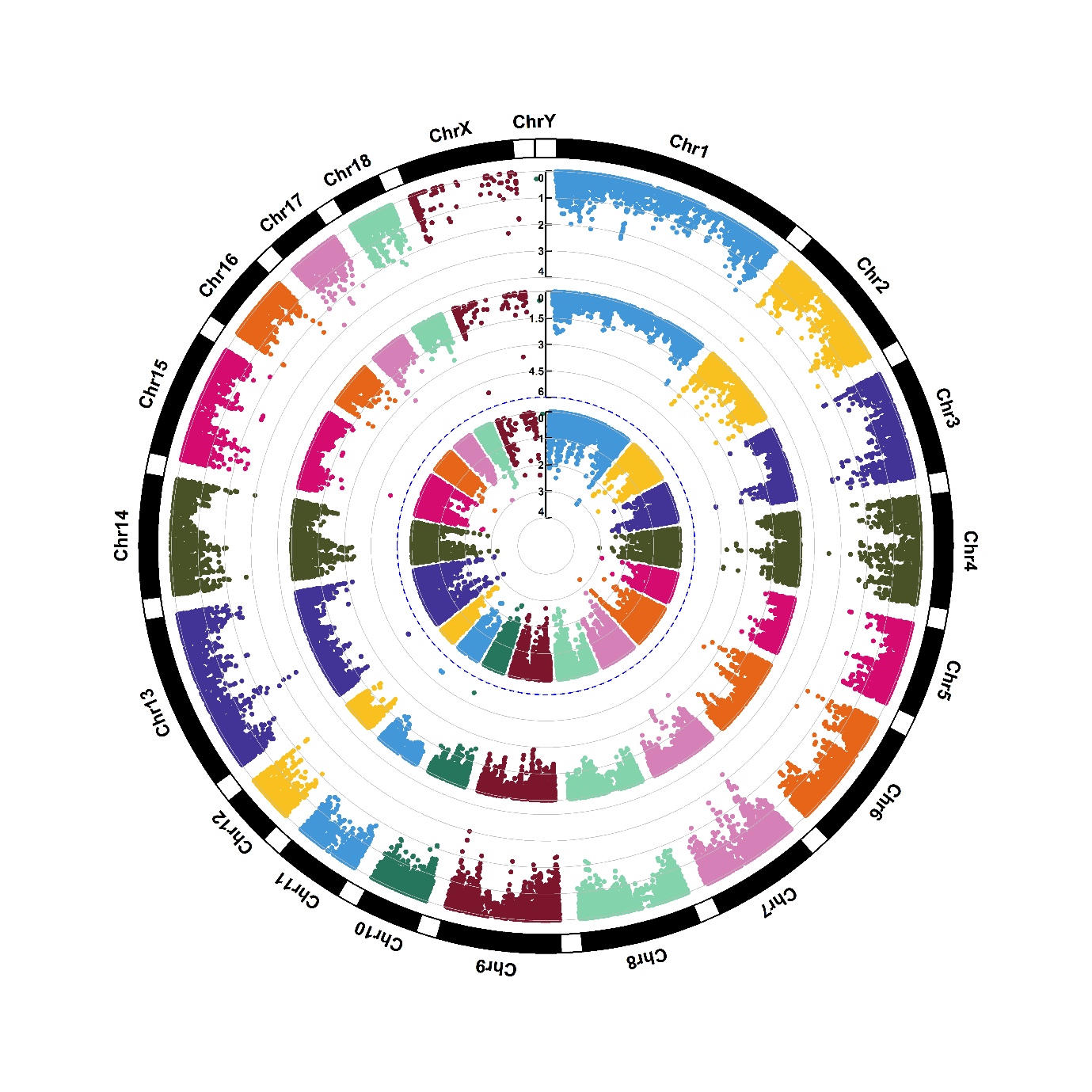

Supplementary figure 1.29. Manhattan plots of genome-wide association for manNAc (M30). Note: Y-axis indicates the log_10_(*P*-value). Blue dotted and red solid lines indicate the genome-wide threshold of 0.05 and 0.01 after Bonferroni multiple testing, respectively. The three tracks indicate the metabolites from first sampling time, second sampling time and combined two sampling times, respectively, from outside to inside.

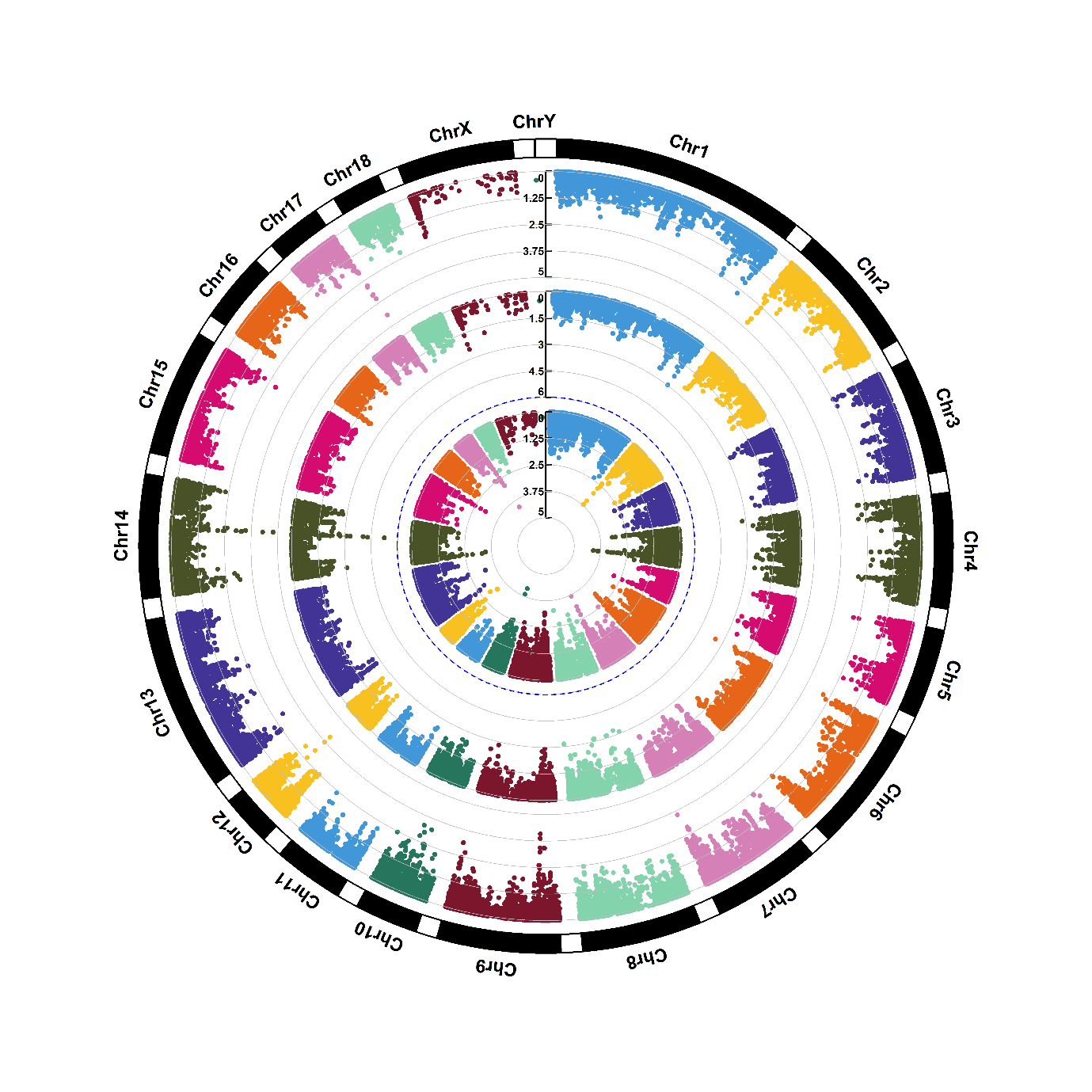

Supplementary figure 1.30. Manhattan plots of genome-wide association for methionine (M31). Note: Y-axis indicates the log_10_(*P*-value). Blue dotted and red solid lines indicate the genome-wide threshold of 0.05 and 0.01 after Bonferroni multiple testing, respectively. The three tracks indicate the metabolites from first sampling time, second sampling time and combined two sampling times, respectively, from outside to inside.

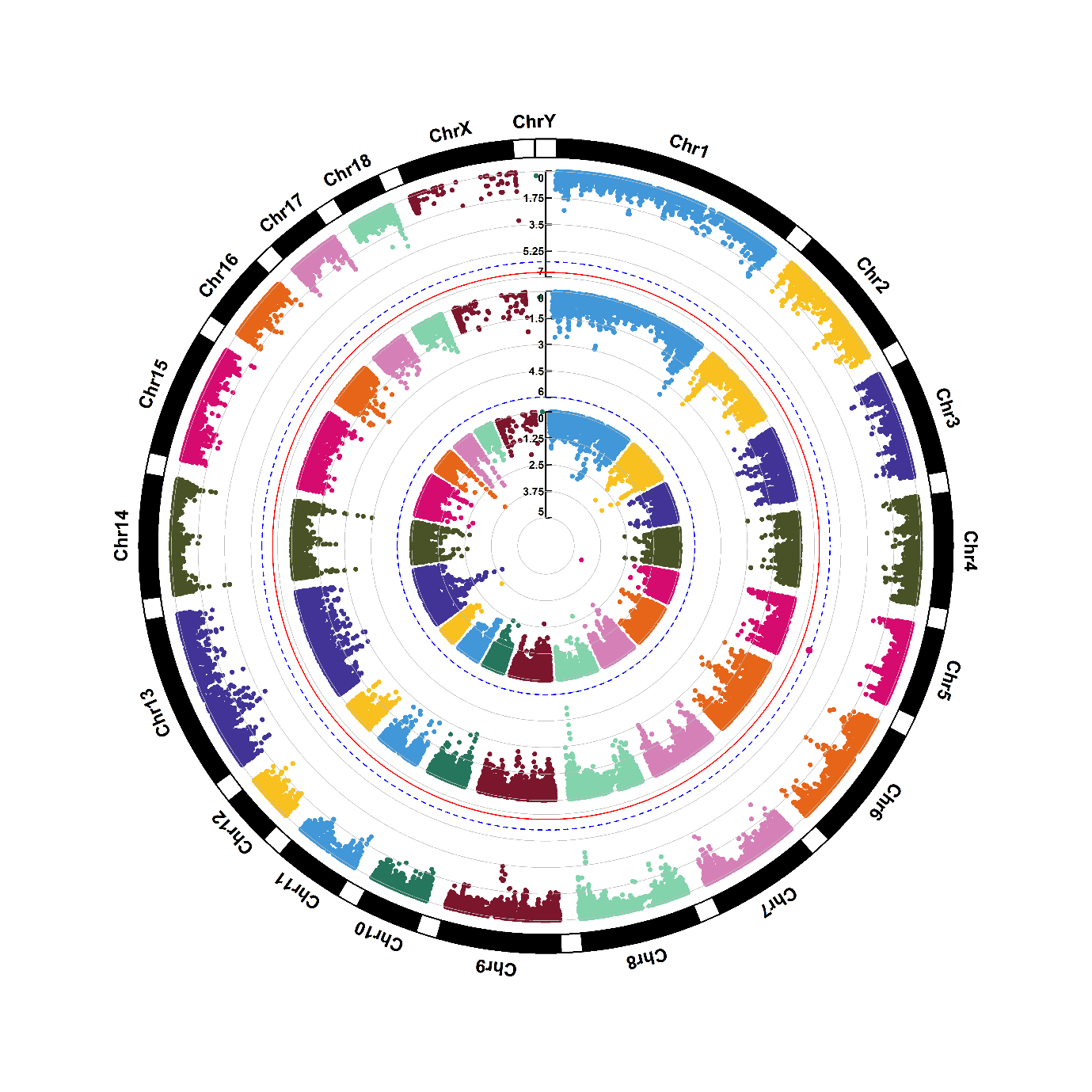

Supplementary figure 1.31. Manhattan plots of genome-wide association for monosaccharide (M32). Note: Y-axis indicates the log_10_(*P*-value). Blue dotted and red solid lines indicate the genome-wide threshold of 0.05 and 0.01 after Bonferroni multiple testing, respectively. The three tracks indicate the metabolites from first sampling time, second sampling time and combined two sampling times, respectively, from outside to inside.

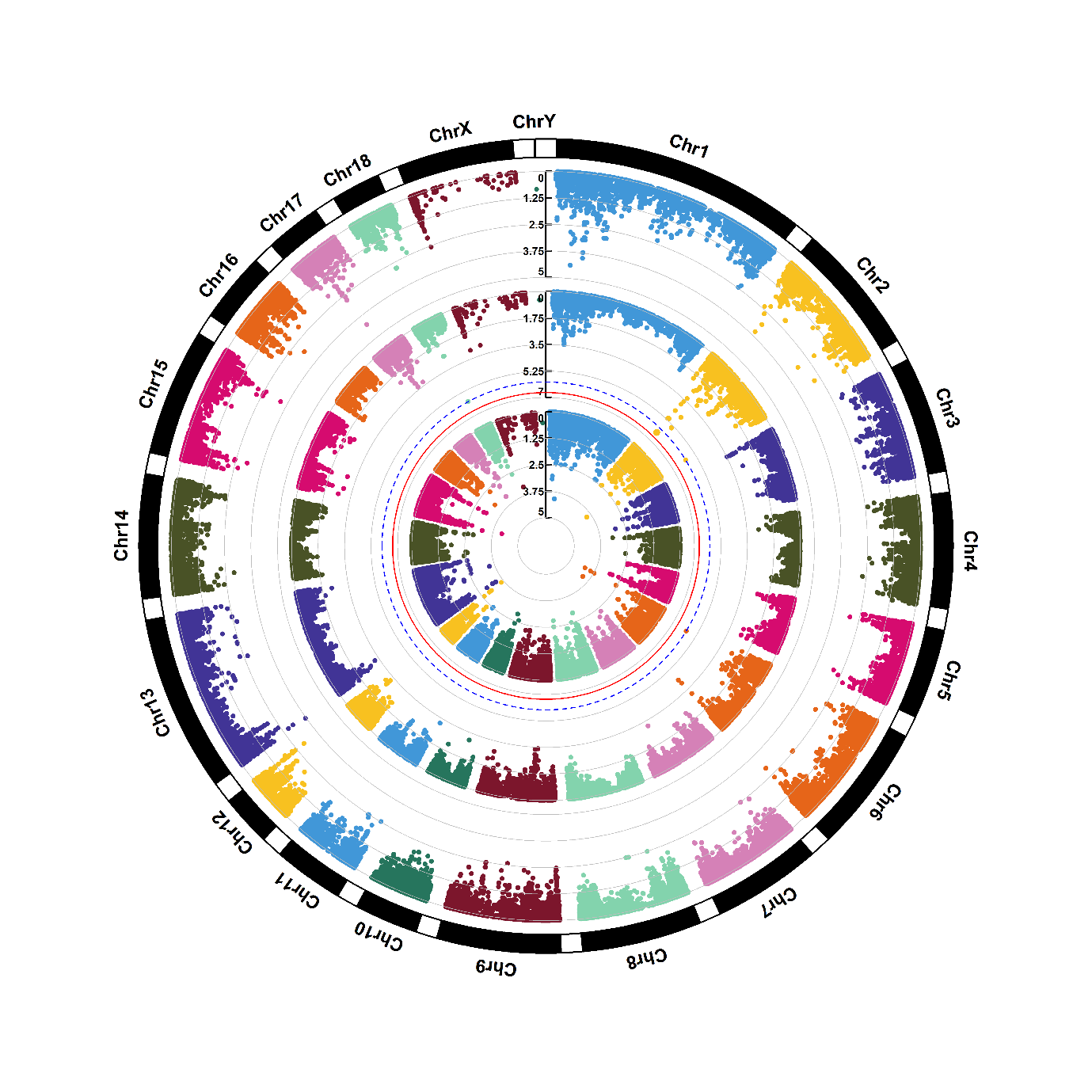

Supplementary figure 1.32. Manhattan plots of genome-wide association for nicotine amide (M33). Note: Y-axis indicates the log_10_(*P*-value). Blue dotted and red solid lines indicate the genome-wide threshold of 0.05 and 0.01 after Bonferroni multiple testing, respectively. The three tracks indicate the metabolites from first sampling time, second sampling time and combined two sampling times, respectively, from outside to inside.

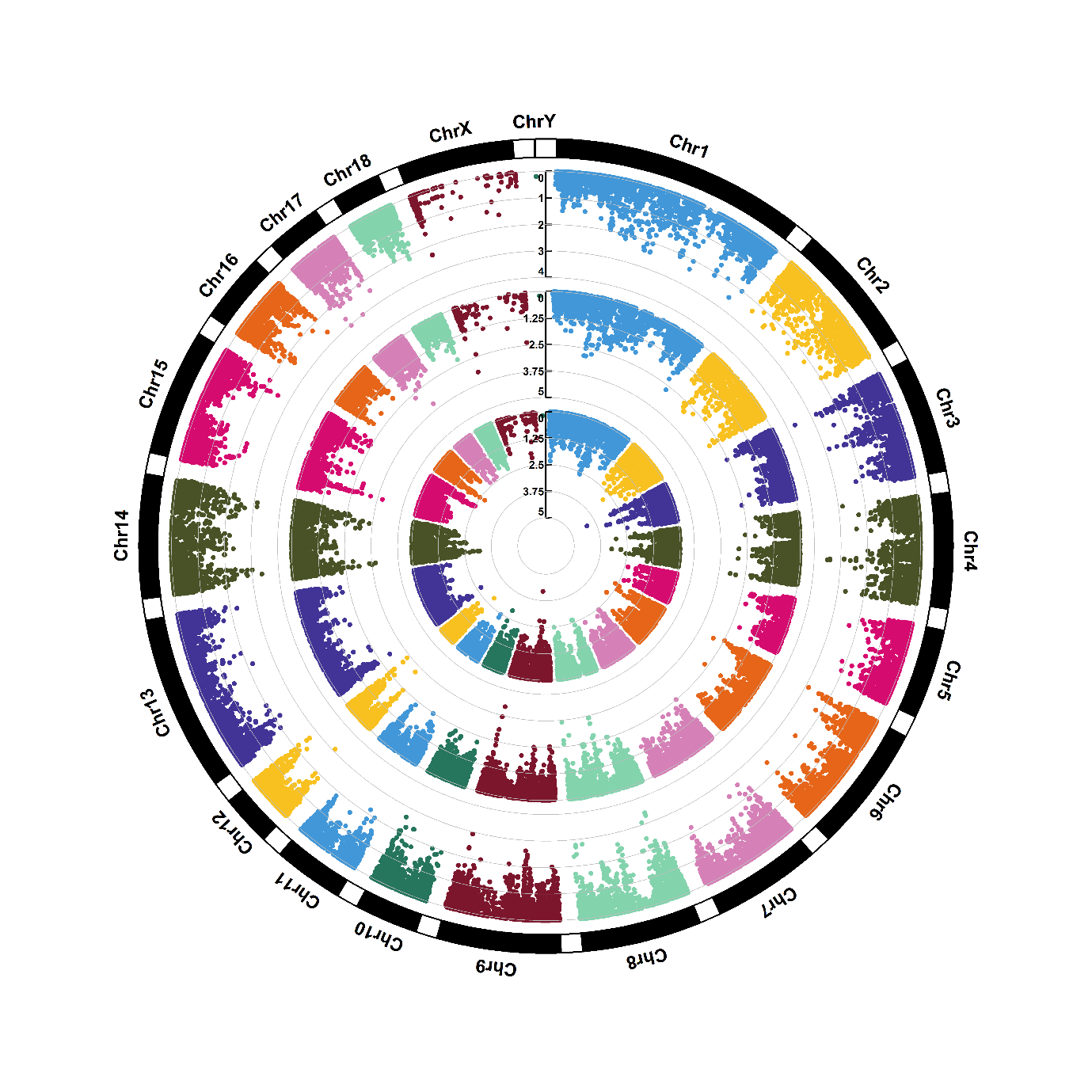

Supplementary figure 1.33. Manhattan plots of genome-wide association for ornithine (M34). Note: Y-axis indicates the log_10_(*P*-value). Blue dotted and red solid lines indicate the genome-wide threshold of 0.05 and 0.01 after Bonferroni multiple testing, respectively. The three tracks indicate the metabolites from first sampling time, second sampling time and combined two sampling times, respectively, from outside to inside.

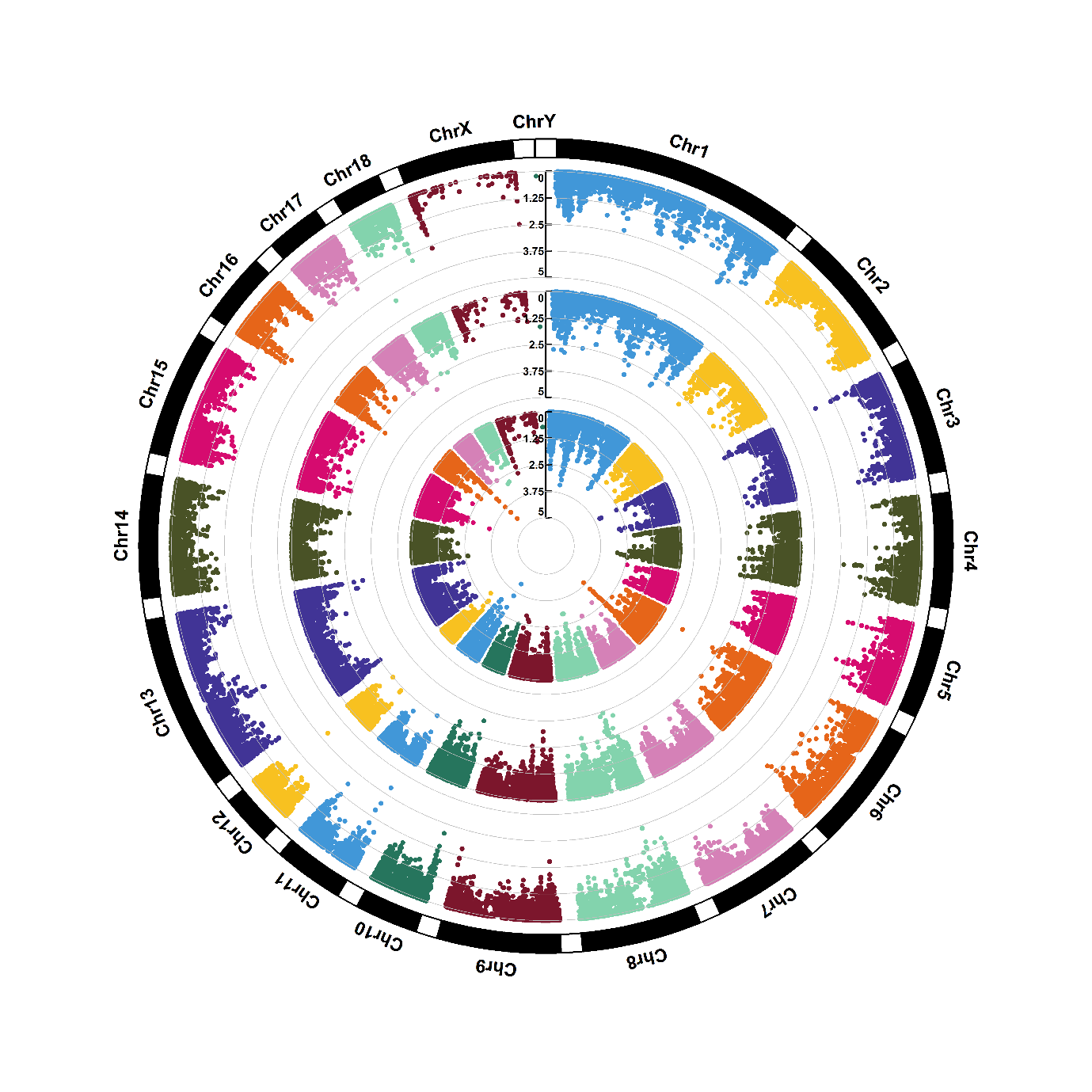

Supplementary figure 1.34. Manhattan plots of genome-wide association for phenylalanine (M35). Note: Y-axis indicates the log_10_(*P*-value). Blue dotted and red solid lines indicate the genome-wide threshold of 0.05 and 0.01 after Bonferroni multiple testing, respectively. The three tracks indicate the metabolites from first sampling time, second sampling time and combined two sampling times, respectively, from outside to inside.

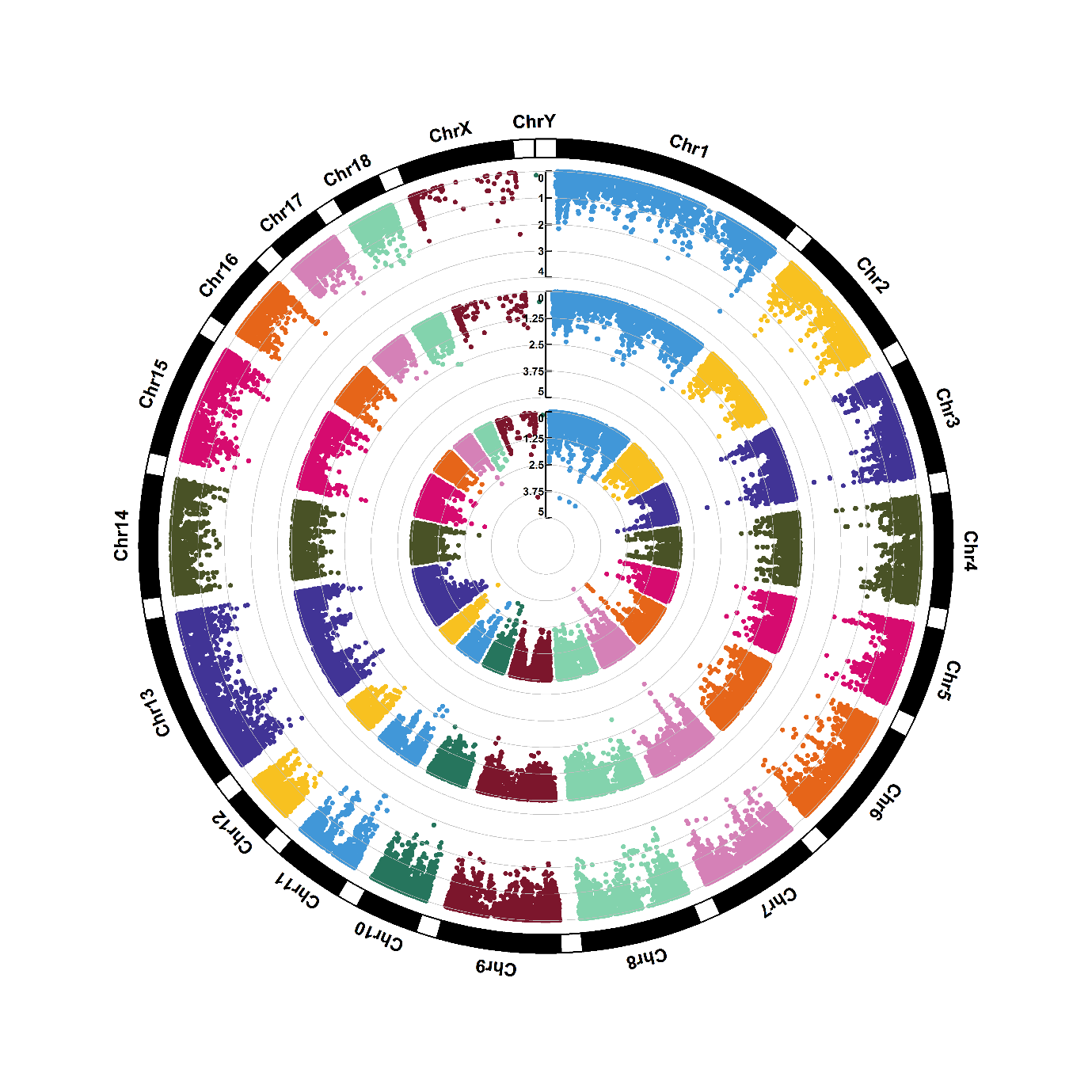

Supplementary figure 1.35. Manhattan plots of genome-wide association for proline (M36). Note: Y-axis indicates the log_10_(*P*-value). Blue dotted and red solid lines indicate the genome-wide threshold of 0.05 and 0.01 after Bonferroni multiple testing, respectively. The three tracks indicate the metabolites from first sampling time, second sampling time and combined two sampling times, respectively, from outside to inside.

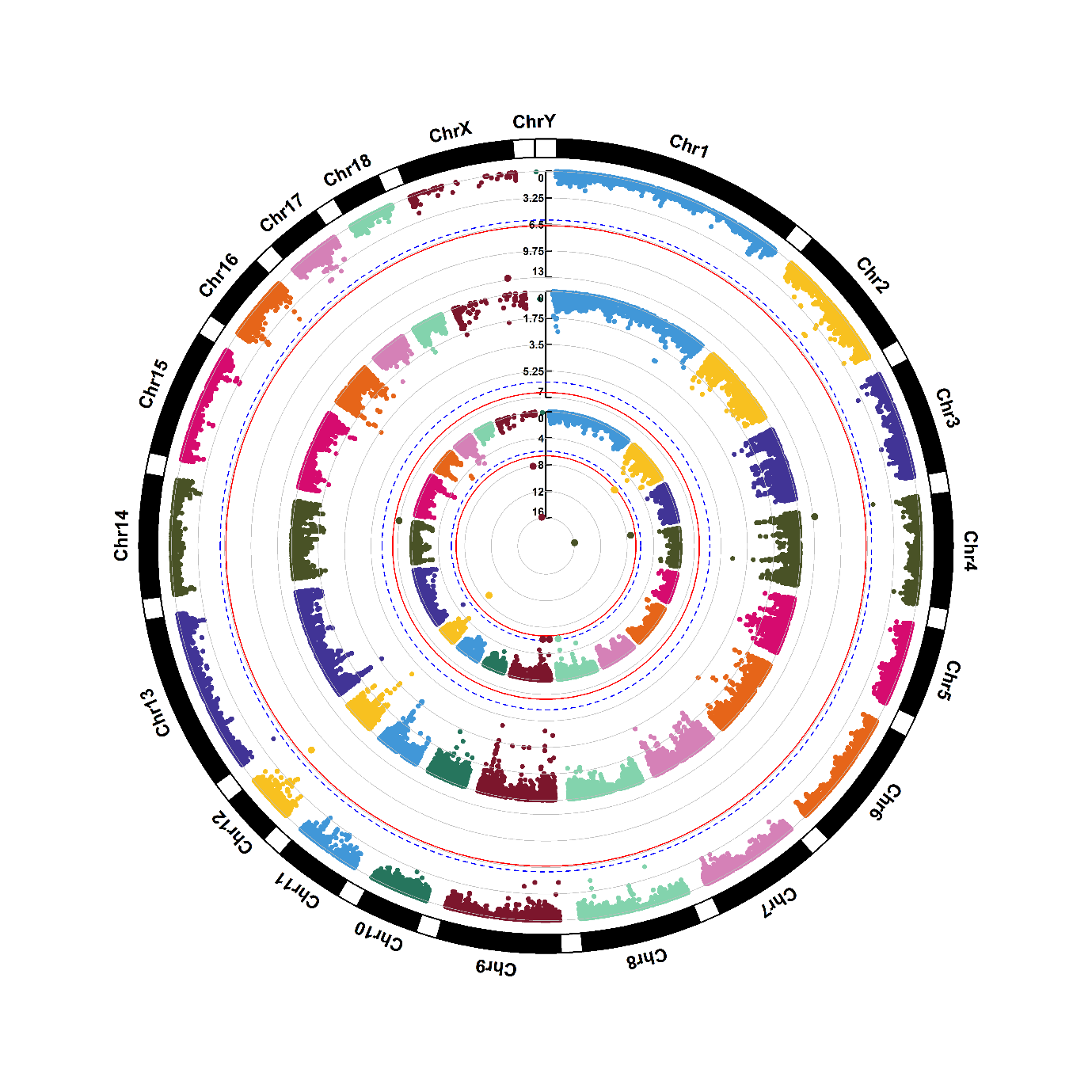

Supplementary figure 1.36. Manhattan plots of genome-wide association for pyruvic acid (M38). Note: Y-axis indicates the log_10_(*P*-value). Blue dotted and red solid lines indicate the genome-wide threshold of 0.05 and 0.01 after Bonferroni multiple testing, respectively. The three tracks indicate the metabolites from first sampling time, second sampling time and combined two sampling times, respectively, from outside to inside.

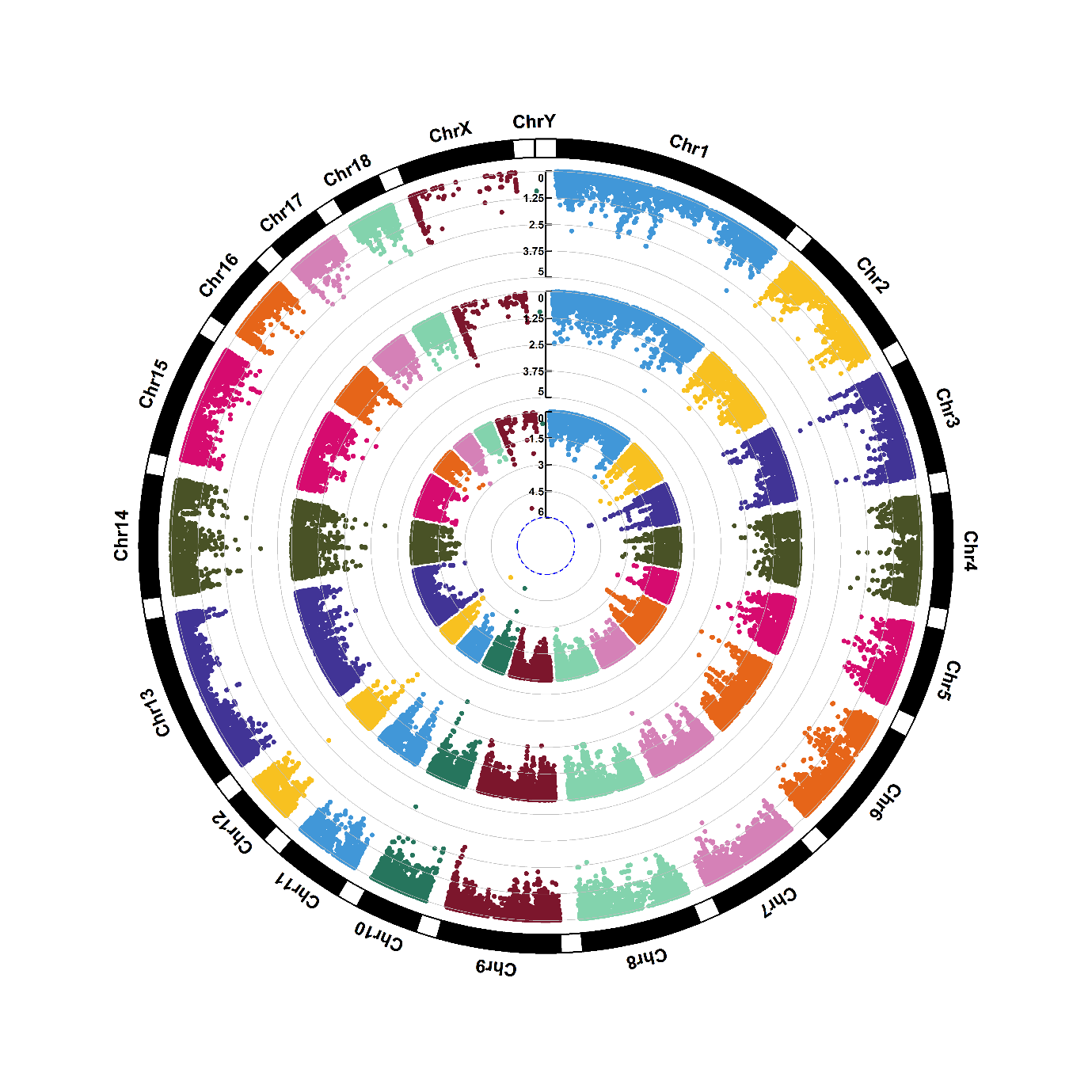

Supplementary figure 1.37. Manhattan plots of genome-wide association for riboflavin (M39). Note: Y-axis indicates the log_10_(*P*-value). Blue dotted and red solid lines indicate the genome-wide threshold of 0.05 and 0.01 after Bonferroni multiple testing, respectively. The three tracks indicate the metabolites from first sampling time, second sampling time and combined two sampling times, respectively, from outside to inside.

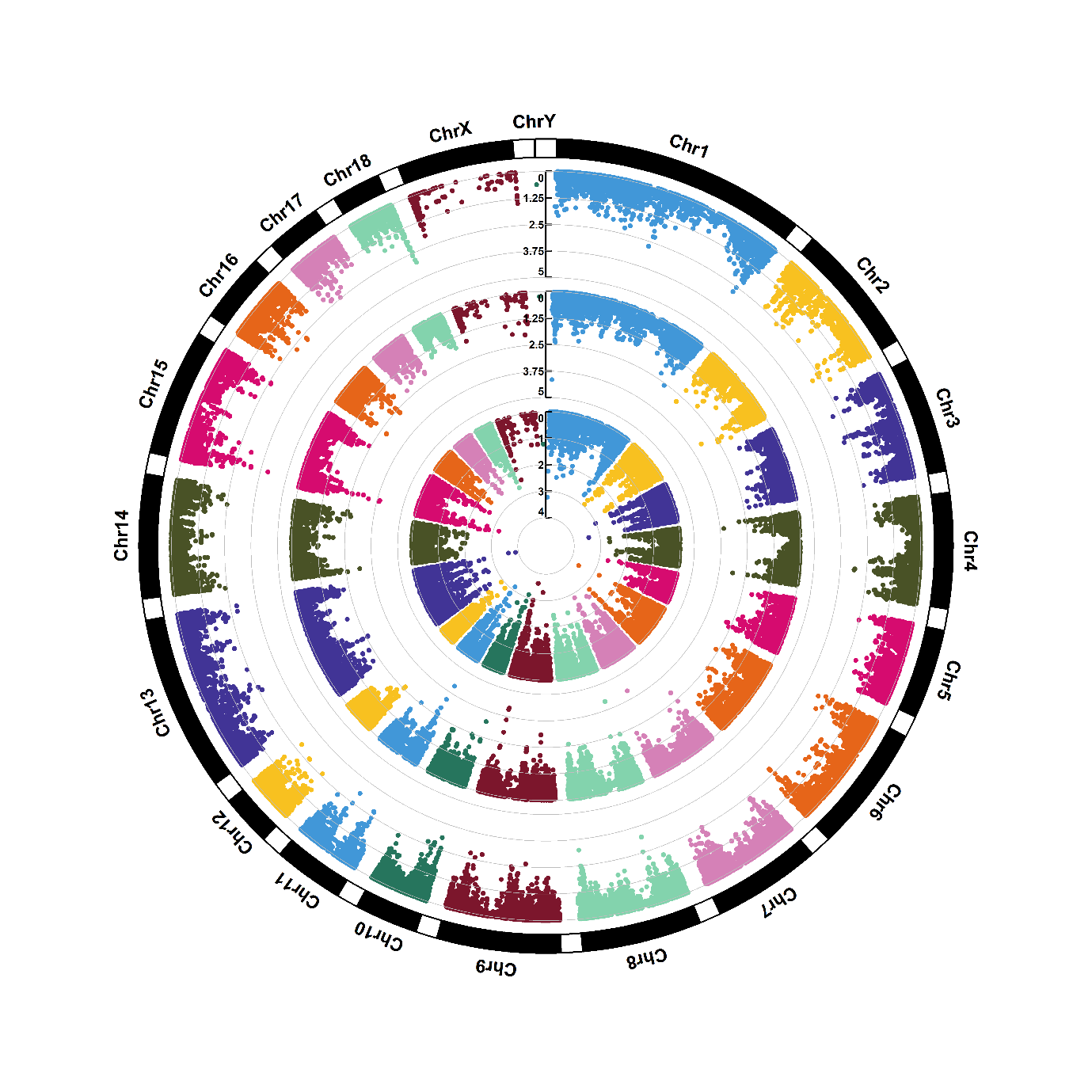

Supplementary figure 1.38. Manhattan plots of genome-wide association for sorbitol (M40). Note: Y-axis indicates the log_10_(*P*-value). Blue dotted and red solid lines indicate the genome-wide threshold of 0.05 and 0.01 after Bonferroni multiple testing, respectively. The three tracks indicate the metabolites from first sampling time, second sampling time and combined two sampling times, respectively, from outside to inside.

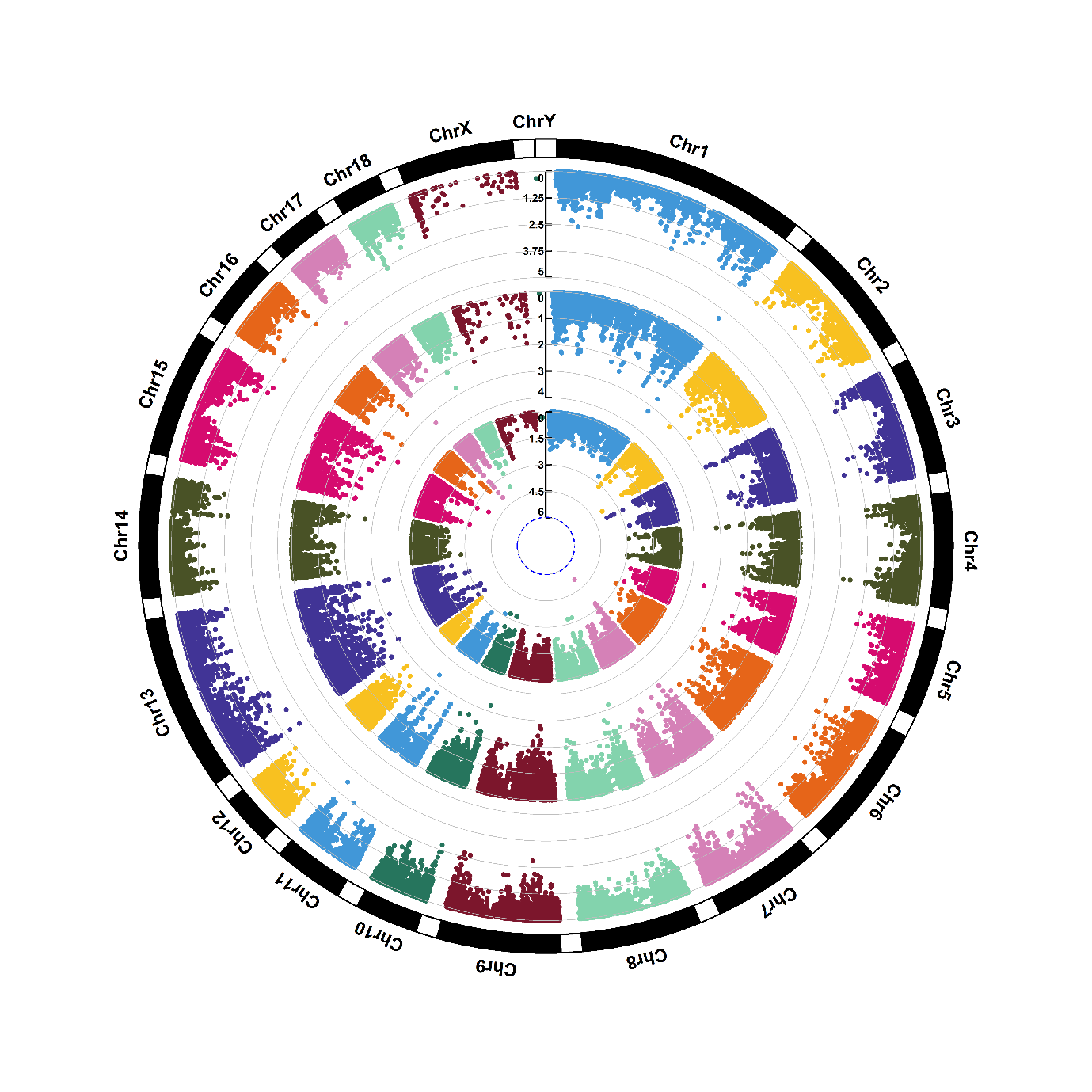

Supplementary figure 1.39. Manhattan plots of genome-wide association for thiamine (M41). Note: Y-axis indicates the log_10_(*P*-value). Blue dotted and red solid lines indicate the genome-wide threshold of 0.05 and 0.01 after Bonferroni multiple testing, respectively. The three tracks indicate the metabolites from first sampling time, second sampling time and combined two sampling times, respectively, from outside to inside.

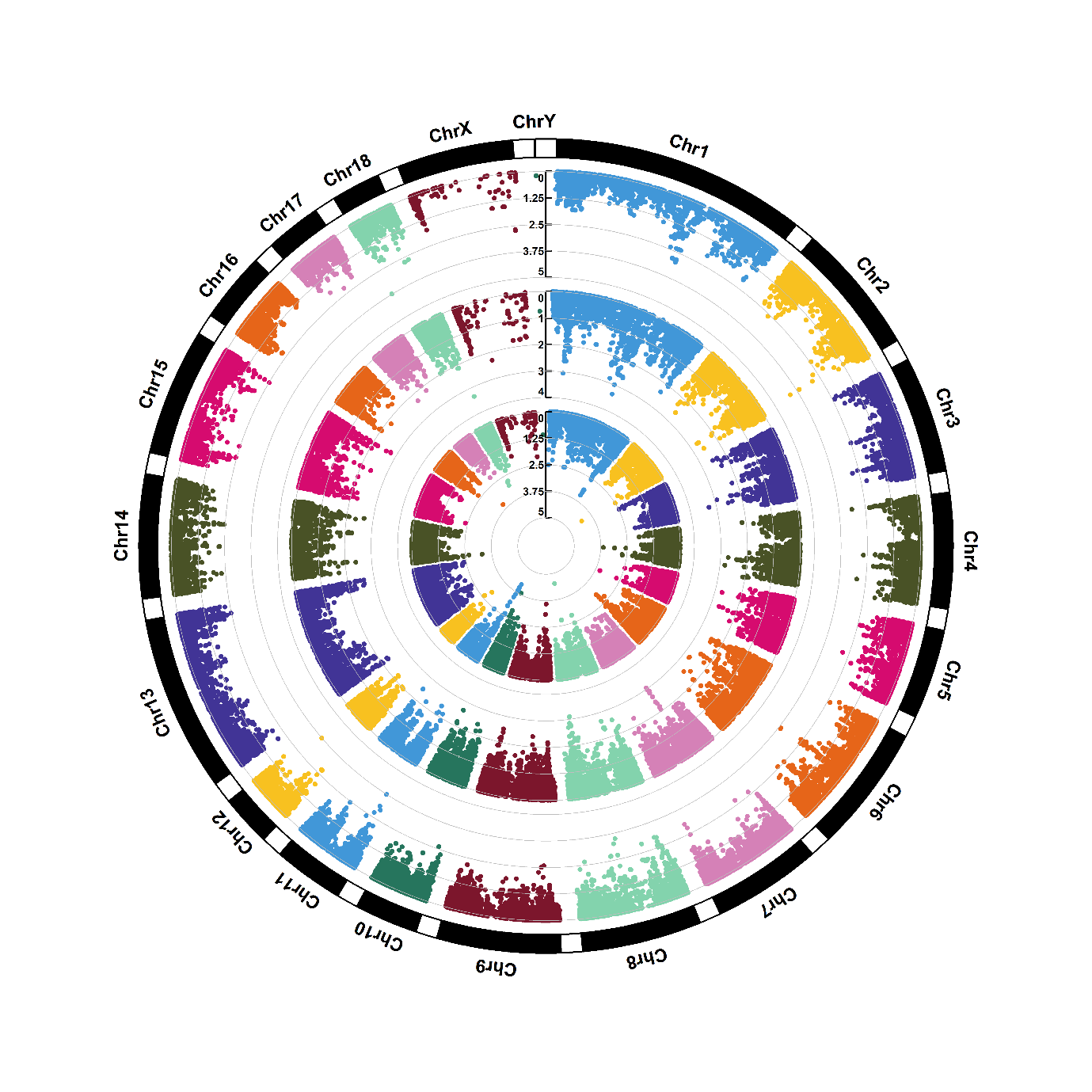

Supplementary figure 1.40. Manhattan plots of genome-wide association for threonine (M42). Note: Y-axis indicates the log_10_(*P*-value). Blue dotted and red solid lines indicate the genome-wide threshold of 0.05 and 0.01 after Bonferroni multiple testing, respectively. The three tracks indicate the metabolites from first sampling time, second sampling time and combined two sampling times, respectively, from outside to inside.

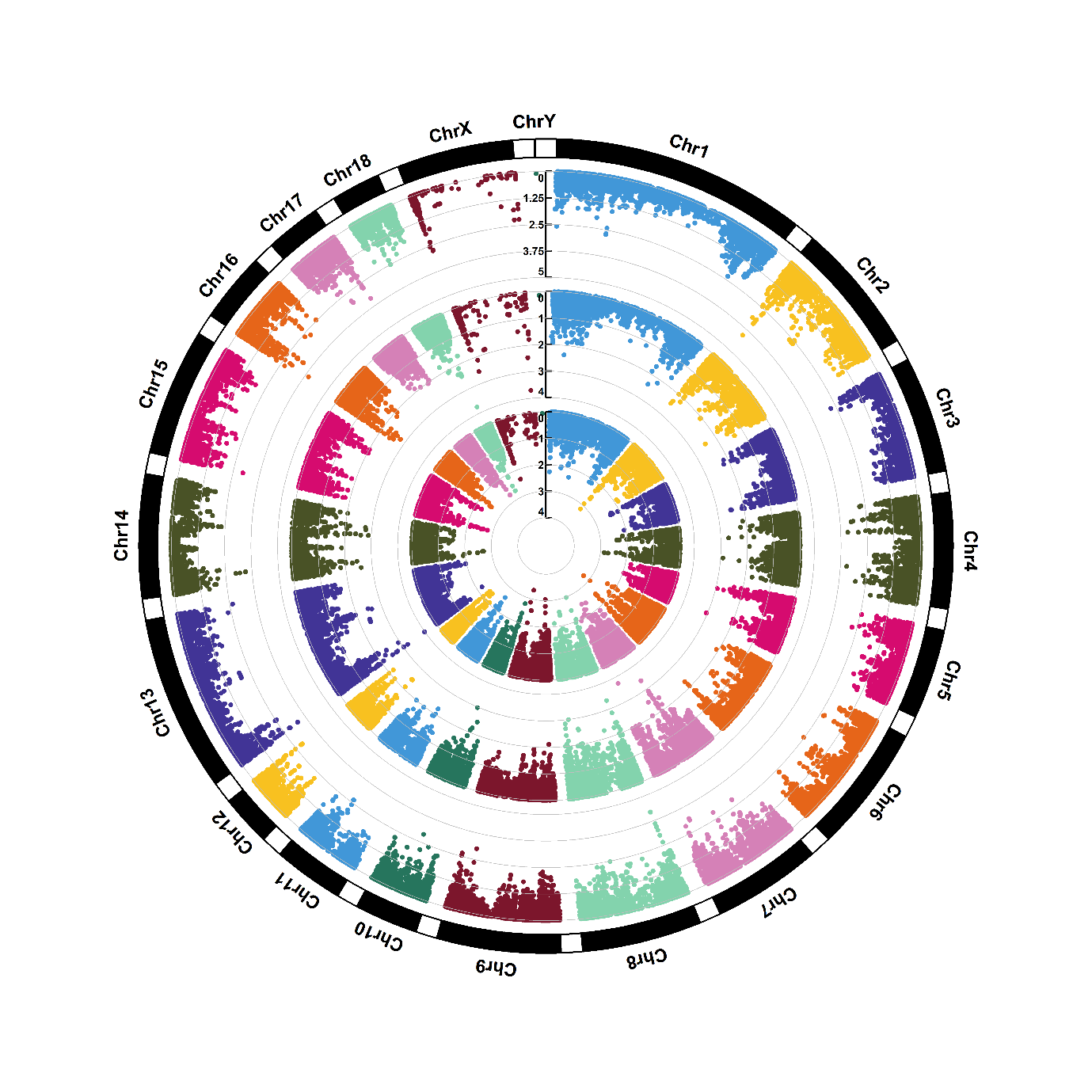

Supplementary figure 1.41. Manhattan plots of genome-wide association for thymidine (M43). Note: Y-axis indicates the log_10_(*P*-value). Blue dotted and red solid lines indicate the genome-wide threshold of 0.05 and 0.01 after Bonferroni multiple testing, respectively. The three tracks indicate the metabolites from first sampling time, second sampling time and combined two sampling times, respectively, from outside to inside.

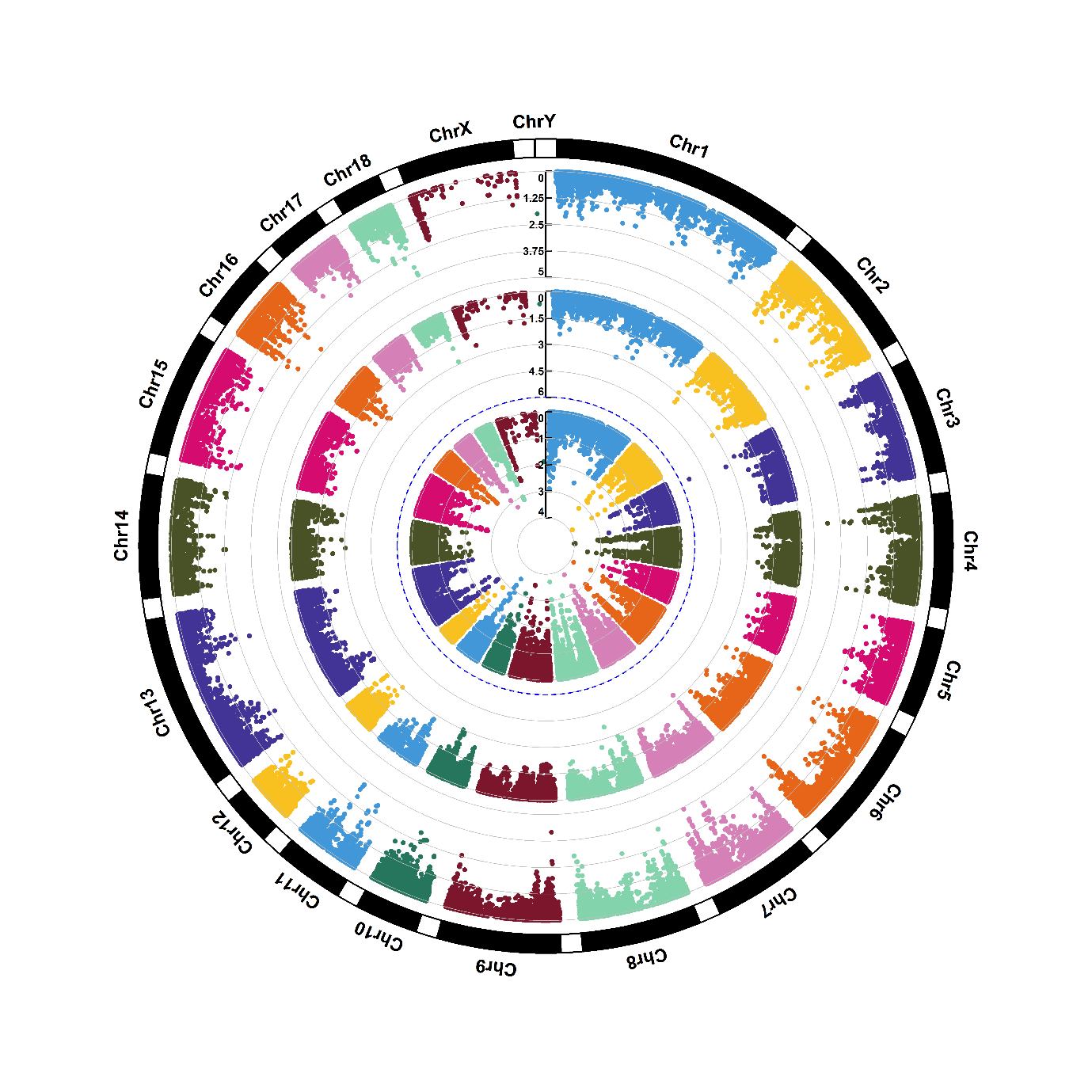

Supplementary figure 1.42. Manhattan plots of genome-wide association for uridine (M44). Note: Y-axis indicates the log_10_(*P*-value). Blue dotted and red solid lines indicate the genome-wide threshold of 0.05 and 0.01 after Bonferroni multiple testing, respectively. The three tracks indicate the metabolites from first sampling time, second sampling time and combined two sampling times, respectively, from outside to inside.

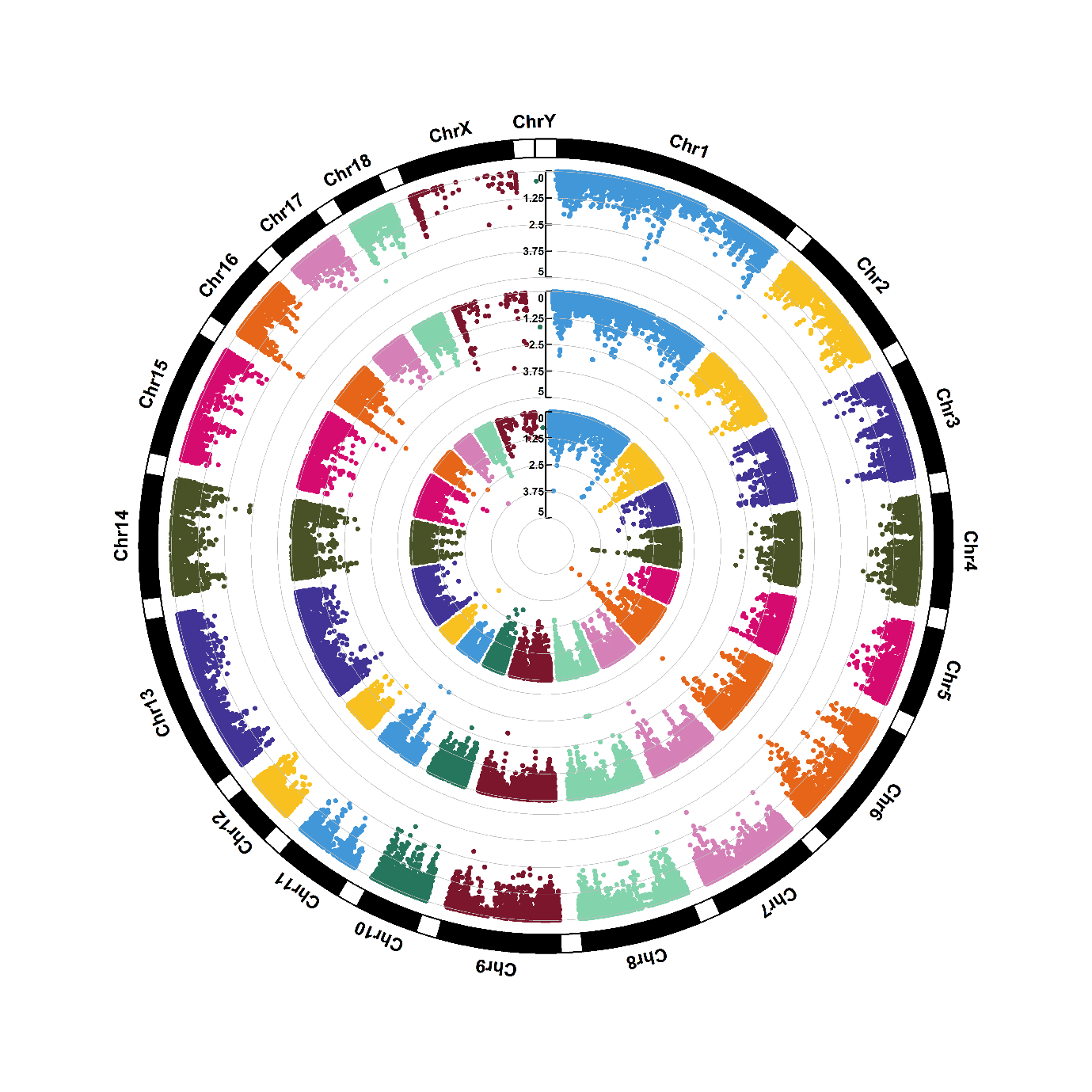

Supplementary figure 1.43. Manhattan plots of genome-wide association for xanthine (M45). Note: Y-axis indicates the log_10_(*P*-value). Blue dotted and red solid lines indicate the genome-wide threshold of 0.05 and 0.01 after Bonferroni multiple testing, respectively. The three tracks indicate the metabolites from first sampling time, second sampling time and combined two sampling times, respectively, from outside to inside.

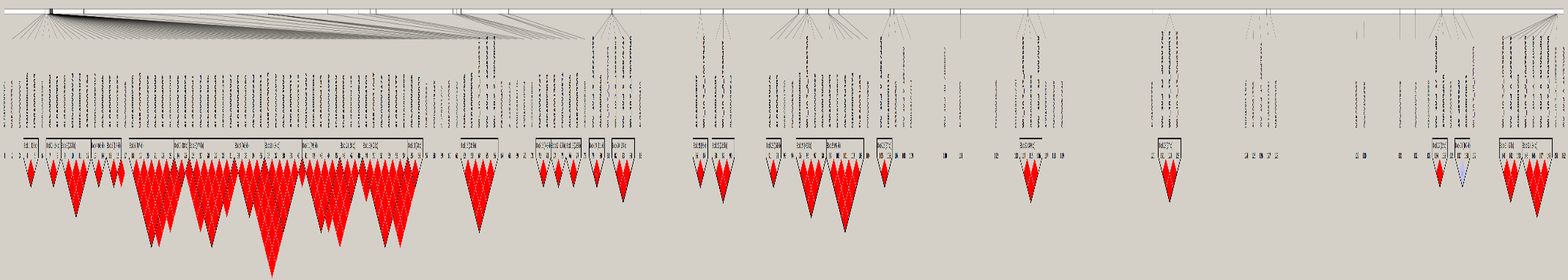

Supplementary figure 2. Linkage disequilibrium (LD) pattern for all significant SNPs. Note: Solid line triangle indicates LD. One square indicates LD level (r^2^) between two SNPs and the squares are colored by D’/LLOR standard scheme. D’/LLOR standard scheme is that red indicates LLOR > 2, D’ = 1; pink indicates LLOR > 2, D’ < 1; blue indicates LLOR < 2, D’ = 1; white indicates LLOR < 2, D’ < 1. LLOR is the logarithm of likelihood odds ratio and the reliable index to measure D’.
